# Unveiling a missing component of the atypical type IV secretion system required for natural transformation of *Helicobacter pylori*

**DOI:** 10.64898/2026.04.01.715814

**Authors:** J. Francisco Villa, Sumedha Kondekar, Yoann Fauconnet, Mérick Machouri, Céline Lacrouts, Xavier Veaute, Raphaël Guérois, Eduardo P. C. Rocha, Jessica Andreani, J. Pablo Radicella

## Abstract

Exchange of genetic information by natural transformation shapes bacterial evolution. In *Helicobacter pylori* it is thought to drive its unusually high recombination rate, which has a crucial role in the evolution of virulence and the propagation of antibiotics resistance genes. While in most cases uptake of the incoming DNA into the periplasm is mediated by type IV pili, in *H. pylori* this initial step of natural transformation requires ComB, a unique competence-specific type IV secretion system (T4SS). The mechanisms by which ComB mediates DNA uptake are still poorly understood, since T4SS are usually involved in an opposite process of DNA export. Here, we identify a gene (*hp1421*) that is absolutely required for uptake of the transforming DNA into the periplasm, although distant from the *comB* operons. We show that *hp1421* codes for a hexameric ATPase from the VirB11 family. HP1421 is present in the cytoplasm and interacts with ComB4, another ATPase of the T4SS inner membrane subcomplex. The structural modelling and functional analysis of HP1421 and its interaction with ComB4 indicate that HP1421 is a missing component of the ComB inner-membrane subcomplex that we propose to name ComB11. Phylogenetic analyses show that *comB11* is a *H. pylori* core gene and suggest that the competence-dedicated ComB T4SS was a recent acquisition within *Helicobacteraceae.* Hence, co-option of the T4SS for DNA transformation requires nearly all the proteins that were previously essential for DNA conjugation.

**Author Summary:** The capacity of bacteria to exchange genetic information contributes in the case of pathogens to the spreading of antibiotic resistance and virulence factors. For *Helicobacter pylori,* a Gram-negative pathogen that colonises about half of the world population and is at the origin of diseases such as ulcers and gastric cancers, natural transformation is the major mechanism of horizontal gene transfer. However, *H. pylori* uses a very unusual system to capture and internalise the foreign DNA. Indeed, a Type 4 secretion system mediates this process. Here, we identify a so far missing and essential component of the T4SS, coded by a gene distant from the operon coding the other subunits. Through a combination of structural modelling, biochemical and microscopy approaches we show that this ATPase is an indispensable part of the ComB T4SS. Our study provides new insights into the mechanism by which the peculiar ComB T4SS works backwards to allow the passage of the tDNA from the bacterial environment into the periplasm.

## Introduction

Horizontal gene transfer strongly contributes to the adaptive potential of bacterial populations by accelerating evolutionary rates through the spread of non-selected genetic variants[1–4]. In pathogens, virulence and antibiotic resistance are often associated with horizontal gene transfer. This has been well documented in the case of *Helicobacter pylori,* where high spontaneous mutation rates combined with extensive horizontal gene transfer underlie the amazing genome plasticity of the species that contributes to its success as a pathogen [5,6].

Natural transformation (NT) is one of the three main pathways of horizontal gene transfer. During NT, free DNA present in the environment is captured and integrated into the bacterial genome. This mechanism is considered to be the main pathway for horizontal genetic exchange in *H. pylori[7]* and to play a crucial role in the pathogen persistence in the host and chronicity due to adaptation and immune response evasion[8,9]. Unlike other horizontal gene transfer pathways, NT only relies on molecular machineries coded by the recipient cell[10,11]. The initial event in NT consists in the capture of the exogenous DNA on the cell surface. It is then transferred into the cytoplasm through a two-step mechanism[12,13]. In naturally competent bacteria, uptake of the transforming DNA (tDNA) from the bacterial surface into the periplasm, through either the cell wall in gram-positive bacteria or the outer membrane in gram-negative, requires type IV pili or pilus-like structures. The tDNA is then bound by ComEA (ComH in *H. pylori*) and delivered, as a single-stranded DNA, to the inner membrane protein ComEC that, together with the cytosolic ComFC[14], mediates its transport into the cytosol. Remarkably, uptake to the periplasm in *H. pylori* is mediated by a unique set of type IV secretion system (T4SS) proteins rather than by type IV pili[15,16]. T4SS originated as a system for DNA exchange by conjugation in diderm bacteria to give rise to a variety of protein secretion systems present in both monoderm and diderm bacterial species[17]. One of the most surprising adaptations of T4SS to new functions is the ComB system from *H. pylori[18]*, required for NT. Indeed, ComB is the only T4SS studied so far that, instead of mediating a secretion process, mediates the internalisation of a macromolecule. Components of this simplified T4SS are coded by two independent operons, *comB2-comB4* and *comB6-comB10,* named after the homologous proteins in the well characterised VirB/D4 T4SS from *Agrobacterium tumefaciens[19].* However, when compared to the *A. tumefaciens* minimized system, some ComB components seem to be missing[20]. B1, involved in murein hydrolysis, might be replaced by the ComL homologue, required for NT in *Neisseria gonorrhoeae.* B1 could also be dispensable for the process since it is not essential for DNA export. In addition to B1, the *comB* operons do not code for VirB5, VirD4 or VirB11 homologues. While the absence of VirB5, a minor pilin, and VirD4, a traffic ATPase involved in substrate recruitment, could be expected in a T4SS mediating the internalisation of DNA rather than the secretion of macromolecules, that of a VirB11 homologue is surprising. VirB4 and VirB11 are ATPases localised at the cytoplasmic face of the inner membrane[21]. In the studied T4SS systems, these so-called secretion ATPases are essential for T4SS functions. Here we identified and characterised a gene coding for a VirB11 homologue essential for NT. The resulting protein is a *bona fide* ATPase, transiently associated with the bacterial membrane. Its interaction with ComB4 is indispensable for the entry of the tDNA into the periplasm. We propose therefore to name it ComB11.

## Results

### HP1421 is required for natural transformation

A homology search for VirB11-type ATPases pointed, in addition to the Cag-associated and well characterised HP0525 (Cagα), to the protein coded by *hp1421.* Sequence alignment of the two *H. pylori* genes with *A. tumefaciens* VirB11 show a high degree of homology and, in particular, a strong conservation of residues essential for the VirB11 and HP0525 ATPase activities (S1 Fig.). While HP1421 has been proposed to be a VirB11 homologue for the ComB system[20], a transposon insertion in codon 20 of *hp1421* did not result in a defect in plasmid transformation[22]. Since that mutant preserved most of the open reading frame (ORF), including the region harbouring the putative active sites, production of an active protein could not be ruled out. Furthermore, transformation with genomic DNA was not tested on that strain. We thus decided to create a null mutant by replacing the whole *hp1421* ORF by a non-polar antibiotic resistance cassette and test its capacity to be transformed by genomic DNA of a streptomycin resistant (Strep^R^) but otherwise isogenic strain.

Fig. 1A shows that inactivation of *hp1421* resulted in transformation levels below the detection limit of the assay, which is defined by the frequency of spontaneous Strep^R^ mutations. In order to rule out polar effects caused by the insertion of the cassette, we reintroduced in non-essential loci constructs expressing the *hp1421* ORF with a FLAG tag coding sequence fused 5’ of it. In the first case, the gene was under the control of the strong *ureA* promoter, while in the other, introduced into the *rdx* locus, the weaker *comH* promoter was used. In both cases, the NT capacity was completely restored to wild type levels, confirming the essential role of HP1421 in NT.

**Figure 1.**
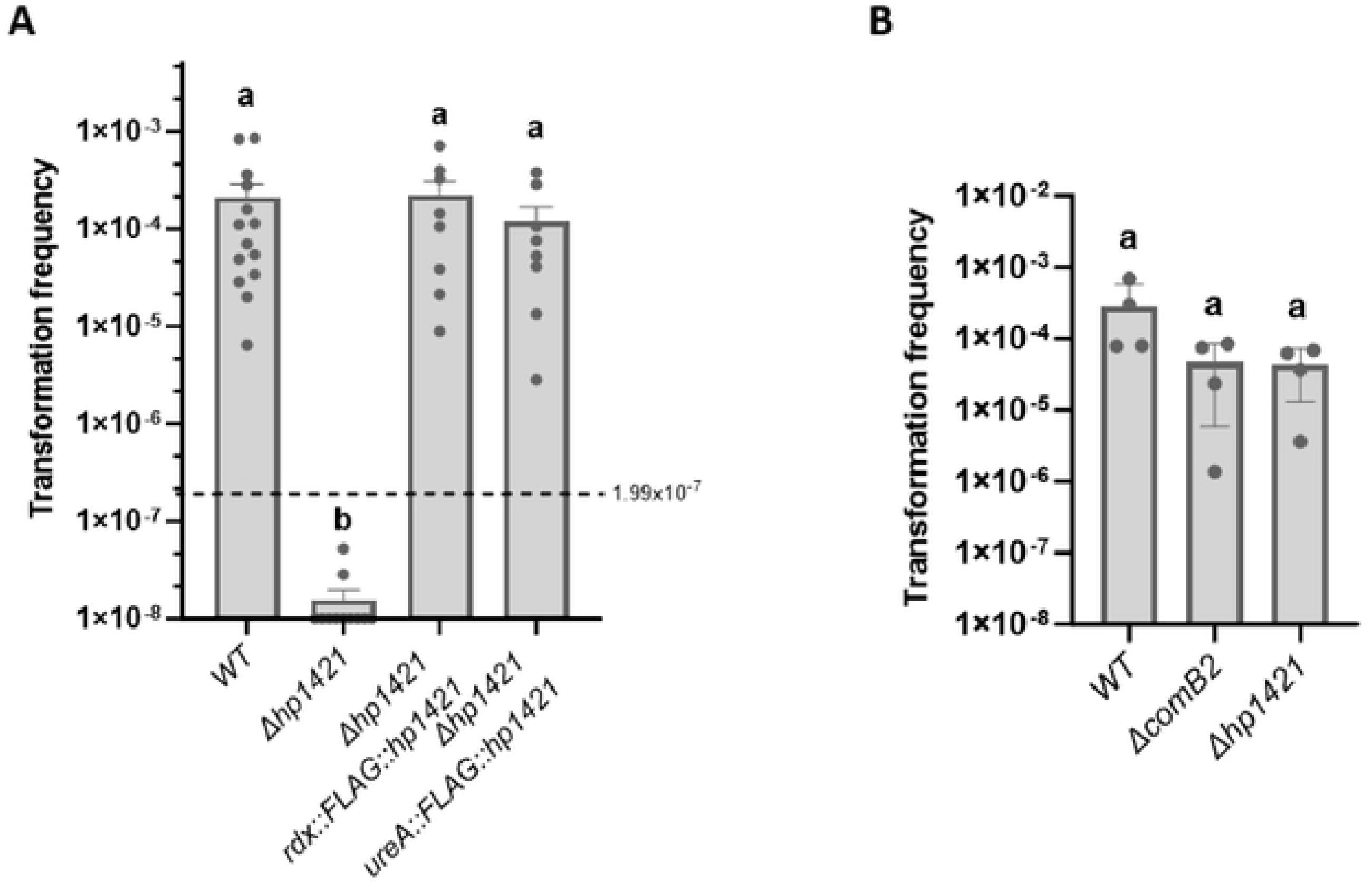
HP1421 is essential for natural transformation. **A)** Natural transformation frequencies for indicated *H. pylori* strains. Data represent Mean + SEM (n=7→17). Different lowercase letters indicate significant differences (p<0.05) between treatments (SNK test). The horizontal line at 1.99×10^-7^ indicates frequencies of revertant due to spontaneous mutations in absence of donor DNA in the wild type strain. **B)** Transformation frequencies after electroporation with streptomycin resistance genomic DNA donor. Data represent Mean + SEM (n=4). Different lowercase letters indicate significant differences (p<0.05) between treatments (Tukey’s Test).

### HP1421 participates in the initial steps of NT

The NT process can be divided in four distinct steps[10]: capture of the tDNA at the surface of the cell, its uptake into the periplasm, its transport through the inner membrane and finally its integration into the bacterial genome by homologous recombination. To establish where in this process HP1421 acts, we first tested the capacity of the Δ*hp1421* strain to be transformed by electroporation. As shown in Fig. 1B, introducing the tDNA by this method resulted in transformation frequencies of the Δ*hp1421* strain equivalent to those of the *wild type*, indicating that HP1421 is required at some point upstream of the recombination step, in agreement with the hypothesis of a role linked to the ComB system.

To confirm that HP1421 is required for the tDNA entry to the cell, we used the whole-cell duplex PCR-based DNA uptake assay that allows the detection of heterologous tDNA presence either in the periplasm or in the cytoplasm [14,23,24]. The assay showed that the tDNA cannot be detected inside the Δ*hp1421* cells but the signal was recovered after complementation of the knockout strain with HP1421 expressed from the *ureA* locus (Fig. 2A,B).

**Figure 2.**
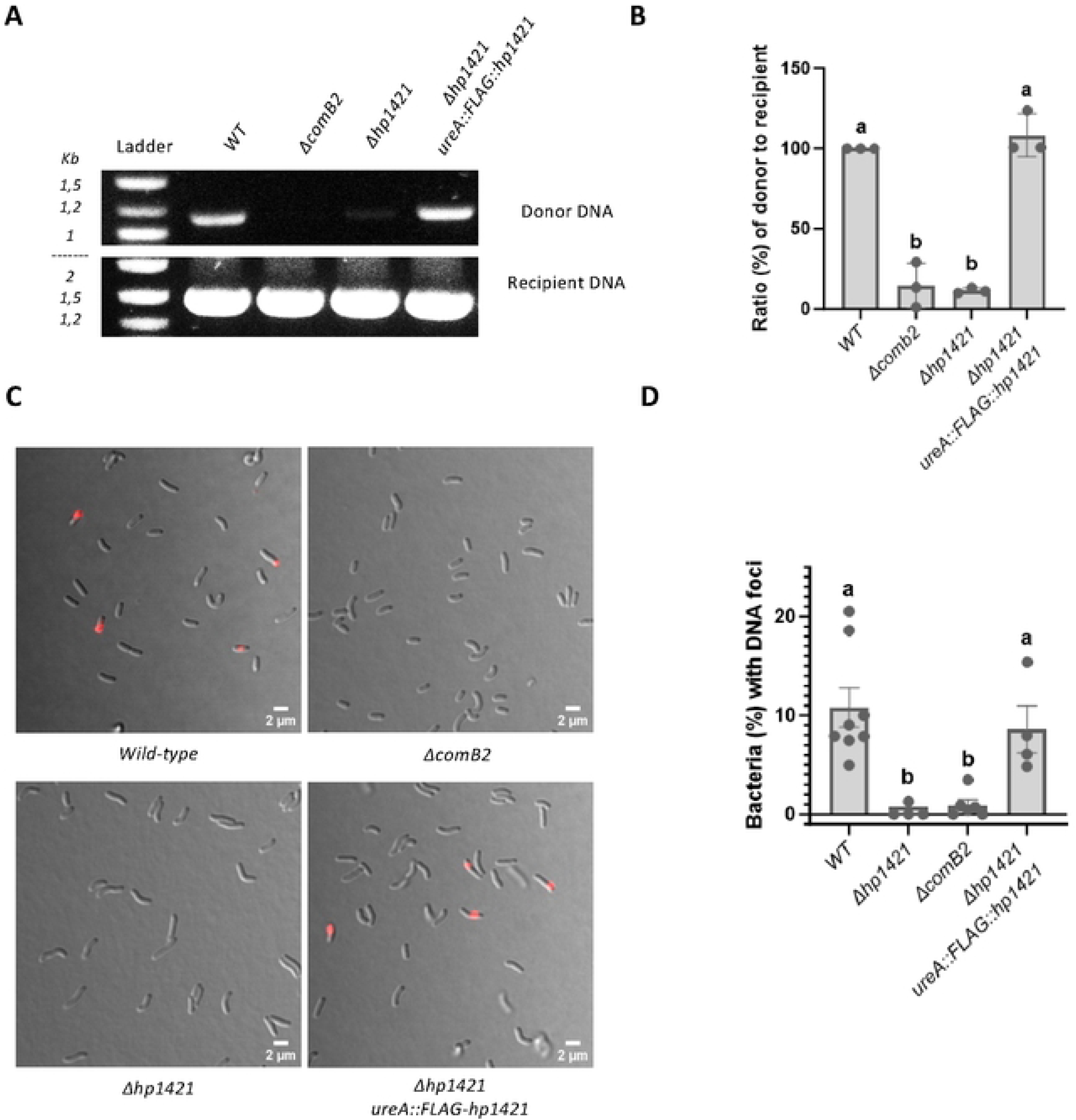
HP1421 is involved in capture and/or uptake of transforming DNA. **A)** Internalized donor DNA was monitored by PCR. Amplification products were visualized by 1% agarose gel electrophoresis. **B)** Percentage ratios of donor versus recipient amplification, results were normalized with respect to the wild-type strain. Data represent Mean + SEM (n=3). Different lowercase letters indicate significant differences (p<0.05) between treatments (Tukey’s Test). **C)** Fluorescent cy3-λDNA foci formation in *H. pylori wild-type, ΔcomB2 Δhp1421 and Δhp1421 ureA::FLAG-hp1421* strains. Z maximum projections of merged images of Cy 3.5 (red channel) and bright field are presented. **D)** Percentage of bacteria with fluorescent DNA foci. Data represent Mean + SEM (n=4→8). Different lowercase letters indicate significant differences (p<0.05) between treatments (Tukey’s Test).

Canonical T4SS systems provide a direct passage between the cytosol and the extracellular compartment. During NT, the delivery of the tDNA from the surface of the cell to the cytoplasm is a two-step mechanism[12,13]. In *H. pylori*, the uptake across the outer membrane is mediated by ComB while the passage through the inner membrane depends on the conserved ComEC protein[25,26]. If HP1421 is a VirB11 homologue associated with ComB, we expected HP1421 to be implicated in the first of those steps. To validate this hypothesis, we monitored the accumulation of tDNA in the periplasm using a fluorescently labelled tDNA. Previous work had shown that fluorescent tDNA foci are detected and stabilised in *H. pylori comEC* mutants[26,27]. In the absence of HP1421, periplasmic foci were undetectable, as in a Δ*comB2* mutant (Fig. 2C,D). Again, complementation by expression of the *wild-type* HP1421 protein resulted in the recovery of the foci. Taken together, these results implicate HP1421 in the initial step of NT leading to the delivery of the tDNA into the periplasmic space.

### HP1421 is a cytoplasmic ATPase

Sequence homology with ATPases of the VirB11 family (S1 Fig.) strongly suggests that HP1421 has an ATPase activity. To confirm this, we expressed in *Escherichia coli* the HP1421 protein fused through its N-terminal end to a maltose binding protein (MBP) tagged with six histidine residues (6-His-MBP-HP1421, herein MBP-HP1421). The fusion protein was purified (S2A Fig.) but attempts to obtain HP1421 without the MBP were unsuccessful, the protein precipitating after cleavage of the MBP. The ATPase activity was measured by determining the release of phosphate as a function of the substrate (ATP) concentration (Fig. 3A). MBP-HP1421 displayed a strong ATPase activity, with a calculated Km of 29.8 μM ATP, a value in agreement with those obtained for other enzymes of the VirB11 family[28,29]. In parallel, equivalent constructs were used to express Walker A motif mutants of the protein (E176A and E176K). Neither of those variants showed ATPase activity (Fig. 3A and S6A Fig.). Since the purification protocol was identical to the one used for the wild-type protein, these results rule out the presence of contaminating activities and confirm that HP1421 is a *bona fide* ATPase.

**Figure 3.**
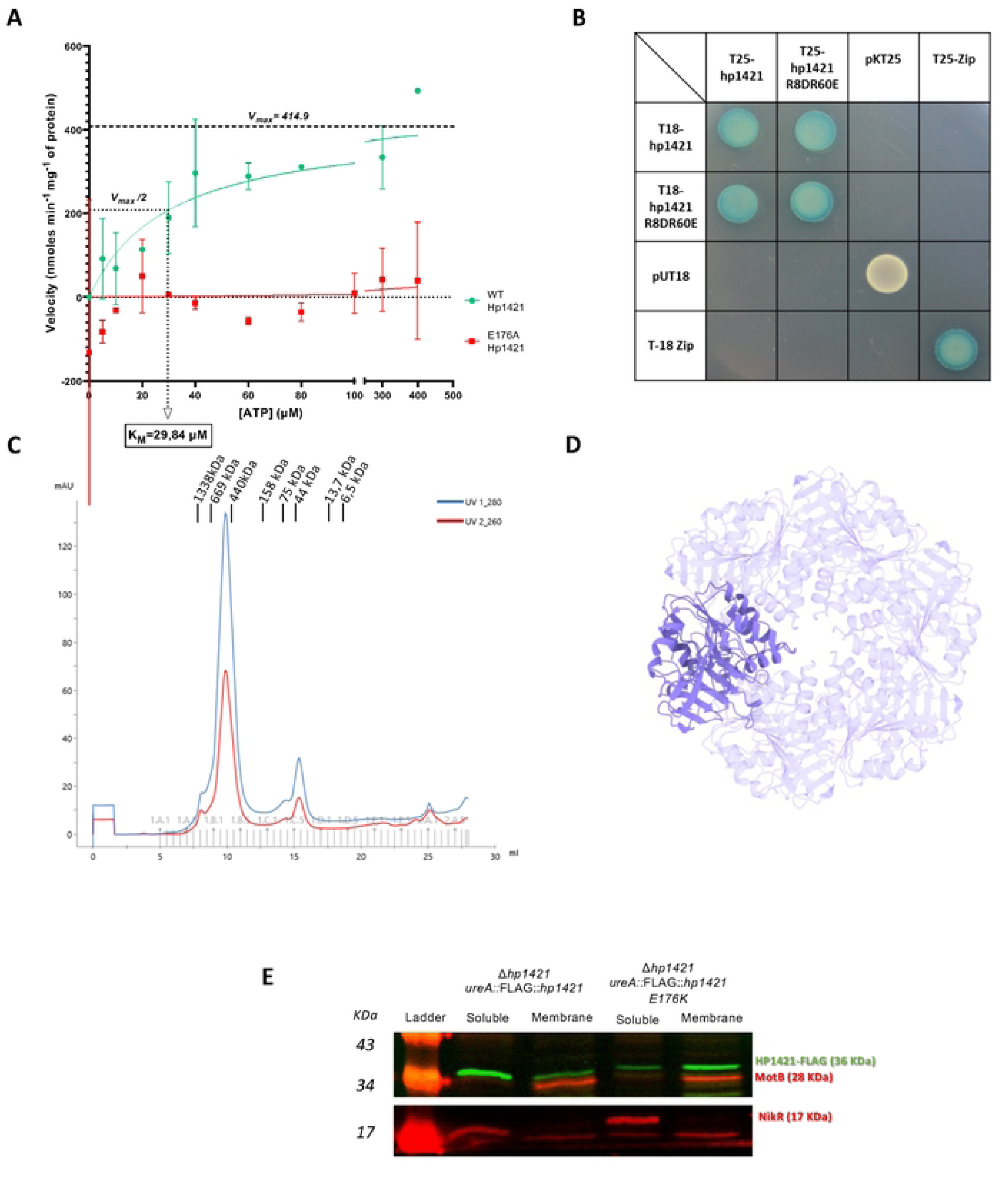
HP1421 is a cytoplasmic hexameric ATPase. **A)** ATPase activity of purified HP1421 versions as a function of ATP concentration, Pi released during ATPase enzymatic reactions was followed by spectrometry. **B)** BACTH analysis to assess the interaction between the indicated alleles of HP1421. **C)** Chromatogram for size exclusion chromatography (SEC) of 6xHis-MBP-HP1421 using a Superdex 200 column **D)** AlphaFold3 prediction made using 6 copies of HP1421 and 6 ATP molecules (pTM = 0.77, ipTM = 0.75, and chain-pair ipTM = 0.73 between any two monomers of HP1421 interacting in the model). One copy of HP1421 is highlighted from the model for visualization purpose. **E)** Western blot of subcelular fractions for indicated *H. pylori* strains.

As it is the case with most ATPases, HP1421 is expected to form multimers. To test whether the HP1421 monomers interact between themselves, a bacterial adenylate-cyclase two hybrid (BACTH) assay was carried out. This analysis revealed that indeed HP1421 form at least dimers (Fig. 3B). To establish the degree of multimerization, gel filtration assays on the purified MBP-HP1421 were performed with a Superdex 200 column. The main peak containing the fusion protein eluted at a volume corresponding to a molecular weight of approximately 460kDa which is close to that of an MBP-HP1421 hexamer, 473kDa (Fig. 3C, S2B Fig.). This is in agreement with a structural model of a HP1421 hexamer predicted by AlphaFold3[30] with a high level of confidence (Fig. 3D). This predicted HP1421 hexamer is structurally very similar to the experimental structure of the HP0525 hexamer (PDB identifier: 6BGE, S3A Fig.). Additionally, including 6 ATP molecules in the AlphaFold3 model of the HP1421 showed that ATP was confidently predicted as bound into a pocket on the surface of the HP1421 hexamer and in contact with residue E176 in the model (S3B Fig.).

T4SS-associated VirB11-like ATPases were reported to be localised in both, soluble and membrane fractions[31–33]. To examine HP1421 distribution within the cell, the soluble and membrane fractions from a *H. pylori* strain expressing the active FLAG tagged protein (Fig. 1A) were separated. Western blot analysis showed that while the majority of HP1421 is present in the soluble extract, a fraction of the protein can be detected associated with the membranes (Fig. 3E). When we analysed the localisation of HP1421 E176K, the balance between the two fractions was inverted, the majority of the mutant protein being associated with the membranes. Interestingly, in *Legionella pneumophila* Dot/Icm T4SS, DotB, the VirB11 homologue, is found essentially in the cytoplasm with only in a fraction of bacteria displaying DotB foci at the cellular pole. However, the E191K mutant, equivalent to HP1421 E176K and previously shown to bind but not hydrolyse ATP[34], was almost exclusively found at the bacterial pole[35].

The active site mutants provide a tool to determine if HP1421 ATPase activity is needed for its function in NT. We thus tested complementing the Δ*hp1421* strain phenotypes with either HP1421 E176K or HP1421 E176A expressed from the *ureA* promoter. Unlike that of the wild-type protein (Fig. 1A), the expression of the active site mutants did not allow the recovery of the transformation capacity (Fig. 4A) nor the accumulation of fluorescently labelled tDNA in the periplasm (Fig. 4B). Together, these results show that HP1421 is an hexameric ATPase whose enzymatic activity is indispensable for NT.

**Figure 4.**
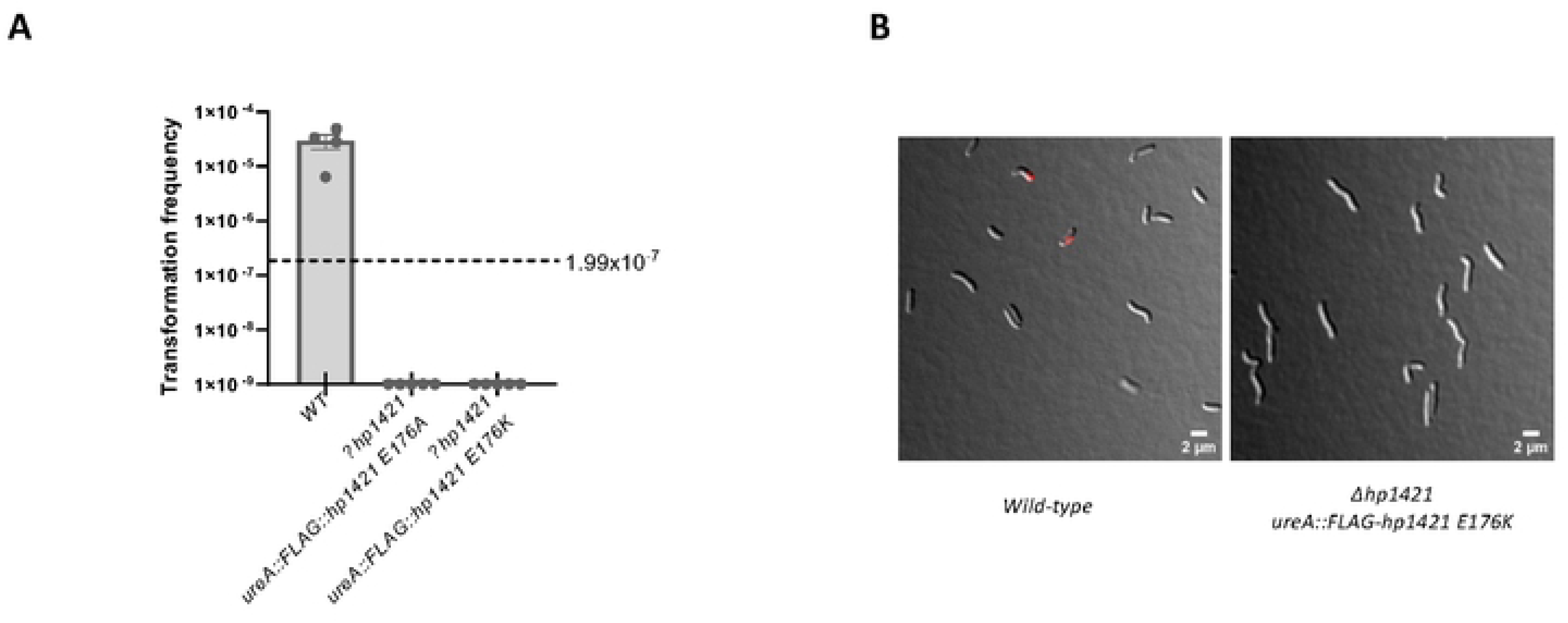
HP1421 ATPase activity is crucial to for NT. **A)** Natural transformation frequencies for indicated *H. pylori* strains. Data represent Mean + SEM (n=4→5), the horizontal line 1.99X10-7 indicates frequencies of revertant due to spontaneous mutations in absence of donor DNA in the wild type strain. **B)** Fluorescent cy3-λDNA foci formation in *H. pylori strains*. Z maximum projections of merged images of Cy 3.5 (red channel) and bright field are presented.

### HP1421 interaction with ComB4 is critical for NT

Structural and biochemical approaches revealed an interaction between VirB11 orthologues and their cognate VirB4 in various T4SS models[36–38]. In silico modelling of the *H. pylori* proteins predicted with good confidence that such an interaction could also occur for the HP1421-ComB4 pair. We performed AlphaFold3 modeling of the interaction between a HP1421 hexamer and a ComB4 C-terminal domain hexamer, yielding a very confident structural model (ipTM =0.81, pTM = 0.82, Fig. 5 and S4A Fig.). This model is highly structurally similar to models of the HP1421-ComB4 interface using single copies of the full-length proteins, obtained with high confidence by AlphaFold2[39,40]and AlphaFold3 (ipTM = 0.67 with AlphaFold2, 0.66 with AlphaFold3 and pTM = 0.78 with AlphaFold2, 0.75 with AlphaFold3) and to an AlphaFold2 model using only the N-terminal domain of HP1421 and the C-terminal domain of ComB4 (ipTM = 0.88, pTM = 0.89, S4B Fig.). The HP1421 ATPase activity is mediated by the C-terminal domain and the model hints that this activity should not be affected by the binding of HP1421 to ComB4. In addition, the HP1421-ComB4 modeled interface is perfectly compatible with the formation of a full-length ComB4 hexamer (S5 Fig).

**Figure 5.**
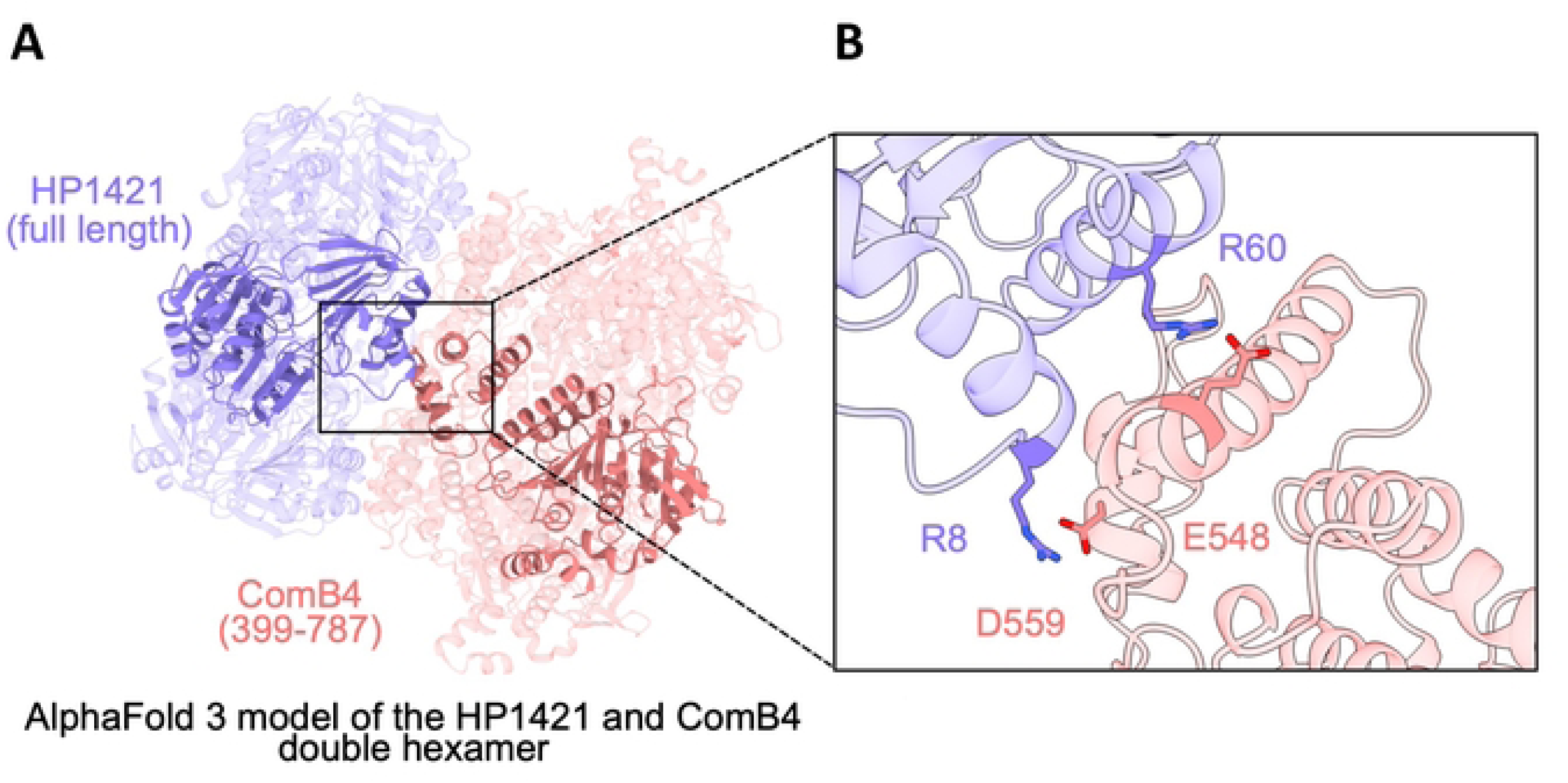
HP1421 and ComB4 double hexamer complex and interface. **A)** HP1421 full length (blue) and ComB4 C-terminal domain (salmon) hexamers confidently predicted as interacting with AlphaFold3 (pTM = 0.82, ipTM = 0.81). One copy of each protein was highlighted for visualisation purposes. **B)** Zoom on the interface of the AlphaFold3 heterodimeric model highlight important residues (R8 and R60 on HP1421, E548 and D559 on ComB4).

In order to empirically probe the interaction, a split luciferase assay was carried out with *H. pylori* strain expressing codon-optimized long (lgbit) and small (smbit) nanoluciferase fragments (S7 Fig.) fused to the N-terminal end of HP1421 and the C-terminal end of ComB4, respectively. Proximity between lgbit-HP1421 and ComB4-smbit would reconstitute the luciferase and produce a luminescent signal. This was indeed the case, strongly suggesting that HP1421 and ComB4 interact in the cellular context (Fig. 6A). The model presented in Fig. 5 displayed a strong electrostatic complementary of the interaction interfaces (S8 Fig.). We hypothesized that switching charges in some key interface residues could abolish the interaction and thus validate the model and provide a tool to assess the biological role of the interaction. Two amino-acid changes were chosen for each of the proteins based on the structural model of the HP1421-ComB4 interface (Fig. 5). For HP1421, arginine 8 and 60 were substituted by aspartic acid and glutamic acid, respectively. For ComB4, glutamic acid 548 and aspartic acid 559 were replaced by arginine. The substitutions were introduced in the split luciferase constructs and the assays were carried out. As shown on Fig. 6A, substituting either the arginine on the N-terminal domain of HP1421 with acidic amino-acids or the acidic amino-acids in the C-terminal domain of ComB4 with arginine abolished the luminescent signal, indicating a disruption of the interaction.

**Figure 6.**
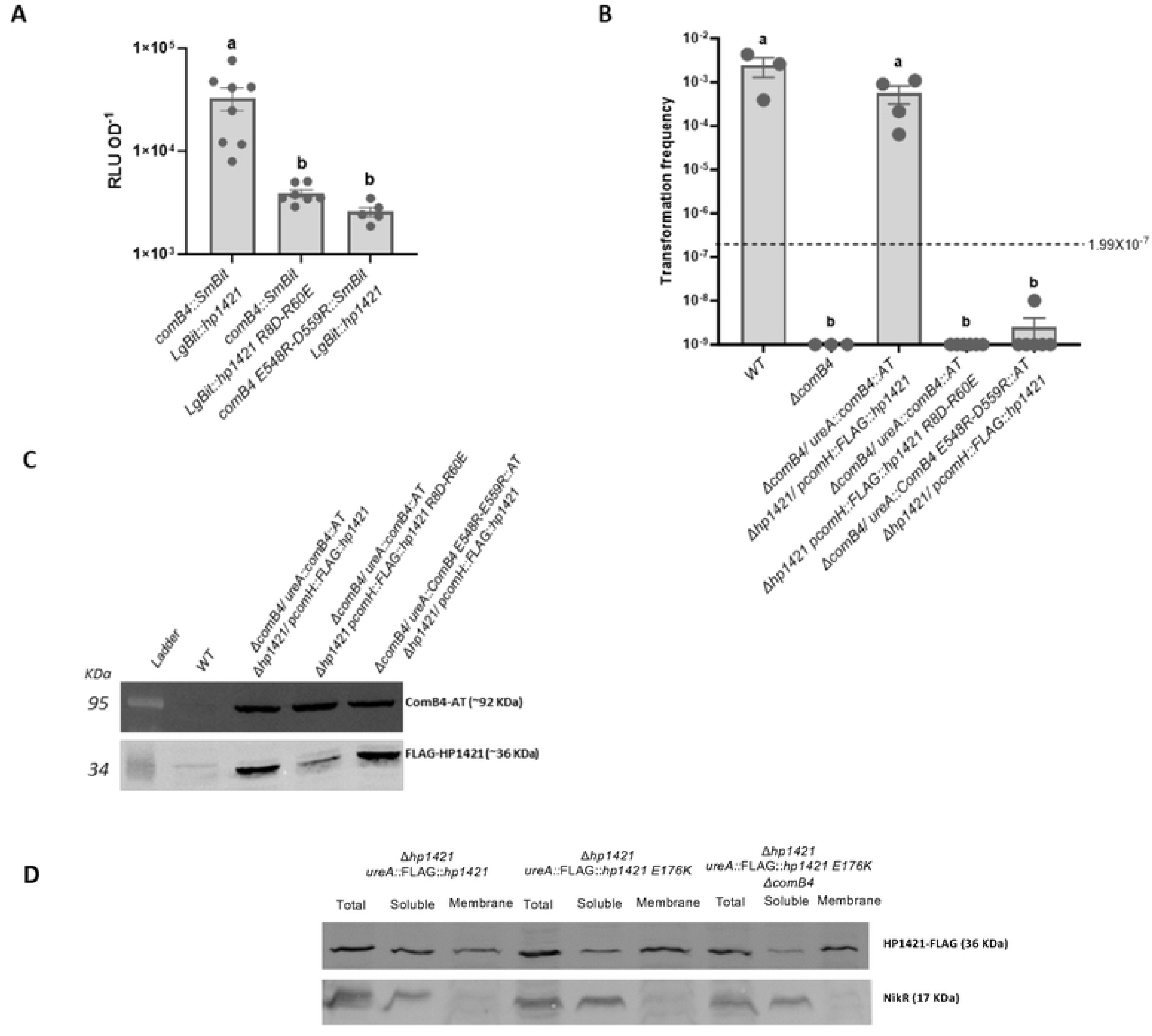
HP1421-ComB4A interaction is necessary to allow NT. **A)** Split-luciferase assay of interaction between ComB4::SmBit and LgBit::HP1421 variants, data represent Mean + SEM (n=5→8). Different lowercase letters indicate significant differences (p<0.05) between treatements (Tukey’s test). **B)** Natural transformation frequencies for indicated *H. pylori* strains. Data represent Mean + SEM (n=3→6). Different lowercase letters indicate significant differences (p<0.05) between treatments (SNK test). The horizontal line 1.99X10^-7^ indicates frequencies of revertant due to spontaneous mutations in absence of donor DNA in the wild type strain. **C)** Western blot of total fractions for indicated *H. pylori* strains. **D)** Western-blot of subcelular fractions for indicated *H. pylori* strains.

We then tested those interaction mutants for their capacity to sustain NT. For that purpose, we constructed *H. pylori* strains that expressed either both wild-type genes or one or the other interaction mutants. A strain expressing the HP1421 R8D-R60E variant was completely impaired for transformation (Fig. 6B,C). The same result was obtained with the ComB4 E548R-D559R variant (Fig. 6B,C). To rule out an effect of the amino-acid substitutions on the biochemical functions of HP1421, we expressed in *E. coli* and purified MBP-HP1421 R8D-R60E to assess its enzymatic activity and its capacity to assemble as hexamers, two properties required for its function in other T4SS[41]. As shown in S6A Fig., HP1421 R8D-R60E was able to hydrolyse ATP with the same specific activity than that of the wild-type protein. When the purified mutant was subjected to a gel filtration assay to determine its oligomerization status, the elution profile obtained indicated the presence of an approximately 460 KDa complex corresponding to a hexamer, as it was found for the wild-type protein MBP-HP1421 (S6B Fig.). Taken together, these results reinforce our confidence in the proposed structural model and show that the HP1421 interaction with ComB4 is essential for the functionality of the ComB system.

All the results presented so far indicate that HP1421 is indeed the VirB11 homologue linked to the ComB T4SS. We therefore propose herein naming it ComB11.

### The association of ComB11 with the membrane is not dependent on ComB4

The polar localization of DotB E191K was dependent on DotO, the VirB4 orthologue[35]. In view of the above results revealing the interaction between ComB11 and ComB4, we asked whether the enrichment of ComB11 E176K at the membranes was affected by the absence of ComB4. Western blots of the membrane and soluble fractions obtained from a Δ*comB4* strain expressing the mutant ComB11 showed the same distribution of the ATPase than in a ComB4 proficient strain (Fig. 6D) indicating that the interaction between the two ComB components does not drive the association, possibly transient, of ComB11 with the inner membrane.

### *comB11* is a core gene in *H. pylori* whose Com system is recent

To analyse the distribution of the ComB system and how it relates to ComB11, we analysed the 364 completely assembled genomes of *Helicobacteraceae* from RefSeq[42]. To be noted, 326 of these genomes are from *H. pylori*. We first tested that the protein ComB11 in the focal strain (26695) is sufficiently similar to the VirB11 protein profile of the T4SS model of TXSScan[43] to be regarded as a *bona fide* homolog. This is indeed the case, since it was significantly matched by the Virb11 protein profile of TXSScan (i-value=8×10^-39^). We then took every protein of the ComB system and searched for homologs in all the genomes of *Helicobacteraceae* by sequence similarity using diamond blastp[44]. The results show that most of the genes encoding ComB are very frequent in *H. pylori*. The least common genes are the *comB7* homologs present in only 2/3rds of the genomes. It was previously observed that *virB*7 genes are often missed in genome scans because the gene is small and evolves fast[45]. The most common gene, and the only gene present in every single strain of *H. pylori*, is *comB11* (Fig. 7B). We identified *bona fide* homologs of most of these genes in two other *Helicobacteraceae*: *Helicobacter cetorum* and *Helicobacter acinonychis*, the two closest species in the family for which we had complete genomes[46]. Systems associated with natural transformation are known to be occasionally defective in strains of a species, *e.g. comM* of *A. baumanni* was found to be defective or missing in more than 20% of the strains[47]. Yet, when present, the components of orthologous systems are expected to have similar genetic organisation. In the focal strain, the ComB system was split in three loci, one locus with *comB2-B3*-*B4*, another locus ca. 20 genes apart with *comB6*-*B7*-*B8*-*B9*-*B10*, and finally *comB11* which is distant from the others. We observed that the organization of each locus is highly conserved. When genes were present, the two first loci were strictly conserved in both *H. pylori* and the two close related species. Yet, relative distance between the three loci is variable (Fig. 7C) and they are never contiguous (minimal distance between the *comB4* and the *comB6* loci is 18 genes). The gene *comB11* was always distant from the others, although in many genomes it is between 30 and 40 genes apart from the *comB6* locus. Hence, the ComB system is present in most *H. pylori* strains and the two closest species in a conserved genetic organization, suggesting that the acquisition of the system predated the divergence of these three species.

**Figure 7.**
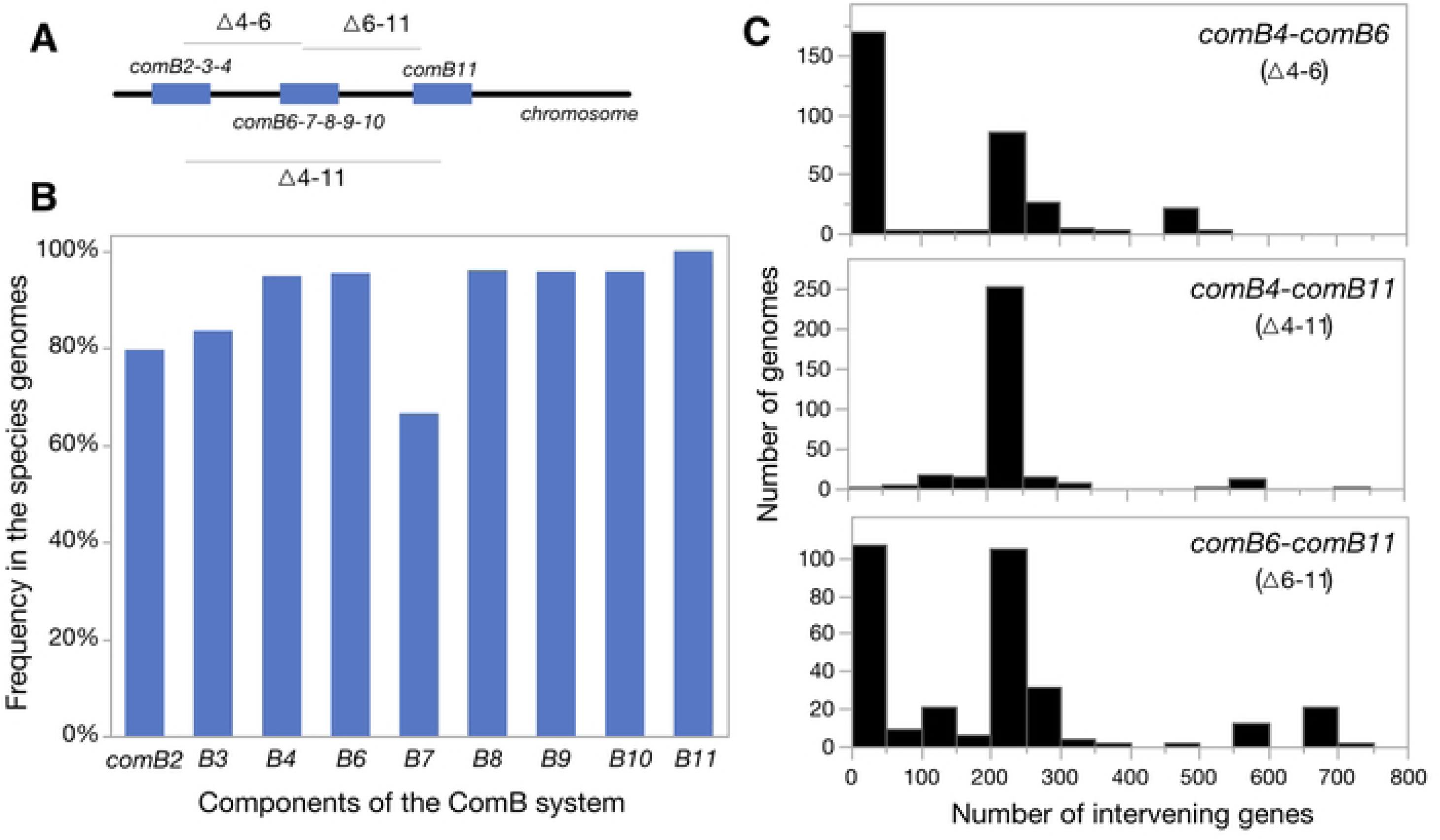
Frequency of the ComB system across *H. pylori*. **A)** Typical organisation of the loci. **B)** frequency of each gene in the strains of *H. pylori*. Only *comB11* is present in all the strains. **C)** Distances between key genes of each of the tree loci (see A for reference). Distances are computed as the minimal circular distance between the genes, counting the number of intervening genes.

We have observed homologs of *comB* genes in species more distant from *H. pylori*. *Helicobacter heilmannii* (strain type) encodes homologs of *comB2-3-4-6-8-9-11* and *Helicobacter felis* ATCC 49179 has homologs of *comB2-3-4-6-9-9-10-11*. In both genomes the genes are scattered in different loci, albeit the loci are not similar to those of *H. pylori* (e.g. in *H. heilmannii* comB6 and comB8 are isolate and separated from the locus with *comB9-10*). These scattered loci are almost certainly not part of a conjugative system of an integrative conjugative element, where they are usually co-localised in one single locus[48]. Further work will be needed to test if they are driving natural transformation in these species. Yet, we have observed that all these genomes lack a homolog of the ComH protein with the N-terminal domain, which was shown to be absolutely required for transformation in *H. pylori*[49] and was only identified in this species, *Helicobacter cetorum* and *Helicobacter acinonychis*.

## Discussion

During natural transformation, *H. pylori* can efficiently import double-stranded DNA from the milieu into the bacterial periplasm[12,25], where it is bound by the DNA receptor ComH[49]. While in other naturally transformable species characterised so far this process of capture and uptake of the tDNA is mediated by type IV pili or pilus-like structures[10], in *H. pylori* it relies on a T4SS, ComB[15,16,50]. This is a surprising finding since T4SSs are known as nanomachines that span a donor bacterium cell envelope to transfer macromolecules into target cells, whereas in the case of ComB it mediates the internalisation of the tDNA. Moreover, most T4SSs export proteins or DNA directly from the cytoplasm, while in *H. pylori* ComB delivers the incoming DNA into the periplasm, carrying out the first step of the two-step translocation mechanism mediating the passage of the DNA through the cell wall[25]. This process is completed by the transformation-specific ComH and ComEC proteins. ComH binds the DNA exiting ComB and delivers it to ComEC that, together with ComFC, allows the passage of the DNA through the inner-membrane[46]. This unusual function, mediating the uptake rather than the secretion of DNA, raises the question of how this particular T4SS works. ComB components coded by the two *comB* operons previously described[15,16] point to a minimised T4SS structure comparable to the canonical *A. tumefaciens* VirB system[51]. However, even for a minimized system, the lack of a VirB11 homologue in a Gram-negative T4SS was puzzling[19]. The results presented above allowed the identification of this key component of the ComB system required for the capture and uptake of DNA for natural transformation of the *H. pylori*.

The work presented here indicates that from the structural and biochemical points of view and the consequences of its inactivation, HP1421 is clearly a *bona fide* VirB11 homologue essential for the functioning of the T4SS ComB system. In accordance with the proposal to unify the nomenclature of the conserved T4SS subunits[19] we propose to name it ComB11.

In strain 26695 ComB11 is encoded by a gene distant from the two operons coding for the rest of the ComB genes. Even if the distance to the other *comB* genes is variable, this is true for all *H. pylori* strains for which assembled genomes are available. Since T4SSs are usually encoded in a single locus, this suggests that after the system was acquired and coopted to participate in natural transformation, its encoding locus was split and relative positions between the novel loci depend on the larger chromosomal rearrangements frequently observed in the species[52]. Similar cases of secretion systems loci being split have been identified before, but they are rare[53]. Amongst the *Helicobacteraceae* species for which complete genomes are available, only two other, *Helicobacter cetorum* and *Helicobacter acinonychis,* the closest to *H. pylori,* have a similar set of *comB* genes. Interestingly, in those two strains, ComH, which is indispensable for *H. pylori* natural transformation[49], displays the highest degree of homology with the *H. pylori* (74% and 58% for *H. acinonychis* and *H. cetorum,* respectively) when compared to the other members of the family. Moreover, for these two species closest to *H. pylori*, the alignment of the homologs’ sequences is along all the length of the protein while for the others it is limited to the C-terminal region of ComH. Previous work showed that the N-terminal domain of ComH is absolutely required for transformation[49]. Together these observations suggest that *Helicobacteraceae* other than *H. pylori, H. acinonychis* and *H. cetorus* are either non-transformable or use a system different than ComB/ComH.

Transcriptome analyses indicated that *hp1421* is downregulated in bacteria exposed to low pH[54], similarly to what was observed for *comB8* and *comH*[55]. Interestingly, development of competence requires an increase in pH of the culture[56,57]. The levels of other ComB components were shown to be limiting for the transformation capacity of *H. pylor*i[58]. Our results thus suggest that the expression of ComB11 could impact *H. pylori* transformation capacity by determining the availability of the tDNA uptake machinery.

Previous studies pointed to a potential association of HP1421 with *H. pylori* infection capacity. Indeed, *hp1421* mutants were unable to colonise the stomach in a Mongolian gerbil model[59]. Because *hp1421* is within a conserved operon harbouring genes involved in flagellar synthesis and export, a defect in flagellum formation could explain the lack of virulence provoked by the HP1421 absence. However, non-polar mutants of *hp1421* displayed normal flagella and motility[60]. While a defect in natural transformation was shown to affect *H. pylori* infection capacity in a mouse model[8], deletion of the *comB8-10* operon did not impair gerbil colonisation[59]. These observations suggest that, besides its essential role in natural transformation described in this work, HP1421 could have so far undefined functions required for infection. Such a possibility is supported by our finding that while *comB11* is a core gene present in all *H. pylori* strains, this is not the case for the other *comB* genes.

What is ComB11 function within the ComB T4SS? It had been proposed that the energy required for assembly and the retraction of a hypothetical pseudopilus capable of binding DNA, and therefore deliver it into the periplasm, is provided solely by ComB4[19]. ComB11, through its enzymatic activity and its interaction with ComB4 would complete the energy hub linked to the inner membrane complex. In other systems, the double-ring ATPase complex (VirB4/11) generates the energy required for pilus biogenesis[61]. Consistently with the lack of a VirB5 homologue which is required not only for adhesion of conjugative pili to target cells, but also in the initiation of pilus extrusion from the cell[19,21], no pilus structure associated with natural transformation has been described for *H. pylori.* However, the ComB system does harbour a pilin homologue, ComB2. It is tempting to speculate that ComB4/11 provides the energy to extract ComB2 from the inner membrane to form a polymer capable of reaching the surface of the cell where it could interact with a yet unidentified DNA-binding protein (Fig. 8). In any case, energy is certainly required for the uptake of the transforming DNA through the outer-membrane and into the periplasm. The ComB4/11 complex could also provide such energy for the internalisation of the DNA through the retraction of a pseudopilus structure formed by ComB2 (Fig. 8). T4SS pilus retraction has only been shown for the system encoded by the F plasmid[62,63], but the mechanisms underlying this process remain to be determined. A dual function implicating VirB11 as a switch between pilus biogenesis and DNA transport has been proposed[64]. The finding of the VirB11 homologue for the ComB T4SS provides a new model in which to analyse the role of these ATPases.

**Figure 8.**
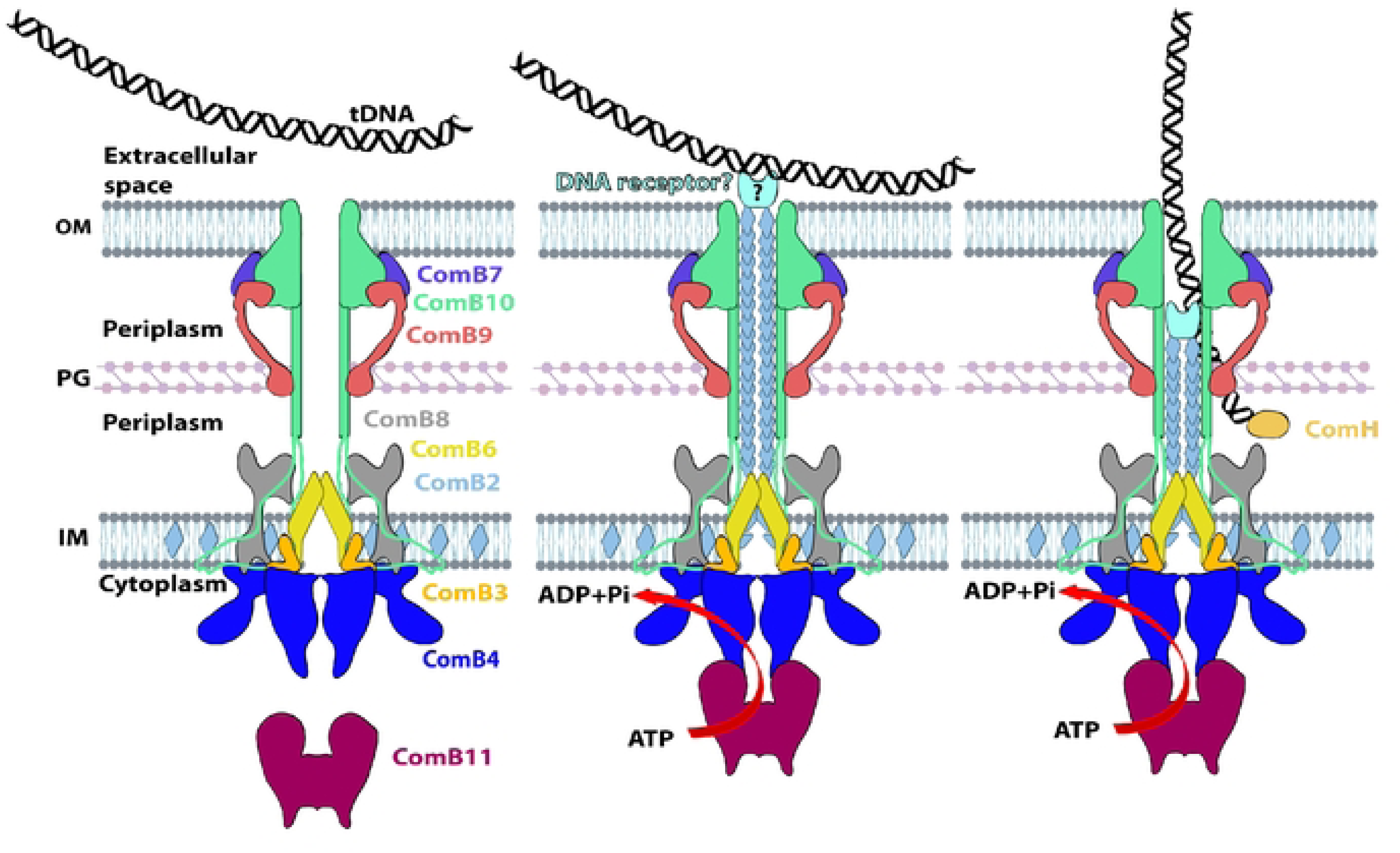
Model for the ComB role during natural transformation. See discussion section

## Materials and methods

### Strains culture

*Helicobacter pylori* 26695 strains listed on Supplementary Table 1 were cultured on blood agar base (Millipore) or brucella broth base (Millipore) supplemented with 1,5% bacteriological agar (Sigma-Aldrich), both enriched with 10 % of horse defibrinated blood (Thermo Scientific Oxoid). For liquid cultures, brain heart infusion (BHI) (Millipore) media was supplemented with 10% heat-inactivated fetal calf serum (FCS) (Eurobio scientific). All media were supplemented with an antimicrobial cocktail containing polymyxin B (0.155 mg/ml), vancomycin (6.25 mg/ml), trimethoprim (3.125 mg/ml), and amphotericin B (1.25 mg/ml) (all from Sigma/Sigma-Aldrich). Bacteria were grown at 37 °C under microaerophilic conditions obtained with the Whitley Jar Gassing System (Don Whitley scientific, United Kingdom) and a standardized gas mixture (3% O2, 10% CO2) (Air Products, France) using agitation when required (150 rpm) (INFORS HT Ecotron-Switzerland). When indicated, media were supplemented with kanamycin (20 μg/ml) (Sigma), apramycin (12.5 μg/ml) (Sigma), chloramphenicol (8 μg/ml) (Fisher) and/or streptomycin (10 μg/ml) (Sigma).

*Escherichia coli* strains used in this study are listed in Supplementary Table 2. Cultures used were carried out in liquid or solid Lysogeny broth (LB-Sigma) supplemented with the adequate antibiotic (ampicillin 100 µg/mL or kanamycin 50 µg/ml).

### Plasmid constructions

Constructions for gene disruption or complementation in *H. pylori* strains were carried out by using Sequence and Ligation Independent Cloning (SLIC) reaction[65]. Simultaneous and independent PCR reactions were performed in order to obtain an insert and a vector with overlapping sequences bigger than 18 bp. Briefly, 150 ng of the purified vector were mixed with the insert following a 1:5 ratio, 2 µL of NEBuffer r2.1 (New England Biolabs), 0.25 µL of T4 DNA polymerase (New England Biolabs) and H2O up to a final volume of 20 µL. Sample was incubated at room temperature 6 min and reaction stopped by using 2 µL of dCTP 10 mM, followed by a 15 min incubation at 37 °C. Right after 7 µL were used to transform 50 µL of chemo-competent DH5-α *E. coli* cells. Plasmids and oligonucleotides used in this study are listed in supplementary Table 3 and 4 respectively.

### *H. pylori* gene inactivation

Genomic sequences including 200 bp upstream and downstream of the gene of interest (chr: 1490990 bp to 1491904 bp for *hp1421;* chr: 14248 bp to 16611 bp for *comB4*) were amplified by PCR and cloned in pJET 1.2 backbone using SLIC reaction. A kanamycin non-polar cassette was used to replace the open reading frame for hp1421 gene and apramycin for *comB4*. Reference sequences were obtained from https://mage.genoscope.cns.fr/ *Helicobacter pylori* 26695 - chromosome NC_000915.1[66].

### Complementation

pJET 1.2 plasmids containing loci *ureA* (Chr: 77653 bp to 78221 bp) or *rdx* (Chr: 1013570 bp to 1014169) fused to the *comH* promoter (Chr: 1607419 bp to 1607618 bp) in position 1013912 bp were used for *hp1421* ectopic complementation. For *ureA* locus, the gene of interest was inserted on position 77956 bp. Alternatively, *hp1421* ORF was inserted downstream of the *comH* promoter. For both constructs a chloramphenicol non-polar cassette was inserted right after the gene. A FLAG tag was added at N-terminal side to follow up expression. Regarding constructs to test interacting comB4 and hp1421 mutants, *comB4* gene (Chr: 14248 bp to 16611 bp) was inserted on a pJET 1.2 plasmid containing loci *ureA* (Chr: 77653 bp to 78221 bp) at position 77956 bp. Adjacent to the gene, the previous described construct “*rdx*-p*comH* promoter-FLAG-*hp1421*-chloramphenicol cassette” was fused. An Alfa-tag was added at C-terminal side of *comB4* to follow up expression. Different *hp1421* and *comB4* gene variants were obtained by site directed mutagenesis[67] of these plasmids. Introduction of the various constructs in *H. pylori* was carried out by natural transformation or electroporation.

For natural transformation, plate-cultured *H. pylori* cells were taken to 4 OD600 in 40 µL of peptone water, containing 600 ng of plasmidic or genomic DNA harboring the construct of interest. The mixture was spotted in two 20 µL aliquots on blood agar brucella and incubated at 37 °C overnight in microaerophilic conditions. In some cases, complementation of KO strains deficient for the first steps of transformation was performed by electroporation (detailed in “*Helicobacter pylori* electroporation”). Grown spots were spread separately on blood agar brucella plates supplemented with the corresponding antibiotic and incubated until colonies were observed (4 to 6 days). Isolated colonies were transferred to new plates as patches keeping always the selection to escalate the inoculum to finally spread over half plate. Finally, a small fraction of cells was collected to extract gDNA by resuspension with 100 µL 50 mM NaOH followed by 20 minutes heating at 95 °C and a 5 minutes cooling with the addition of 20 µL 1 M Tris-HCl pH 7. PCR verifications were performed by using Kit Taq’Ozyme OneMix (Ozyme). For the complementing constructs, the genes were amplified using Q5 DNA high fidelity polymerase (New England Biolabs) and sequenced (Eurofins Genomics, Cologne, Germany).

### Transformation efficiency tests

Plate-cultured *H. pylori* strains were taken to OD600 = 4 in 45 µL final volume of peptone water, containing 600 ng of genomic DNA from a streptomycin resistant 26695 strain. 15 µL aliquots were spotted by triplicate over blood agar brucella and incubated overnight in microaerophilic conditions at 37 °C. Each spot was collected in 500 µl of peptone water and decimal dilutions were performed (up to 10^-6^). 10 µL aliquots were spotted by triplicate on blood agar brucella plates in absence (dilutions 10^-5^ and 10^-6^) or presence of streptomycin (non-diluted to 10^-3^). Plates were incubated until colonies appeared and CFU/mL were calculated (n° of colonies * dilution factor * aliquot factor). Transformation efficiency values were finally calculated by dividing the transformant CFU/mL over the total number of bacteria.

### *H. pylori* electroporation

Plate-cultured *H. pylori* cells were taken to OD600 = 10 in 1 mL final volume of peptone water, pelleted and resuspended in 1 mL of sterile ice-cold electrocompetent solution (9% sucrose - 15% glycerol). After centrifugation pellets were resuspended in 250 µL electrocompetent solutionr and 1/5 of the reaction was transferred to an electroporation cuvette and mixed with 600-1000 ng plasmidic or genomic DNA. Electroporation was performed on ice, at 2.5 kV cm−1; 25 µF; 200 Ω (Gene Pulser II, Bio-Rad), right after 100 µL of BHI were added and 50 µL spotted onto brucella blood agar plates.

### Subcellular fractionation

Plate-cultured *H. pylori* cells were resuspended in peptone water. Cells were washed once with sterile PBS (Gibco) 1X and normalized to OD600= 10 in 1mL of buffer A (Tris-HCl 10 mM pH 7.5 (Sigma); 1X protease inhibitor cocktail (cOmplete, Mini, EDTA-free, Roche). Samples were sonicated on ice (amplitude 20%, 30s, 2s pulses, 2s intervals) using a 3 mm microtip (Branson Digital Sonifier 450) and centrifuged 14000 g 15 min at 4°C (Eppendorf 5417R). The supernatant was collected and 100 µL were stored as total extract. The remaining volume was ultra-centrifuged 119000 g 45min at 4°C (Beckman Coulter Optima XPN-100), and supernatant was stored as soluble extract. The pellet containing total membranes was washed using in 1 mL buffer A and centrifuged using the same conditions as before. The pellet was resuspended in Buffer C (Tris-HCl 10 mM pH 7.5; 1X protease inhibitor cocktail; 1 % N-Lauroylsarcosine) to constitute the membrane fraction. Protein identification in different bacterial compartments was done by western blot. Additionally, as control of the experiment MotB (membrane marker) and NikR (cytoplasmic marker) localization, was followed by using primary antibodies designed in rabbit and secondary Goat-anti-rabbit IgG IR700 (Advansta).

### Western blots

To verify protein expression in complemented strains, exponential phase liquid cultures (18-24 h) were normalized at OD600 = 10 in 1 mL of 20 mM Tris-HCl pH 7.5 at 4°C (Sigma), 1X protease inhibitor (cOmplete, Mini, EDTA-free, Roche), right after samples were sonicated on ice (amplitude 20%, 30s, 2s pulses, 2s intervals) using a 3 mm microtip (Branson Digital Sonifier 450) and centrifuged 14000 g 10 min at 4°C (Eppendorf 5417R). 15 µL of extract were mixed with 5 µL of fresh Laemmli 4X, and heated 5 min at 95 °C for HP1421 constructs and 50 °C 15 min for ComB4 ones. Total volume was loaded into 10-12% hand-casted acrylamide gels (Mini-PROTEAN® Tetra 2-Gel, BioRad) and run using Mini-PROTEAN Tetra Vertical Electrophoresis Cell at 0.75 A-150V-30W (PowerPac™ HC High-Current Power Supply, BioRad) until running front was out. Resolved gels were transferred to 0.2µM nitrocellulose membranes (Trans-Blot Turbo Mini 0.2 µm Nitrocellulose Transfer Packs, BioRad) using Mixed MW program on a Trans-Blot Turbo Transfer System (BioRad). Membranes were blocked using 1X PBS, 0.3% Tween 20 (Sigma), 1% BSA (Sigma-Aldrich). For FLAG-tagged proteins, a 1/5000 dilution of ANTI-FLAG M2 primary antibody produced in mouse were used (Merck), and a 1/10000 dilution of secondary Goat-anti-mouse IgG IR800 (Advansta). Regarding ALFA-tag tagged proteins, a 1/1000 dilution of FluoTag®-X2 anti-ALFA ATTO 488 antibodies designed in alpaca (NanoTag Biotechnologies) were used. Membranes were washed with 1X PBS, 0.3% Tween 20 and revealed with iBright FL1500 (Thermo Fisher Scientific).

### DNA Uptake

Exponential phase liquid cultures (18-24 h) were normalized at OD600 = 1 in 1 mL of BHI-FCS. After a washing step using BHI sterile media, 250 ng of heterologous DNA linear plasmid were added as donor to the standardized suspension in prewarmed BHI-FCS. Samples were incubated 15 min at 37 °C in agitation (180 rpm) under microaerophilic atmosphere. After incubation, suspensions were vortexed twice (2500 rpm/min) during 15 s and centrifuged (8000 rpm-3 min). Two washing cycles were performed using sterile PBS 1X (Gibco), vortexing at high speed between each step. In order to eliminate free DNA, a nuclease treatment was performed at 37 °C during 20 min by pellet resuspension in 500 µL of a mixture containing 10 mM MgCl2 (Merck), 50 UI/mL DNase I (Sigma) and 62.5 UI/mL Benzonase (Millipore) in 1X PBS. This step was followed by high speed vortexing, pelleting and 2 washing cycles as detailed before. Cells were resuspended in 200 µL of PBS 1X and 200 µL of phenol:chloroform:isoamyl alcohol (25:24:1) (Sigma) and vortexed 30 s at high speed. Samples were centrifuged 12000 rpm for 1min and 100 µL of the upper phase were collected in a new tube, after an equal volume of chloroform was added and DNA was extracted by vortexing-centrifugation. Two simultaneous PCR reactions using Kit Taq’Ozyme OneMix (Ozyme) were performed using oligos op284 and op285 to detect donor DNA (plasmid) and Op31 and Op32 (Table XXXX) for recipient DNA (gDNA). Amplified fragments were resolved in a 1% agarose gel stained with SYBR™ Safe (Invitrogen), and revealed with transillumination (EBOX VX5/20M, Vilber Lourmat). After image acquisition, by using Fiji software band intensity of each reaction was measured and background subtracted to obtain an absolute valor, with those values a ratio recipient/donor was performed and internalization percentages were calculated taking the wild-type strain as 100%.

### DNA foci detection

*H. pylori* cells collected from blood agar plates were used to inoculate BHI broth at OD600= 0.05-0.1. When cultures reached exponential phase (18-24h), OD was normalized to 4 in 20 µL and 200 ng of λ-DNA (New England Biolabs), labelled with Cy3 using Label IT Nucleic Acid Labeling Kit (Mirus Bio) according to manufacturer instructions, were added to the reaction. After 7 min at 37°C in agitation (150 rpm) and microaerophilic conditions, cells were pelleted and washed with 100 µL of BHI media. In order to remove free DNA, cell suspensions (50µL) were treated with 10 U DNase I (Sigma) 5 minutes at 37°C followed by high speed vortexing. Bacteria were washed once more and resuspended in 20µL BHI, 3.5 µL of this suspension were spread on µ-Slide 8 Well (Ibidi) and 400 µL of soft agar (BHI-FCS 1% agarose) was added to immobilize cells. 2D imaging was performed using an inverted-confocal Nikon A1 microscope at 37°C (The Cube 2, Life Imaging Services), 10% CO2, 3% O2 and 85% humidity (The Brick, Life Imaging Services); 100-x oil objective was used and z-stack 0.125 μm acquisition 1248×1248-pixel resolution was carried out. Image processing was done using ImageJ software. The used criteria included the manual counting of entire rod shape cells and red foci over the cells in the composite image of bright field and red channel (594 nm). After foci percentage were calculated (total n° of cells /n° of cells with DNA foci).

### Protein purification

*E. coli* BL21 (DE3) strains, transformed with pRSF duet containing N-terminal 6His-MBP tags fused to hp1421 gene in wild type, E176A, E176K and R8D-R60E versions, were cultured in 500 mL LB at an initial OD600 of 0.1 at 37°C in agitation (180 rpm). At OD600= 0.6-0.7, IPTG (Thermo Scientific) was added to 1 mM and cultures were incubated overnight at 18°C in agitation. Cultures were centrifuged and pellets resuspended in buffer 500 mM NaCl (Sigma), 50 mM Tris-HCl (pH 8) (Sigma), 10% glycerol (Sigma-Aldrich), 1 mM DTT (Sigma), 1 mg/mL lysozyme (Sigma), 1x cOmplete EDTA-free protease inhibitor (Roche). The amount of buffer was calculated using the following formula: [(DO600 in 1 ml* total volume of culture)/40]. After samples were sonicated on ice (amplitude 20%, 3min, 7s pulses, 7s intervals) using a 13 mm tip probe (Branson Digital Sonifier 450). The protein of interest was then purified from the soluble fraction, obtained by centrifugation at 20000 rpm 60 min at 4°C. 3 mL of equilibrated Ni-NTA Superflow resin (Qiagen) were incubated 3h at 4°C in agitation with the soluble fraction, right after the whole suspension was transferred to a 2.5 × 10 cm Econo-Column (Bio-Rad), and flowthrough was obtained through pumping at 25 rpm (Mini peristaltic pump, Gilson). After, washing steps with 1.5 column volumes was performed with buffer A (20 mM Tris pH8; 500 mM NaCl; 1 mM DTT; 10% glycerol; 20 mM Imidazole), followed by 1.5 column volumes of buffer B (20 mM Tris pH8; 100 mM NaCl; 1 mM DTT; 10% glycerol; 20 mM Imidazole). Proteins were eluted in steps in 2 mL fractions with: 20 mM Tris pH8; 100 mM NaCl; 1 mM DTT; 10% glycerol; 500 mM Imidazole. Purified fractions were aliquoted, and stored at −80 °C.

### ATPase activity

All tests were performed using the EnzChek® Phosphate Assay Kit (Molecular probes, Invitrogen), following the protocol provided by the company. To determine HP1421 Michaelis constant (Km), a concentration rank of 0-4 mM ATP was used with a fixed amount of protein. Briefly the reaction was incubated 30 min at 22 °C, and absorbance was read at 360 nm on a microplate reader (CLARIOstar – BGM Biotech). Released phosphate was calculated by interpolation and corrected by subtraction of ATPase contamination measurement in absence of ATP. Km values were calculated using nonlinear least squares regression (GraphPad Prism 10.0.1). Regarding specific activity determinations on HP1421 R8DR60E mutants, phosphate release was followed on 5 minutes intervals during 45 min at 37 °C in a rank of 1-10 µM of protein at a fixed 4 mM concentration of ATP. Obtained values were corrected as indicated before, and plotted in function of time, common linear intervals were selected and slopes calculated. Finally, the slope of slope versus protein concentration graph was calculated for each determined genotype.

### Bacterial two hybrid assay

T25*-hp1421* and T18-*hp1421* constructs designed in plasmids pKT25 and pUT18C (Euromedex) variants were co-transformed *E. coli* BTH101 chemocompetent cells. Two to three transformant colonies were resuspended in 100 µl of LB, and 10 µL were spotted on solid LB supplemented with 100 µg/mL Ampicillin, 50 µg/mL Kanamycin, 0.5 mM IPTG (Sigma) and 40 µg/mL X-gal (Roche). Plates were incubated at 30 °C 24h and images registered.

### Split luciferase constructs and tests

*comB4* gene (Chr: 14248 bp to 16611 bp) was inserted on a pJET 1.2 plasmid containing loci *ureA* (Chr: 77653 bp to 78221 bp) at position 77956 bp. Adjacent to this gene the construct “*comH* promoter (Chr: 1607419 bp to 1607618 bp), *hp1421* gene (chr: 1490990 bp to 1491904), chloramphenicol cassette” was added. Nano-luciferase optimized fragment sequences for *H. pylori* expression Sm-Bit and Lg-Bit[68] were fused to N-terminal hp1421 and C-terminal ComB4 respectively. Random 15 bp linkers were added between the segments and gene sequences to allow flexibility. Different *hp1421* and *comB4* gene variants were obtained by site directed mutagenesis of these plasmids. All luciferase tests were performed using the Nano-Glo® Luciferase Assay System (Promega). Briefly, plate-cultured *H. pylori* cells were taken to OD600 = 0.1 in BHI. 50 µL of each suspension were mixed with 50 µL of reconstituted reaction buffer, by duplicate in Nunc F96 MicroWell White Polystyrene Plates (Thermo Scientific). Plate was gently mixed by tapping and 3 minutes after, readings were performed each 30s up to 5 minutes final using a microplate reader (CLARIOstar – BGM Biotech). Averaged blank corrected values were summed and corrected by OD to normalize. To each replicate, wild type non-specific activity was subtracted to each condition.

### AlphaFold modelling

Most structural models were obtained using the AlphaFold3 web server[30]. HP1421 hexamer with ATP, HP1421-ComB4, ComB4 hexamer were modeled with full-length proteins. The HP1421/ComB4 double hexamer was modeled with shorter delimitations due to AlphaFold3 size limit: only the C-terminal domain of ComB4 was used (from the UniProt sequence reference O24862, residues 399-787 (end)). The HP1421-ComB4 interface was also modeled with AlphaFold2[39] using the ColabFold[40]implementation.

### Structural data visualization

Molecular graphics and analyses performed with the UCSF ChimeraX software[69], developed by the Resource for Biocomputing, Visualization, and Informatics at the University of California, San Francisco, with support from National Institutes of Health R01-GM129325 and the Office of Cyber Infrastructure and Computational Biology, National Institute of Allergy and Infectious Diseases. The command line version of matchmaker module was used to align protein chains and obtain RMSD values with 5.0 Å pruning iteration cut-off distance (to exclude outlier atoms). The “pair ss” parameter were applied when aligning complexes (hexamers) to ensure a global alignment instead of a single pair based one. The electrostatic potential at HP1421-ComB4 interface was estimated using ChimeraX “electrostatic” (coulombic) module. The plDDT colouring was obtained directly from AlphaFold plDDT values in the json output file.

### Size exclusion chromatography

The oligomerization state of the protein is determined by passage through a Superdex 200 column. Calibration of the gel filtration column is performed using a cocktail of eight proteins: aprotinin (6.5 kDa), ribonuclease A (13.7 kDa), carbonic anhydrase (29 kDa), ovalbumin (44 kDa), conalbumin (75 kDa), aldolase (158 kDa), ferritin (440 kDa), and thyroglobulin (669 kDa). The column is equilibrated with 20 mM Tris-HCl, pH 8.0, 100 mM NaCl, 1 mM DTT, and 10% glycerol. The purified protein (100 µg) is loaded onto the column, and the elution volume of the protein is determined. By plotting this value on the column calibration curve, the apparent molecular weight of the protein is calculated.

### Phylogenetic analyses

We extracted from RefSeq complete genomes[42] (S5 file) all the 364 genomes of *Helicobacteraceae* (last accessed in May 2023). We used the annotations of the genomes regarding the position of every protein coding gene and their functional annotation. We used the proteins of the focal strain (26695) to identify homologs in the other genomes using diamond blastp v2.0.4.142[44] with the *ultra-sensitive* option. Within the closely related species of *H. pylori, Helicobacter cetorum* and *Helicobacter acinonychis* we regarded proteins as orthologous if they were more than 80% identical. This threshold was verified upon the analysis of the loci loci, by comparing the hits with this threshold with those using a much lower threshold (e-value <0.001, down to less than 30% identity). This confirmed that only homologs at more than 80% identity are found in the loci. Once the list of orthologs was available we computed the minimal distance of each pair in the genome as the number of genes that were identified between them (accounting for the circularity of the genomes).

The test that comB11 is a *bona fide* virB11 homolog was done by checking that it can be identified by the virB11 HMM protein profile from TXSScan[43], using hmmsearch 3.1b2 with default options[70]. The protein profile is given in S6 File.

### Statistical analysis

Independent experiments were performed using a completely randomized design with at least three biological replicates. RStudio software and Graphad Prism 10 were used to perform statistical analysis. Each dataset was analyzed using one way ANOVA and Tukey’s test at P <0.05. Model assumptions were verified using Shapiro–Wilk and Levene’s tests, including Normal Q-Q-plot and residuals versus fits plot analysis. When residuals normality and homoscedasticity was not proven, boxcox function was used to choose the best data transformation. If assumptions weren’t still verified, non-parametrical Kruskal and Wallis test followed by post-hoc SNK test at P<0.05 (global P value adjusted to the number of comparisons) was performed.

## Data availability

All relevant data are available within the main text, the supporting information files and on Zenodo (https://doi.org/10.5281/zenodo.19238504). Data used for graphs and tables are available as a Data Source file. All other data can be requested to the authors.

## Acknowledgements

We thank Anne Marie Di Guilmi for the initial construction of the plasmids used for the BACTH assays, Laurent Terradot (CNRS, Lyon) for sharing plasmids with us and Hilde De Reuse (Institut Pasteur, Paris) for antibodies against NikR and MotB. We thank the BIOI2 core facility for making the ColabFold pipeline easily accessible at the I2BC to run AlphaFold2.

## Author contributions

Conceptualisation: JFV, JPR

Methodology: YF, RG, JA

Formal analysis: JFV, EPCR, YF, RG, JA

Investigation: JFV, SK, MM, CL, XV

Writing - Original Draft: JFV, JPR, EPCR, YF, JA

Writing – Review and Editing: JFV, SK, XV, EPCR, JPR, YF, RG, JA. All authors read and approved the manuscript.

Visualisation: JFV, JPR, YF

Supervision: JPR, JA

Project administration: JPR

Funding acquisition: JA, RG, JPR

## References

1. Good BH, Zuckerberg Biohub-San C, Mcdonald MJ, Ami j, Bhatt S, Mcdonald MJ. Unraveling the tempo and mode of horizontal gene transfer in bacteria. Trends Microbiol. 2025;33: 853–865. doi:10.1016/J.TIM.2025.03.009

2. Arnold BJ, Huang IT, Hanage WP. Horizontal gene transfer and adaptive evolution in bacteria. Nature Reviews Microbiology 2021 20:4. 2021;20: 206–218. doi:10.1038/S41579-021-00650-4

3. Soucy SM, Huang J, Gogarten JP. Horizontal gene transfer: building the web of life. Nat Rev Genet. 2015;16: 472–482. doi:10.1038/nrg3962

4. Woods LC, Gorrell RJ, Taylor F, Connallon T, Kwok T, McDonald MJ. Horizontal gene transfer potentiates adaptation by reducing selective constraints on the spread of genetic variation. Proc Natl Acad Sci U S A. 2020;117: 26868–26875. doi:10.1073/pnas.2005331117

5. Suerbaum S, Achtman M. Evolution of Helicobacter pylori: the role of recombination. Trends Microbiol. 1999;7: 182–184. Available: http://www.ncbi.nlm.nih.gov/entrez/query.fcgi?cmd=Retrieve&db=PubMed&dopt=Citation&list_uids=10383222

6. Suerbaum S, Smith JM, Bapumia K, Morelli G, Smith NH, Kunstmann E, et al. Free recombination within Helicobacter pylori. Proc Natl Acad Sci U S A. 1998;95: 12619–12624. Available: http://www.ncbi.nlm.nih.gov/entrez/query.fcgi?cmd=Retrieve&db=PubMed&dopt=Citation&list_uids=9770535

7. Fischer W, Windhager L, Rohrer S, Zeiller M, Karnholz A, Hoffmann R, et al. Strain-specific genes of Helicobacter pylori: genome evolution driven by a novel type IV secretion system and genomic island transfer. Nucleic Acids Res. 2010;38: 6089–6101. doi:10.1093/nar/gkq378

8. Dorer MS, Cohen IE, Sessler TH, Fero J, Salama NR. Natural Competence Promotes Helicobacter pylori Chronic Infection. Blanke SR, editor. Infect Immun. 2013;81: 209–215. doi:10.1128/IAI.01042-12

9. Suerbaum S, Ailloud F. Genome and population dynamics during chronic infection with Helicobacter pylori. Current Opinion in Immunology. 2023. doi:10.1016/j.coi.2023.102304

10. Dubnau D, Blokesch M. Mechanisms of DNA Uptake by Naturally Competent Bacteria. Annual Review of Genetics. 2019. pp. 217–237. doi:10.1146/annurev-genet-112618-043641

11. Johnston C, Martin B, Fichant G, Polard P, Claverys JP. Bacterial transformation: distribution, shared mechanisms and divergent control. Nat Rev Microbiol. 2014;12: 181–196. doi:10.1038/nrmicro3199

12. Kruger NJ, Stingl K. Two steps away from novelty--principles of bacterial DNA uptake. Mol Microbiol. 2011;80: 860–867. doi:10.1111/j.1365-2958.2011.07647.x

13. Seitz P, Blokesch M. DNA transport across the outer and inner membranes of naturally transformable Vibrio cholerae is spatially but not temporally coupled. mBio. 2014;5. doi:10.1128/mBio.01409-14

14. Damke PP, Celma L, Kondekar SM, Di Guilmi AM, Marsin S, Dépagne J, et al. ComFC mediates transport and handling of single-stranded DNA during natural transformation. Nat Commun. 2022;13. doi:10.1038/S41467-022-29494-Z

15. Hofreuter D, Odenbreit S, Haas R. Natural transformation competence in Helicobacter pylori is mediated by the basic components of a type IV secretion system. Mol Microbiol. 2001;41: 379– 391. Available: http://www.ncbi.nlm.nih.gov/entrez/query.fcgi?cmd=Retrieve&db=PubMed&dopt=Citation&list_uids=11489125

16. Karnholz A, Hoefler C, Odenbreit S, Fischer W, Hofreuter D, Haas R. Functional and topological characterization of novel components of the comB DNA transformation competence system in Helicobacter pylori. J Bacteriol. 2006;188: 882–893. doi:10.1128/JB.188.3.882-893.2006/ASSET/8DA3B1D6-B276-45C2-9752-C4AD8583D250/ASSETS/GRAPHIC/ZJB0030654120006.JPEG

17. Denise R, Abby SS, Rocha EPC. The Evolution of Protein Secretion Systems by Co-option and Tinkering of Cellular Machineries. Trends Microbiol. 2020;28: 372–386. doi:10.1016/J.TIM.2020.01.005

18. Guglielmini J, De La Cruz F, Rocha EPC. Evolution of Conjugation and Type IV Secretion Systems. Mol Biol Evol. 2013;30: 315–331. doi:10.1093/MOLBEV/MSS221

19. Christie PJ, Waksman G, Berntsson RP-A, Soler N, Leblond-Bourget N, Douzi B. Type IV secretion systems: reconciling diversity through a unified nomenclature. FEMS Microbiol Rev. 2026;50. doi:10.1093/femsre/fuaf069

20. Smeets LC, Kusters JG. Natural transformation in Helicobacter pylori: DNA transport in an unexpected way. Trends Microbiol. 2002;10: 159–62; discussion 162. Available: http://www.ncbi.nlm.nih.gov/entrez/query.fcgi?cmd=Retrieve&db=PubMed&dopt=Citation&list_u ids=11912014

21. Paillard P, Rouger Q, Thomet M, Macé K. Type IV secretion systems: from structures to mechanisms. The EMBO Journal 2025 44:22. 2025;44: 6304–6319. doi:10.1038/S44318-025-00584-0

22. Karnholz A, Hoefler C, Odenbreit S, Fischer W, Hofreuter D, Haas R. Functional and topological characterization of novel components of the comB DNA transformation competence system in Helicobacter pylori. J Bacteriol. 2006;188: 882–893. doi:10.1128/JB.188.3.882-893.2006

23. Suckow G, Seitz P, Blokesch M. Quorum sensing contributes to natural transformation of Vibrio cholerae in a species-specific manner. J Bacteriol. 2011;193: 4914–4924. doi:10.1128/JB.05396-11

24. Seitz P, Pezeshgi Modarres H, Borgeaud S, Bulushev RD, Steinbock LJ, Radenovic A, et al. ComEA Is Essential for the Transfer of External DNA into the Periplasm in Naturally Transformable Vibrio cholerae Cells. PLoS Genet. 2014;10: e1004066. doi:10.1371/journal.pgen.1004066

25. Stingl K, Müller S, Scheidgen-Kleyboldt G, Clausen M, Maier B. Composite system mediates two-step DNA uptake into Helicobacter pylori. Proc Natl Acad Sci U S A. 2010;107: 1184–1189. doi:10.1073/PNAS.0909955107

26. Corbinais C, Mathieu A, Kortulewski T, Radicella JP, Marsin S. Following transforming DNA in Helicobacter pylori from uptake to expression. Mol Microbiol. 2016;101: 1039–1053. doi:10.1111/mmi.13440

27. Stingl K, Müller S, Scheidgen-Kleyboldt G, Clausen M, Maier B. Composite system mediates two-step DNA uptake into Helicobacter pylori. Proc Natl Acad Sci U S A. 2010;107. doi:10.1073/pnas.0909955107

28. Arya T, Oudouhou F, Casu B, Bessette B, Sygusch J, Baron C. Fragment-based screening identifies inhibitors of ATPase activity and of hexamer formation of Cagα from the Helicobacter pylori type IV secretion system. Sci Rep. 2019;9. doi:10.1038/s41598-019-42876-6

29. Ripoll-Rozada J, García-Cazorla Y, Getino M, Machón C, Sanabria-Ríos D, de la Cruz F, et al. Type IV traffic ATPase TrwD as molecular target to inhibit bacterial conjugation. Mol Microbiol. 2016;100. doi:10.1111/mmi.13359

30. Abramson J, Adler J, Dunger J, Evans R, Green T, Pritzel A, et al. Accurate structure prediction of biomolecular interactions with AlphaFold 3. Nature. 2024;630: 493–500. doi:10.1038/S41586-024-07487-W,

31. Rashkova S, Spudich GM, Christie PJ. Characterization of membrane and protein interaction determinants of the Agrobacterium tumefaciens VirB11 ATPase. J Bacteriol. 1997;179: 583–591. doi:10.1128/JB.179.3.583-591.1997

32. Christie PJ, Ward JE, Gordon MP, Nester EW. A gene required for transfer of T-DNA to plants encodes an ATPase with autophosphorylating activity. Proc Natl Acad Sci U S A. 1989;86. doi:10.1073/pnas.86.24.9677

33. Kutter S, Buhrdorf R, Haas J, Schneider-Brachert W, Haas R, Fischer W. Protein subassemblies of the Helicobacter pylori cag type IV secretion system revealed by localization and interaction studies. J Bacteriol. 2008;190. doi:10.1128/JB.01341-07

34. Sexton AA, Yeo H-J, Vogel JP, Sexton JA, Yeo H-J, Vogel JP. Genetic analysis of the Legionella pneumophila DotB ATPase reveals a role in type IV secretion system protein export. Mol Microbiol. 2005;57: 70–84. doi:10.1111/J.1365-2958.2005.04667.X

35. Chetrit D, Hu B, Christie PJ, Roy CR, Liu J. A unique cytoplasmic ATPase complex defines the Legionella pneumophila type IV secretion channel. Nature Microbiology 2018 3:6. 2018;3: 678–686. doi:10.1038/S41564-018-0165-Z

36. Macé K, Vadakkepat AK, Redzej A, Lukoyanova N, Oomen C, Braun N, et al. Cryo-EM structure of a type IV secretion system. Nature 2022. 2022; 1–6. doi:10.1038/s41586-022-04859-y

37. Scheibner F. Functional characterization of VirB/VirD4 and Icm/Dot type IV secretion systems from the plant-pathogenic bacterium Xanthomonas euvesicatoria. Front Cell Infect Microbiol. 2023;13: 1203159. doi:10.3389/FCIMB.2023.1203159/BIBTEX

38. Hu B, Khara P, Song L, Lin AS, Frick-Cheng AE, Harvey ML, et al. In Situ Molecular Architecture of the Helicobacter pylori Cag Type IV Secretion System. mBio. 2019;10. doi:10.1128/MBIO.00849-19

39. Jumper J, Evans R, Pritzel A, Green T, Figurnov M, Ronneberger O, et al. Highly accurate protein structure prediction with AlphaFold. Nature 2021 596:7873. 2021;596: 583–589. doi:10.1038/s41586-021-03819-2

40. Mirdita M, Schütze K, Moriwaki Y, Heo L, Ovchinnikov S, Steinegger M. ColabFold: making protein folding accessible to all. Nat Methods. 2022;19: 679–682. doi:10.1038/s41592-022-01488-1

41. Savvides SN, Yeo HJ, Beck MR, Blaesing F, Lurz R, Lanka E, et al. VirB11 ATPases are dynamic hexameric assemblies: new insights into bacterial type IV secretion. EMBO J. 2003;22: 1969–1980. doi:10.1093/EMBOJ/CDG223

42. Haft DH, DiCuccio M, Badretdin A, Brover V, Chetvernin V, O’Neill K, et al. RefSeq: an update on prokaryotic genome annotation and curation. Nucleic Acids Res. 2018;46: D851–D860. doi:10.1093/nar/gkx1068

43. Abby SS, Cury J, Guglielmini J, Néron B, Touchon M, Rocha EPC. Identification of protein secretion systems in bacterial genomes. Sci Rep. 2016;6. doi:10.1038/srep23080

44. Buchfink B, Xie C, Huson DH. Fast and sensitive protein alignment using DIAMOND. Nat Methods. 2015;12: 59–60. doi:10.1038/nmeth.3176

45. Guglielmini J, De La Cruz F, Rocha EPC. Evolution of Conjugation and Type IV Secretion Systems. Mol Biol Evol. 2013;30: 315–331. doi:10.1093/MOLBEV/MSS221

46. Mitchell HM, Rocha GA, Kaakoush NO, O’Rourke JL, Queiroz DMM. The Family Helicobacteraceae. In: Rosenberg E, DeLong EF, Lory S, Stackebrandt E, Thompson F, editors. The Prokaryotes. Berlin, Heidelberg: Springer Berlin Heidelberg; 2014. pp. 337–392. doi:10.1007/978-3-642-39044-9_275

47. Mazzamurro FI, Baby Chirakadavil J, Durieux I, Poiré L, Plantade J, Ginevra C, et al. Intragenomic conflicts with plasmids and chromosomal mobile genetic elements drive the evolution of natural transformation within species. de Visser JAGM, editor. PLoS Biol. 2024;22: e3002814. doi:10.1371/JOURNAL.PBIO.3002814

48. Cury J, Touchon M, Rocha EPC. Integrative and conjugative elements and their hosts: composition, distribution and organization. Nucleic Acids Res. 2017;45: 8943–8956. doi:10.1093/nar/gkx607

49. Damke PP, Di Guilmi AM, Fernández Varela P, Velours C, Marsin S, Veaute X, et al. Identification of the periplasmic DNA receptor for natural transformation of Helicobacter pylori. Nat Commun. 2019;10: 1–11. doi:10.1038/s41467-019-13352-6

50. Hofreuter D, Odenbreit S, Henke G, Haas R. Natural competence for DNA transformation in Helicobacter pylori: identification and genetic characterization of the comB locus. Mol Microbiol. 1998;28: 1027–1038. doi:10.1046/J.1365-2958.1998.00879.X

51. Costa TRD, Patkowski JB, Macé K, Christie PJ, Waksman G. Structural and functional diversity of type IV secretion systems. Nature Reviews Microbiology 2023 22:3. 2023;22: 170–185. doi:10.1038/S41579-023-00974-3

52. Noureen M, Kawashima T, Arita M. Genetic Markers of Genome Rearrangements in Helicobacter pylori. Microorganisms. 2021;9: 621. doi:10.3390/microorganisms9030621

53. Denise R, Abby SS, Rocha EPC. Diversification of the type IV filament superfamily into machines for adhesion, protein secretion, DNA uptake, and motility. PLoS Biol. 2019;17: e3000390. doi:10.1371/JOURNAL.PBIO.3000390

54. Wen Y, Marcus EA, Matrubutham U, Gleeson MA, Scott DR, Sachs G. Acid-adaptive genes of Helicobacter pylori. Infect Immun. 2003;71: 5921–5939. doi:10.1128/IAI.71.10.5921-5939.2003;WGROUP:STRING:PUBLICATION

55. Bury-Moné S, Thiberge JM, Contreras M, Maitournam A, Labigne A, De Reuse H. Responsiveness to activity via metal ion regulators mediates virulense in the gastric pathogen Helicobacter pylori. Mol Microbiol. 2004;53: 623–638. doi:10.1111/J.1365-2958.2004.04137.X;WGROUP:STRING:PUBLICATION

56. Moore ME, Lam A, Bhatnagar S, Solnick J V. Environmental Determinants of Transformation Efficiency in Helicobacter pylori. J Bacteriol. 2014;196: 337–344. doi:10.1128/JB.00633-13

57. Krüger NJ, Knüver MT, Zawilak-Pawlik A, Appel B, Stingl K. Genetic Diversity as Consequence of a Microaerobic and Neutrophilic Lifestyle. PLoS Pathog. 2016;12: 1–24. doi:10.1371/journal.ppat.1005626

58. Corbinais C, Mathieu A, Damke PP, Kortulewski T, Busso D, Prado-Acosta M, et al. ComB proteins expression levels determine Helicobacter pylori competence capacity. 2017. doi:10.1038/srep41495

59. Kavermann H, Burns BP, Angermuller K, Odenbreit S, Fischer W, Melchers K, et al. Identification and characterization of Helicobacter pylori genes essential for gastric colonization. J Exp Med. 2003;197: 813–822. doi:10.1084/jem.20021531

60. Porwollik S, Noonan B, O’Toole PW. Molecular characterization of a flagellar export locus of Helicobacter pylori. Infect Immun. 1999;67: 2060–2070. doi:10.1128/IAI.67.5.2060-2070.1999;PAGE:STRING:ARTICLE/CHAPTER

61. Hospenthal MK, Costa TRD, Waksman G. A comprehensive guide to pilus biogenesis in Gram-negative bacteria. Nature Reviews Microbiology. 2017. doi:10.1038/nrmicro.2017.40

62. Goldlust K, Ducret A, Halte M, Dedieu-Berne A, Erhardt M, Lesterlin C. The F pilus serves as a conduit for the DNA during conjugation between physically distant bacteria. Proc Natl Acad Sci U S A. 2023;120. doi:10.1073/PNAS.2310842120/-/DCSUPPLEMENTAL

63. Clarke M, Maddera L, Harris RL, Silverman PM. F-pili dynamics by live-cell imaging. Proc Natl Acad Sci U S A. 2008;105. doi:10.1073/pnas.0806786105

64. Ripoll-Rozada J, Zunzunegui S, de la Cruz F, Arechaga I, Cabezón E. Functional Interactions of VirB11 Traffic ATPases with VirB4 and VirD4 Molecular Motors in Type IV Secretion Systems. J Bacteriol. 2013;195: 4195–4201. doi:10.1128/JB.00437-13/ASSET/C2933D45-2055-4F32-967C-D87FC83821B7/ASSETS/GRAPHIC/ZJB9990927860005.JPEG

65. Jeong JY, Yim HS, Ryu JY, Lee HS, Lee JH, Seen DS, et al. One-Step Sequence- and Ligation-Independent Cloning as a Rapid and Versatile Cloning Method for Functional Genomics Studies. Appl Environ Microbiol. 2012;78: 5440–5443. doi:10.1128/AEM.00844-12

66. Tomb JF, White O, Kerlavage AR, Clayton RA, Sutton GG, Fleischmann RD, et al. The complete genome sequence of the gastric pathogen Helicobacter pylori. Nature. 1997;388: 539–547. doi:10.1038/41483

67. Liu H, Naismith JH. An efficient one-step site-directed deletion, insertion, single and multiple-site plasmid mutagenesis protocol. BMC Biotechnology 2008 8:1. 2008;8: 91-. doi:10.1186/1472-6750-8-91

68. Grote A, Hiller K, Scheer M, Münch R, Nörtemann B, Hempel DC, et al. JCat: a novel tool to adapt codon usage of a target gene to its potential expression host. Nucleic Acids Res. 2005;33: W526– W531. doi:10.1093/nar/gki376

69. Meng EC, Goddard TD, Pettersen EF, Couch GS, Pearson ZJ, Morris JH, et al. UCSF ChimeraX: Tools for structure building and analysis. Protein Science. 2023;32: e4792. doi:10.1002/pro.4792

70. Eddy SR. Accelerated Profile HMM Searches. PLoS Comput Biol. 2011;7. doi:10.1371/journal.pcbi.1002195

