## Supplemental Figures 1 to 8 for "Unveiling a missing component of the atypical type IV secretion system required for natural transformation of *Helicobacter pylori*"

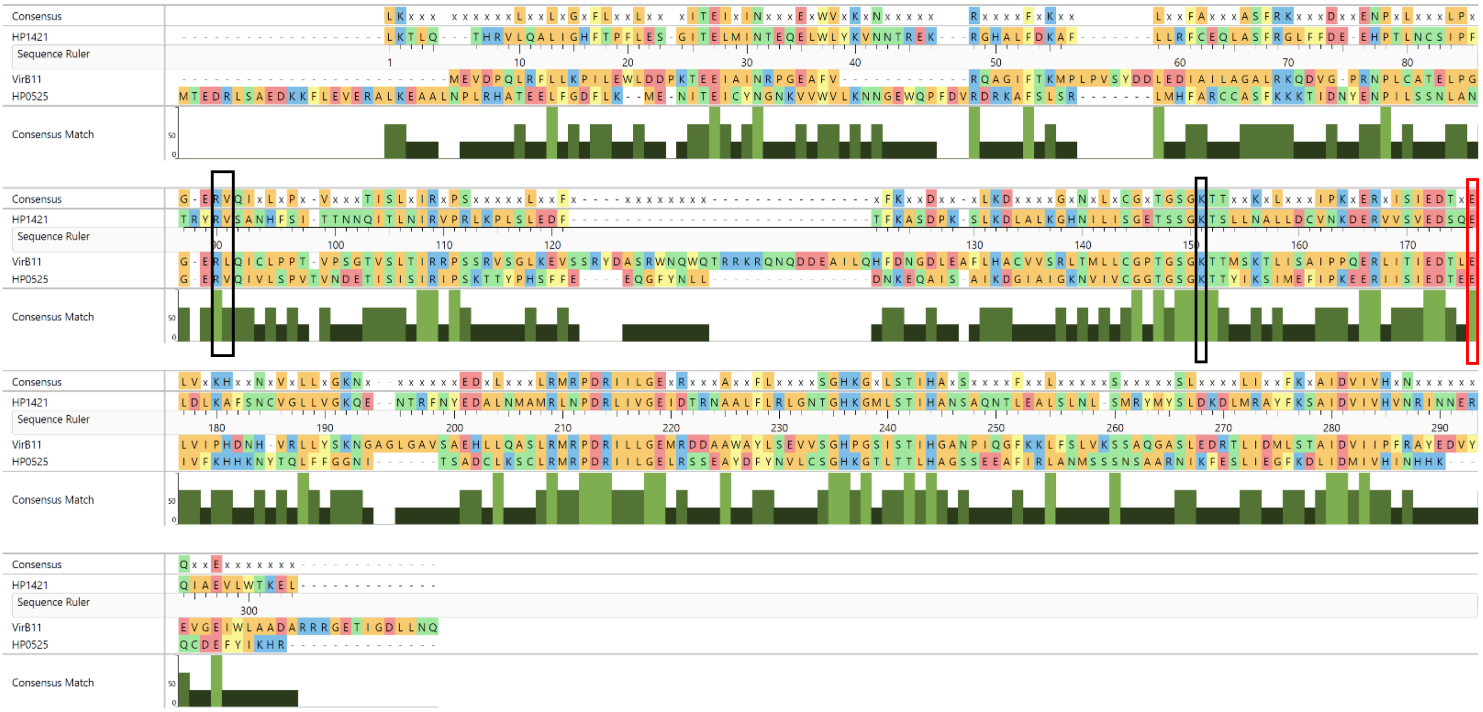


**S1 Figure.** Sequence alignment of the VirB11 ATPases in *H. pylori* HP1421. The following sequences were used in the alignment: VirB11 from Ti plasmid pTiBo542 *Agrobacterium tumefaciens,* HP0525 and HP1421 both from *H. pylori*. Conserved residues also found in other ATPases are shown in black boxes, residues involved in ATP binding and hydrolysis are boxed in red. Amino acid numbering at the sequence ruler refers to HP1421.

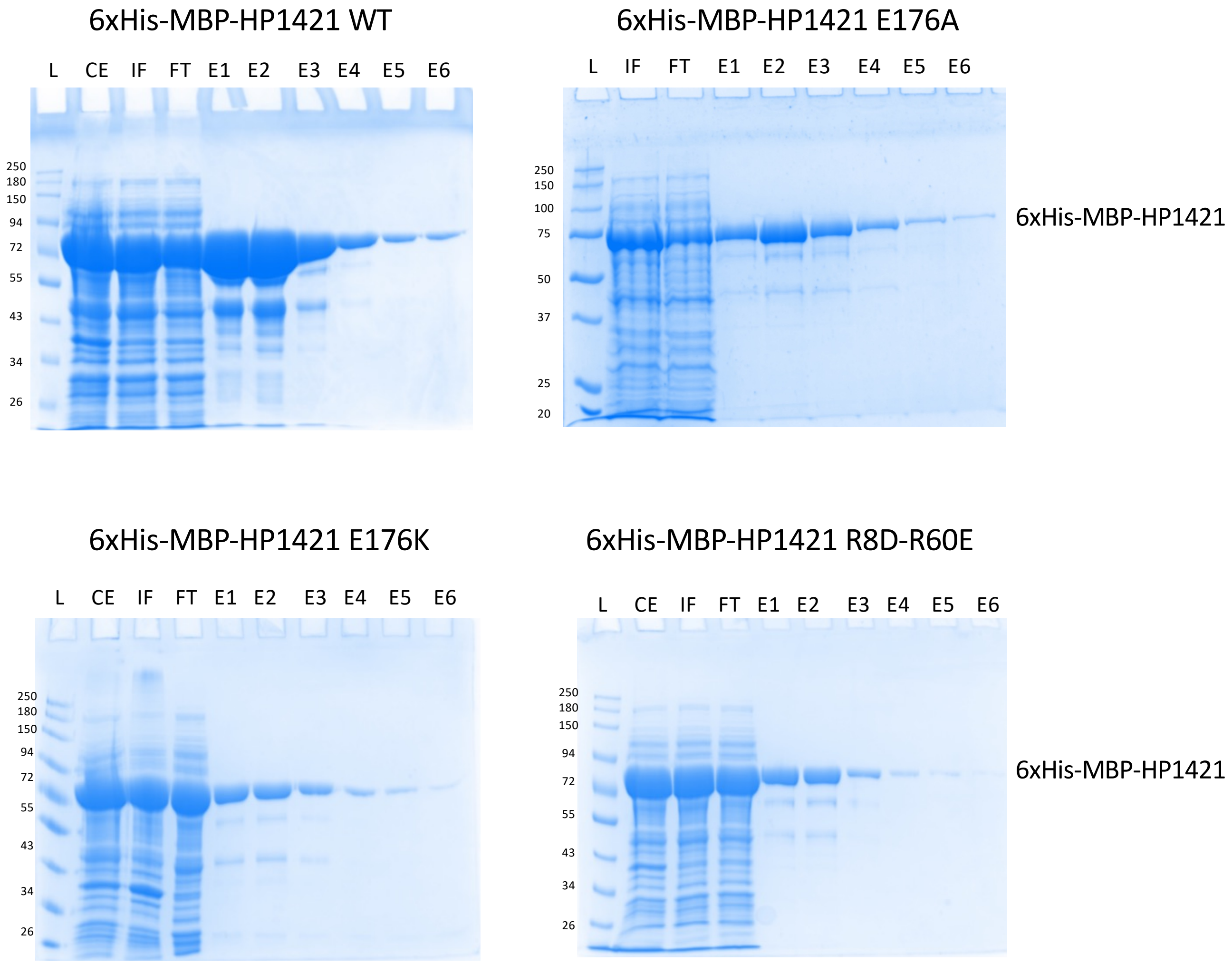


**S2A Figure.** HP1421 purification. 12,5 % acrylamide gels stained by Coomassie blue. L:ladder; CE: crude extract; IF: input fraction; FT: flowthrough; E1-6: elution fractions.

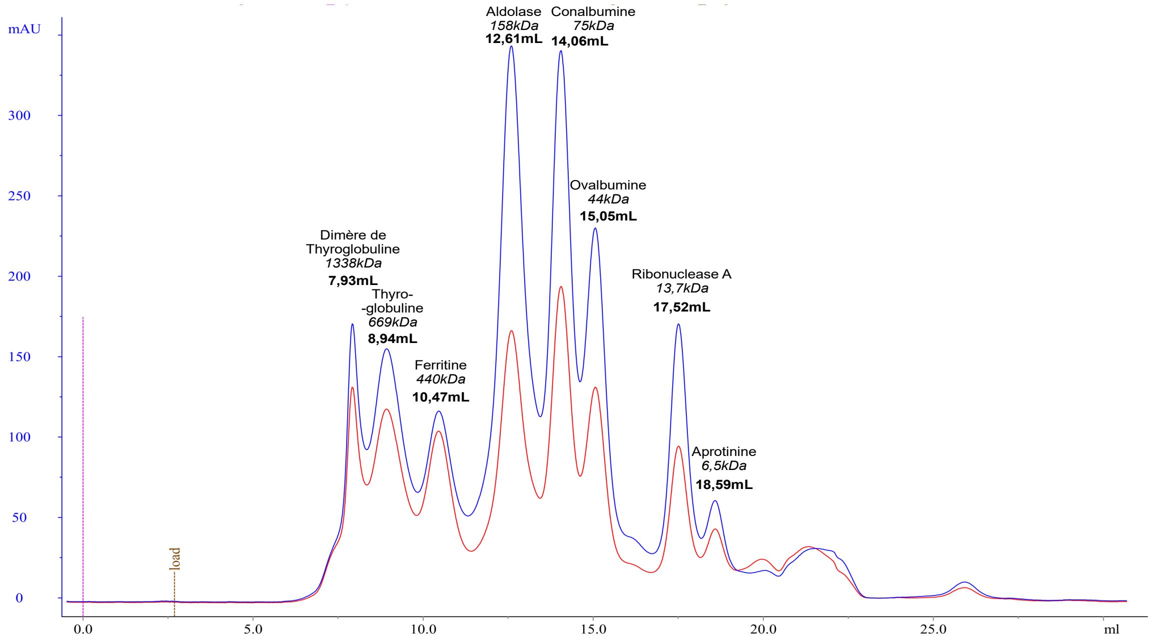


**S2B Figure.** Chromatogram for size exclusion chromatography (SEC) of protein standards using a Superdex 200 column.

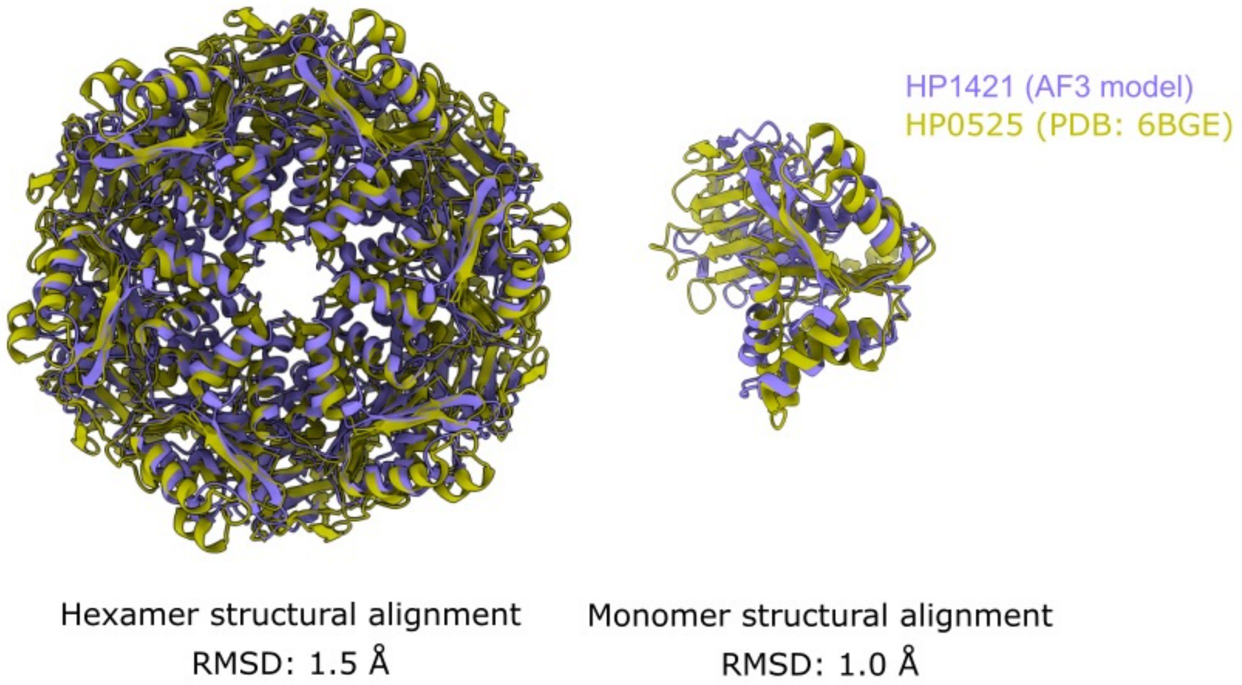


**S3A Figure.** HP1421 hexameric model superimposed with HP0525 experimental structure. Model from AlphaFold3 using 6 copies of HP1421. HP0525 structure from Protein Data Bank (6GBE) a) hexamer structural alignment (RMSD: 1.5 Å). b) monomer structural alignment (RMSD: 1.0 Å).

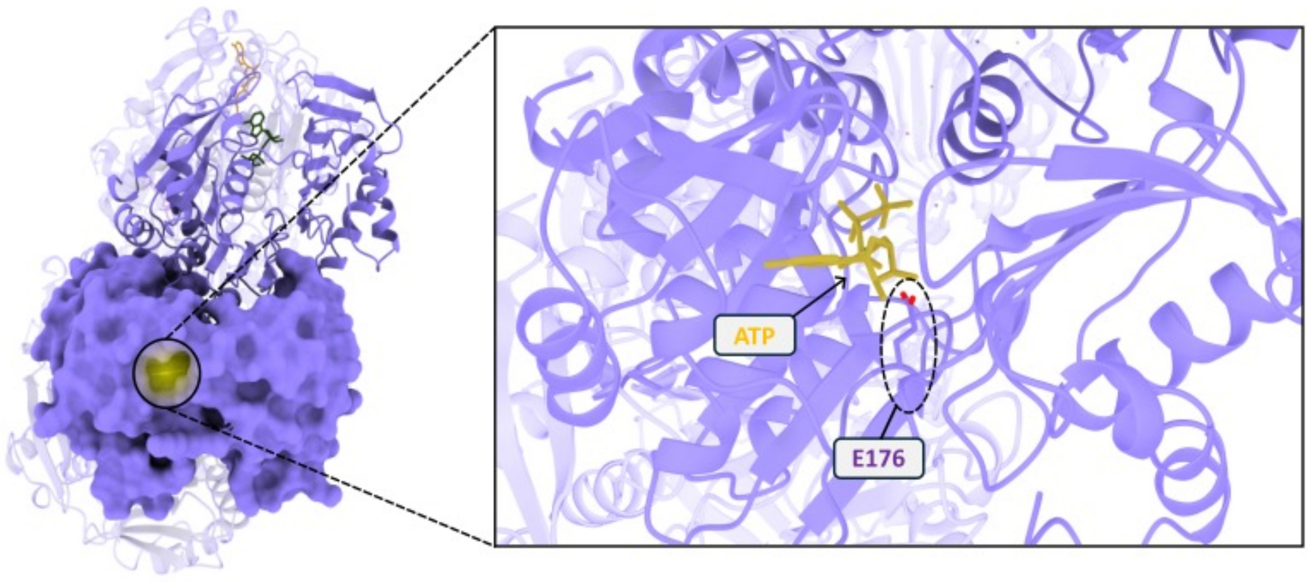


**S3B Figure.** Modelling of ATP binding in HP1421 pocket in hexameric context. AlphaFold3 model of HP1421 with 6 ATP all confidently predicted to bind into a HP1421 pocket, in contact with E176 residue (chain-pair ipTM between HP1421 and the ATP molecule bound to its surface = 0.87 to 0.89).

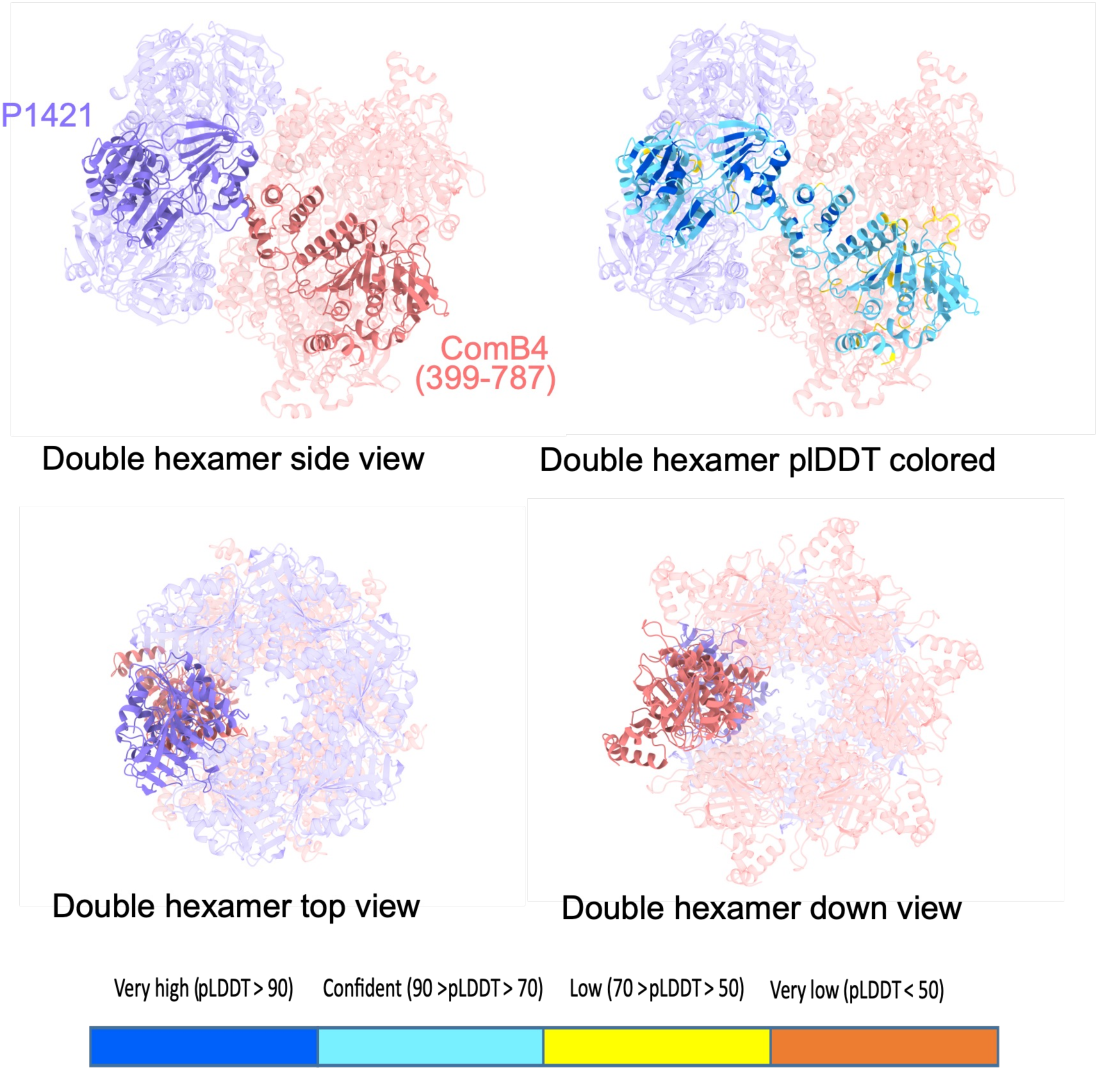


**S4A Figure. AlphaFold3 model of HP1421 and ComB4 double hexamer in different views**. HP1421 (blue) and ComB4 C-terminal domain (salmon) double hexamer prediction (pTM = 0.82, ipTM = 0.81) with one copy of each highlighted to ease visualisation. Panels represents double hexamer model A) side view, B) side view with a plDDT colored HP1421/ComB4 complex, C) top view, D) down view.

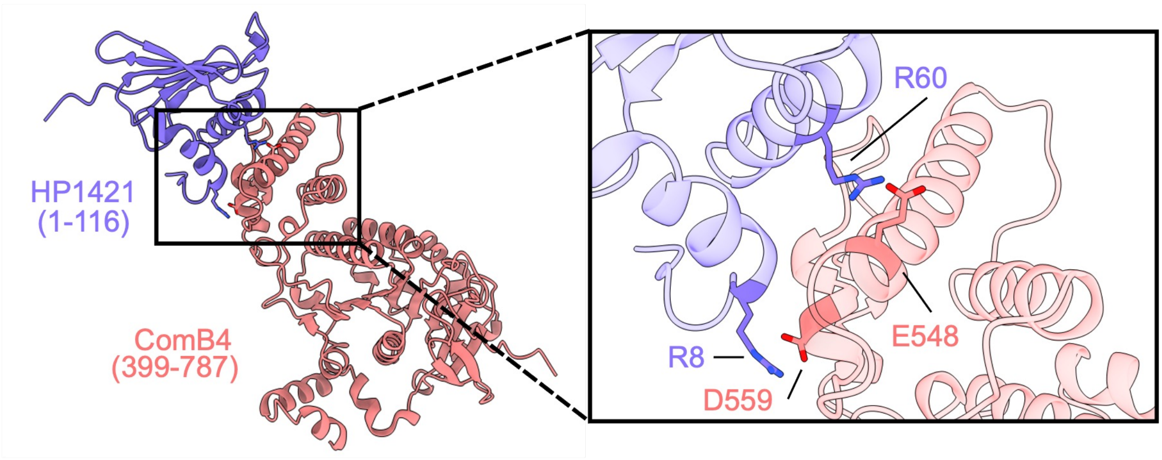


**S4B Figure. AlphaFold 2 model of HP1421 and ComB4**. HP1421 N-ter (1-116) and ComB4 C-ter (399-787) AF2 model (pTM = 0.89, ipTM = 0.88). B) Zoom on HP1421 and Comb4 predicted interface and main evolutionary conserved residues (R8 and R60 on HP1421, E548 and D559 on ComB4).

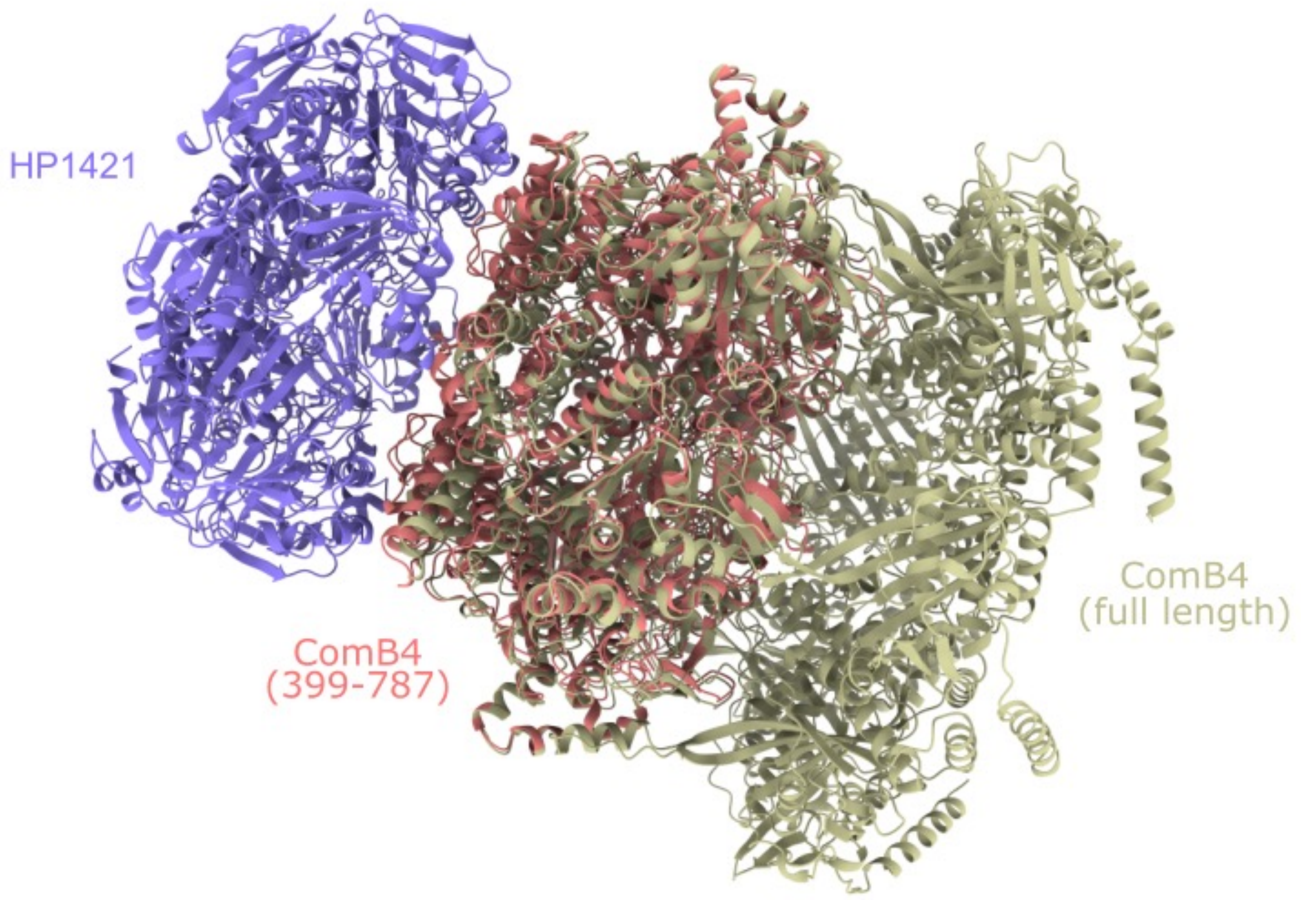


**S5 Figure. HP1421 and ComB4 double hexamer structurally aligned with full length ComB4 hexamer.** AlphaFold3 models of HP1421 (full length) and ComB4 (399-787) double hexamer and full length ComB4 hexamer.

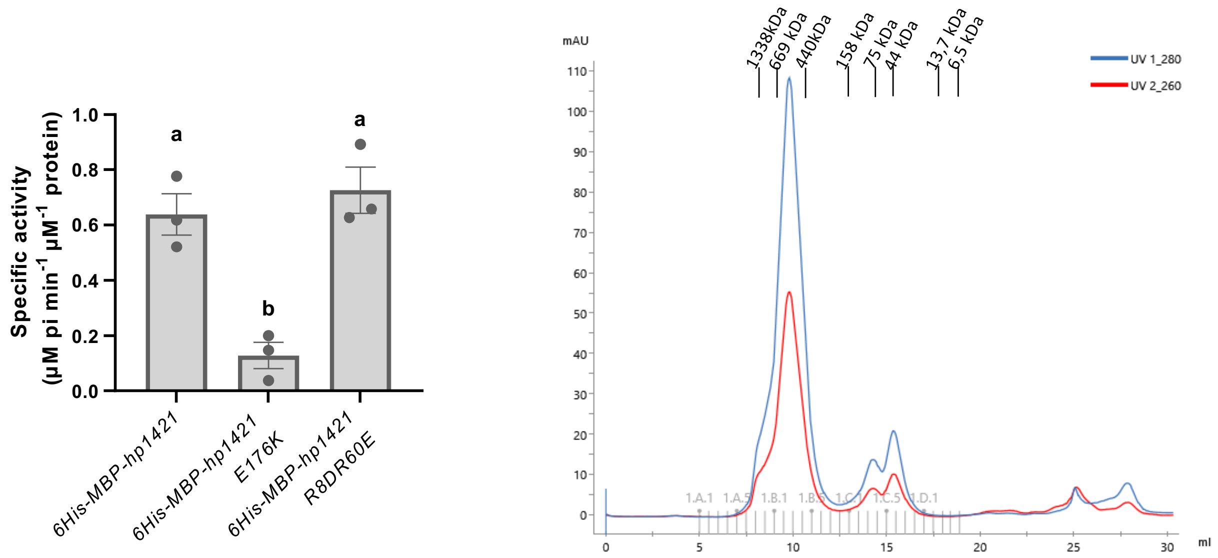


**S6 Figure. A)** Specific ATPase activity of purified HP1421 variants was measured by following Pi release due to ATP hydrolysis, data represent Mean + SEM (n=3). Different lowercase letters indicate significant differences ($p<0.05$) between treatments (Tukey's test). **B)** Chromatogram for size exclusion chromatography (SEC) of 6xHis-MBP-HP1421 R8D-R60E using a Superdex 200 column.

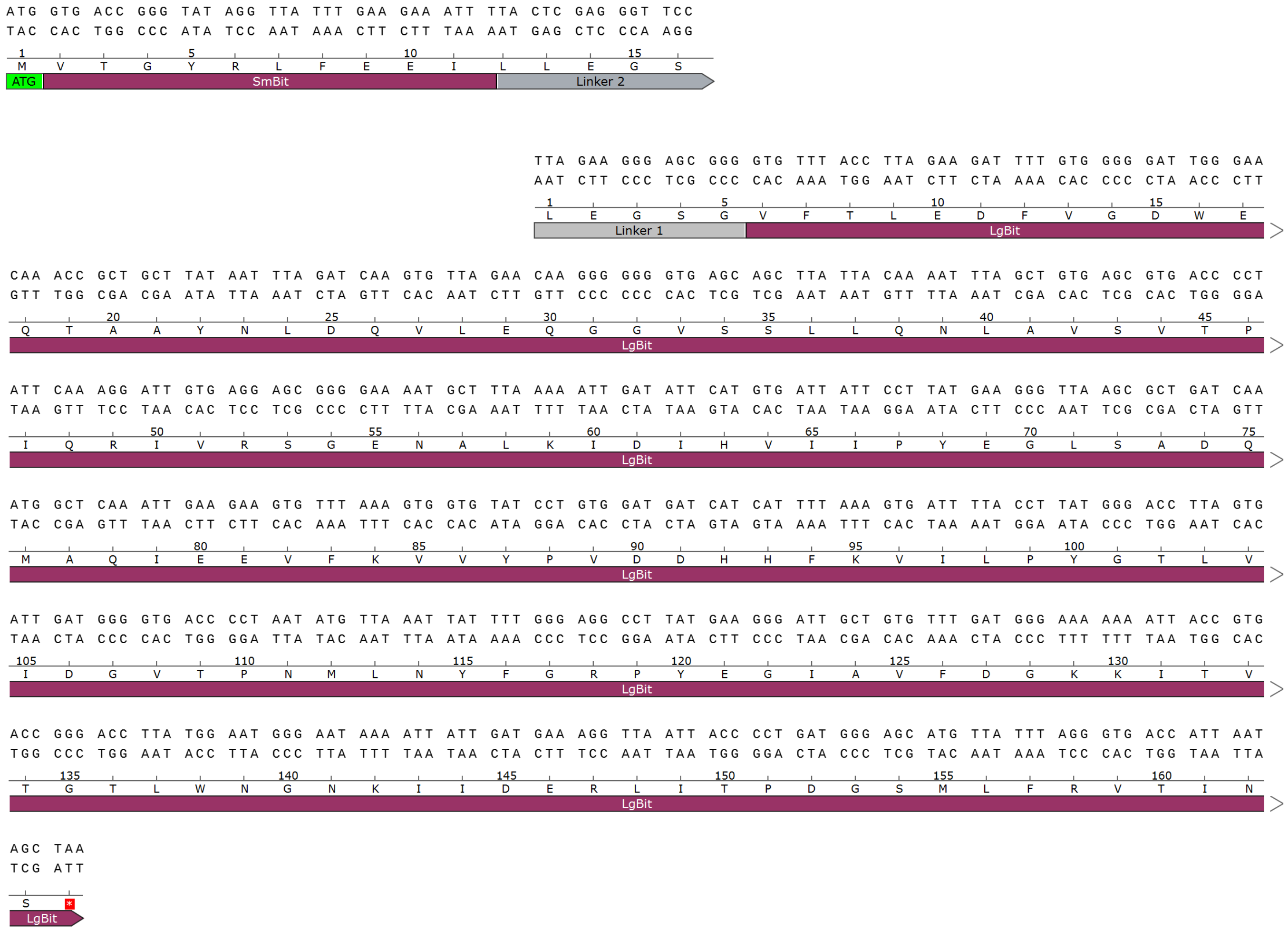


**S7 Figure.** Codon optimized sequences of SmBit for N terminal tagging and LgBit for Cterminal tagging of Nanoluciferase for *H. pylori* expression.

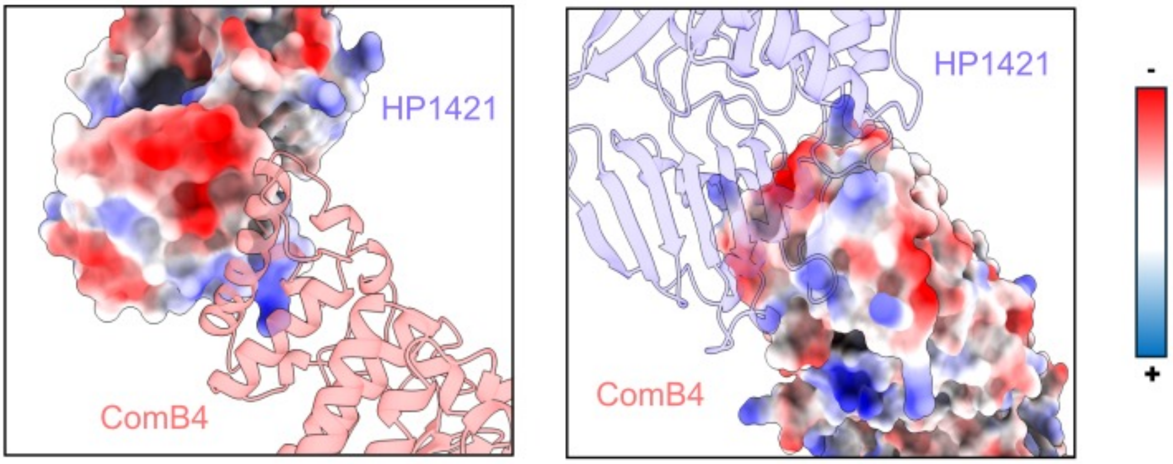


**S8 Figure. Complementary surface electrostatics at HP1421 and ComB4 interface.** AlphaFold3 model of HP1421 and ComB4 zoom on each protein interface area with electrostatic potential shown.
