## Supplemental Tables 1 to 4 for "Unveiling a missing component of the atypical type IV secretion system required for natural transformation of *Helicobacter pylori*"

**Supplementary Table 1**. *H. pylori* strains

| Strain | Genotype | Source |
|---|---|---|
| 1 | 26695 | (1) |
| 134 | 26695 *strep*$^{R}$ | (1) |
| 770 | 26695 *comB2::Km* | (3) |
| 1188, 1189 | 26695 *hp1421::Kan* | This work |
| 1283, 1284 | 26695 *hp1421::Kan pUreA-FLAG-hp1421-Cm* | This work |
| 1303, 1304 | 26695 *hp1421::Kan pUreA-FLAG-hp1421 E176A-Cm* | This work |
| 1376, 1377 | 26695 *hp1421::Kan pUreA-FLAG-hp1421 E176K-Cm* | This work |
| 1378,1399 | 26695 *hp1421::Kan pUreA-FLAG-hp1421 E176K-Cm* comB4::*Apra* | This work |
| 1397 | 26695 comB4::*Apra* | This work |
| 1494, 1495 | 26695 *hp1421::Kan rdxA-pcomH-FLAG-hp1421-Cm* | This work |
| 1630, 1631 | 26695 *hp1421::Kan* comB4::*Apra pUreA-comB4 E548R/D559R-AT-pcomH-FLAG-hp1421-CmR* | This work |
| 1632, 1633 | 26695 *hp1421::Kan* comB4::*Apra pUreA-comB4-AT-pcomH-FLAG-hp1421-CmR* | This work |
| 1634, 1635 | 26695 *hp1421::Kan* comB4::*Apra pUreA-comB4-AT-pcomH-FLAG-hp1421 R8D/R60E-CmR* | This work |
| 1642 | 26695 *pUreA-comB4-SmBit-pcomH-LgBit-hp1421-CmR* | This work |
| 1643 | 26695 *pUreA-comB4-SmBit-pcomH-LgBit-hp1421R8D/R60E-CmR* | This work |
| 1665 | 26695 *pUreA-comB4 E548R/D559R-SmBit-pcomH-LgBit-hp1421-CmR* | This work |

1. Marsin S, Mathieu A, Kortulewski T, Guérois R, Radicella JP. Unveiling novel RecO distant orthologues involved in homologous recombination. PLoS Genet. 2008;4(8).

2.Mathieu A, O'Rourke EJ, Radicella JP. *Helicobacter pylori* genes involved in avoidance of mutations induced by 8-oxoguanine. J bacteriol. 2006; 188(21), 7464–7469.

3. Corbinais C, Mathieu A, Kortulewski T, Radicella JP, Marsin S. Following transforming DNA in Helicobacter pylori from uptake to expression. Mol Microbiol. 2016;101(6):1039– 53

**Supplementary table 2**. *E. coli* strains

| Strain | Description | Source |
|---|---|---|
| DH5-α | For plasmid construction and maintenance | New England Biolabs |
| BL21 (DE3) | For protein expression and purification | New England Biolabs |
| BTH101 | For bacterial two-hybrid tests | Euromedex |

**Supplementary Table 3**. Plasmids used in this study

| Plasmid | Genotype | Source |
|---|---|---|
| p1361 | pBAD33 pEYY166 Dronpa-MTSBs | Lab. collection |
| p1524, p1980 | pJET 1.2 *hp1421::Kan* | This work |
| p1561 | pKT25 T25-Zip | Euromedex |
| p1562 | pUT18C *T18-Zip* | Euromedex |
| p1665 | pUT18C *T18-hp1421-Amp* | This work |
| p1765 | pRSF Duet *6xHis-MBP-TEV-hp1421-Kan* | Laurent Terradot |
| p1770 | pKT25 *T25-hp1421-Kan* | This work |

| | | |
|---|---|---|
| p1781 | pJET 1.2 *pUreA-FLAG-hp1421-Cm* | This work |
| p1787 | pJET 1.2 *pUreA-FLAG-hp1421 E176A-Cm* | This work |
| p1792 | pRSF Duet *6xHis-MBP-TEV-hp1421 E176A-Kan* | This work |
| p1811 | pRSF Duet *6xHis-MBP-TEV-hp1421 E176K-Kan* | This work |
| p1813 | pJET 1.2 *pUreA-FLAG-hp1421 E176K-Cm* | This work |
| p1820 | pJET 1.2 *comB4::Apra* | This work |
| p1854 | pJET 1.2 *rdxA-pcomH-FLAG-hp1421-Cm* | This work |
| p1916 | pRSF Duet *6xHis-MBP-TEV-hp1421 R8D/R60E-Kan* | This work |
| p1946 | pJET 1.2 *pUreA-comB4-AT-pcomH-FLAG-hp1421-CmR* | This work |
| p1947 | pJET 1.2 *pUreA-comB4-AT-pcomH-FLAG-hp1421 R8D/R60E-CmR* | This work |
| p1948 | pJET 1.2 *pUreA-comB4 E548R/D559R-AT-pcomH-FLAG-hp1421-CmR* | This work |
| p1961 | pJET 1.2 *pUreA-comB4-SmBit-pcomH-LgBit-hp1421R8D/R60E-CmR* | This work |
| p1964 | pJET 1.2 *pUreA-comB4 E548R/D559R-SmBit-pcomH-LgBit-hp1421-CmR* | This work |
| p1968 | pJET 1.2 *pUreA-comB4-SmBit-pcomH-LgBit-hp1421-CmR* | This work |
| p1983 | pUT18C *T18-hp1421 R8D/R60E-Amp* | This work |
| p1984 | pKT25 T25-hp1421 *R8D/R60E-Kan* | This work |
| p1985 | pUT18 - negative control for BTH | Euromedex |
| p1986 | pKT25 - negative control for BTH | Euromedex |

**Supplementary table 4.** Oligonucleotides used in this study

| Name | Sequence (5'-3') | Description |
|---|---|---|
| Oc61 | AGACAGCCAAAAATTGGATTTAAAAGC | *Construction p1811 and p1813* |
| Op31 | ATGGAATTGAATCAACCACCACTC | Fw HP0247 (DNA uptake, Recipient) |
| Op32 | TTAACGGCGTTTGGGTTTTTTAG | Rv HP0247 (DNA uptake, Recipient) |
| Oc139 | GAATGTTAGAAAAGCTTTTAAGGGTACCCGGGTGACTAAC | *Construction p1820* |
| Oc140 | CTAGCTCCTCAAAAGTTTCTGGATCCCCGTGTCATTATTC | *Construction p1820* |
| Oc141 | GAATAATGACACGGGGATCCAGAAACTTTTGAGGAGCTAG | *Construction p1820* |
| Oc142 | GTTAGTCACCCGGGTACCCTTAAAAGCTTTTCTAACATTC | *Construction p1820* |
| Oc266 | GAATAAGCATGGATTACAAGGATGACGACGATAAGTTGAAAACTTTAC | *Construction p1854* |
| Oc267 | CCTTGTAATCCATGCTTATTCCTTTTTATTTTAGATTTTTGGATTATAGC | *Construction p1854* |
| Oc306 | ATGACGTTTTACAAGCCCTAATTGGCCATTTTAC | *SDM R8D hp1421 (p1983 and 1984)* |
| Oc307 | CTTGTAAAACGTCATGGGTTTGTAAAGTTTTC | *SDM R8D hp1421 (p1983 and 1984)* |
| Oc308 | TTTGCTGGAGTTTTGCGAGCAATTGGCTAG | *SDM R60E hp1421(p1983 and 1984)* |
| Oc309 | AACTCCAGCAAAAACGCCTTATCAAAAAGTG | *SDM R60E hp1421 (p1983 and 1984)* |
| Oc345 | CTTTATTTTCAGGGCGCCATGGGAAAAACTTTACAAACCCAT | *Construction p1916* |

| | | |
|---|---|---|
| Oc346 | CCGAGCTCGAATTCGGATCCTTAAAGCTCTTTAGTCCATAAG | *Construction p1916* |
| Oc347 | CTTATGGACTAAAGAGCTTTAAGGATCCGAATTCGAGCTCGG | *Construction p1916* |
| Oc348 | ATGGGTTTGTAAAGTTTTTCCCATGGCGCCCTGAAAATAAAG | *Construction p1916* |
| Oc429 | TAGCTCGAGGGTTCCGGACACTTCTTTTCAAATCCCACAAC | *Construction p1946, p1947 and p1948* |
| Oc430 | GGGGTGAGTTTCATGGATCCAGCGGATCCTTAC | *Construction p1946, p1947 and p1948* |
| Oc431 | TAAGGATCCGCTGGATCCATGAAACTCACCCC | *Construction p1946, p1947 and p1948* |
| Oc432 | GTGTCCGGAACCCTCGAGCTATTCGGTCAGGCGG | *Construction p1946, p1947 and p1948* |
| Oc454 | GCTCCTCAAAAGTTTCTCG | *Construction p1961, p1964 and p1968* |
| Oc455 | GCGAGAAACTTTTGAGGAGCCTCGAGGGTTCCGGA | *Construction p1961, p1964 and p1968* |
| Oc456 | GGGATTTGAAAAGAAGTGTTATAAAATTTCTTCAAATAACCTATACCC | *Construction p1961, p1964 and p1968* |
| Oc482 | TACTGGATGAATTGTTTTAGATGCCCCTAAAATCCTTAAAAAAC | *Construction p1980* |
| Oc483 | GATATTCTCATTTTAGCCATAAGCCCTATCCTTTTATCATC | *Construction p1980* |
| Op99 | CACTTCTTTTCAAATCCCACAACC | *Construction p1961, p1964 and p1968* |
| Op284 | GGACACCCGTTCGCGATTG | Fw p1361 (DNA uptake, Donor) |
| Op285 | CGTCAGGATGGCCTTCTGC | Rv p1361 (DNA uptake, Donor) |
| Op376 | GATAAAAGGATAGGGCTTTTGAAAACTTTACAAACCCAATGGCTAAAATGAGAATATC | *Construction p1524* |
| Op377 | GGATTTTAGGGGCATTTAAAGCTCTTTAGTCCATACTAAAACAATTCATCCAG | *Construction p1524* |
| Op378 | CTGGATGAATTGTTTTAGTATGGACTAAAGAGCTTTAAATGCCCCTAAAATCC | *Construction p1524* |
| Op379 | GATATTCTCATTTTAGCCATTGGGTTTGTAAAGTTTTCAAAAGCCCTATCCTTTTATC | *Construction p1524* |
| Op722 | GGTCGACTCTAGAGGATCCCCGGGTACCTAAGTTGAAAACTTTACAAACCCATAGAG | *Construction p1770* |
| Op723 | CGTTGTAAAACGACGGCCGAATTCTTAGTTAAAGCTCTTTAGTCCATAAGACTTCAGC | *Construction p1770* |
| Op724 | CTCTATGGGTTTGTAAAGTTTTCAACTTAGGTACCCGGGGATCCTCTAGAGTCGACC | *Construction p1770* |
| Op725 | GCTGAAGTCTTATGGACTAAAGAGCTTTAACTAAGAATTCGGCCGTCGTTTTACAACG | *Construction p1770* |
| Op726 | GGGTACCGAGCTCGAATTCATCGATATTGAAAACTTTACAAACCCATAGAG | *Construction p1665* |
| Op727 | ATTGTACTGAGAGTGCACCATATTACTTAGTTAAAGCTCTTTAGTCCATAAGACTTCAGC | *Construction p1665* |
| Op728 | CTCTATGGGTTTGTAAAGTTTTCAATATCGATGAATTCGAGCTCGGTACCC | *Construction p1665* |

| | | |
|---|---|---|
| Op729 | GCTGAAGTCTTATGGACTAAAGAGCTTTAACTAAGTAA TATGGTGCACTCTCAGTACAAT | *Construction p1665* |
| Op940 | AGACAGCCAAGCATTGGATTTAAAAGC | *Construction p1787 and 1792* |
| Op941 | TCAACGCTCACCACCCTT | *Construction p1787, p1792, p1811 and p1813* |
| Op1007 | GAATAGGAGAATAAGGAATTCATGGATTACAAGGATG ACGACGATAAGTTGAAAACTTTAC | *Construction p1781* |
| Op1008 | GATCTAGAGTCGCGGCCGCTTTAAAGCTCTTTAGT | *Construction p1781* |
| Op1009 | GTAAAGTTTTCAACTTATCGTCGTCATCCTTGTAATCCAT GAATTCCTTATTCTCCTATTC | *Construction p1781* |
| Op1010 | ACTAAAGAGCTTTAAAGCGGCCGCGACTCTAGATC | *Construction p1781* |
| SeqAM15 | CTAAAACAATTCATCCAGTAAAATATA | Rv KanR |
| SeqAM16 | ATGGCTAAAATGAGAATATCACCGGA | *Fw KanR* |
