## Supplemental Table 6 for "Unveiling a missing component of the atypical type IV secretion system required for natural transformation of *Helicobacter pylori*"

```
HMMER3/b [3.0 | March 2010]
NAME  T_virB11
LENG  328
ALPH  amino
RF    no
CS    no
MAP   yes
DATE  Fri Nov 23 17:23:46 2012
NSEQ  73
EFFN  0.717346
CKSUM 1137158280
STATS LOCAL MSV      -11.1088  0.70062
STATS LOCAL VITERBI  -12.1113  0.70062
STATS LOCAL FORWARD   -5.7466  0.70062
HMM          A        C        D        E        F        G        H
I        K        L        M        N        P        Q        R
S        T        V        W        Y
            m->m     m->i     m->d     i->m     i->i     d->m     d-
>d
  COMPO   2.38854  4.45760  3.00357  2.66473  3.62695  2.78256
3.78549  2.72577  2.81836  2.33338  3.62017  3.19176  3.30926
3.19909  2.77998  2.73579  2.80516  2.48540  4.88696  3.88056
          2.68605  4.42214  2.77536  2.73130  3.46370  2.40518
3.72500  3.29345  2.67757  2.69362  4.24518  2.90355  2.73730
3.18147  2.89807  2.37881  2.77514  2.98519  4.58456  3.61519
          0.32636  1.65496  2.43781  1.51339  0.24867  0.00000
*
      1   2.45979  4.25749  3.43492  2.87307  3.39806  3.42384
3.72736  2.70828  2.78968  2.44639  2.33460  3.26861  3.51311
3.11687  3.05828  2.76998  2.68683  2.53016  4.84660  3.42289     12
- -
          2.68618  4.42225  2.77519  2.73123  3.46354  2.40513
3.72494  3.29354  2.67741  2.69355  4.24690  2.90347  2.73739
3.18146  2.89801  2.37887  2.77519  2.98518  4.58477  3.61503
          0.02930  3.94069  4.66304  0.61958  0.77255  0.53548
0.88042
      2   2.19468  4.67140  2.94043  2.34518  3.91022  3.34654
3.62500  3.31313  2.44382  2.94094  3.59797  2.96094  3.64792
2.75658  2.76149  2.37300  2.73353  2.86974  5.20868  3.89405     13
- -
          2.68618  4.42225  2.77519  2.73123  3.46354  2.40513
3.72494  3.29354  2.67741  2.69355  4.24690  2.90347  2.73739
3.18146  2.89801  2.37887  2.77519  2.98518  4.58477  3.61503
          0.02873  3.95996  4.68231  0.61958  0.77255  0.49457
0.94118
      3   2.41422  4.50682  3.06877  2.58883  3.69869  3.12546
3.68133  2.93284  2.48926  2.73072  3.35994  3.00288  3.24885
2.90144  2.98887  2.60869  2.57220  2.58746  5.06609  3.78593     14
- -
          2.68618  4.42225  2.77519  2.73123  3.46354  2.40513
3.72494  3.29354  2.67741  2.69355  4.24690  2.90347  2.73739
3.18146  2.89801  2.37887  2.77519  2.98518  4.58477  3.61503
          0.02777  3.99342  4.71576  0.61958  0.77255  0.49800
0.93584
```

```
      4   2.43197  4.78481  2.88003  2.35148  3.96592  3.33942
3.41067  3.26806  2.32745  3.00229  3.75035  2.81762  3.78571
2.67633  2.70481  2.33925  2.64533  3.07359  5.28361  3.94580      15
- -
          2.68618  4.42225  2.77519  2.73123  3.46354  2.40513
3.72494  3.29354  2.67741  2.69355  4.24690  2.90347  2.73739
3.18146  2.89801  2.37887  2.77519  2.98518  4.58477  3.61503
          0.02747  4.00411  4.72645  0.61958  0.77255  0.50504
0.92504
      5   2.39738  4.72655  2.92045  2.43477  3.91162  3.40180
3.28308  3.37130  2.42386  2.92624  3.80765  2.90855  3.17509
2.68387  2.85589  2.29522  2.63669  3.03421  5.02030  3.91686      16
- -
          2.68618  4.42225  2.77519  2.73123  3.46354  2.40513
3.72494  3.29354  2.67741  2.69355  4.24690  2.90347  2.73739
3.18146  2.89801  2.37887  2.77519  2.98518  4.58477  3.61503
          0.02747  4.00411  4.72645  0.61958  0.77255  0.50504
0.92504
      6   2.37019  4.46214  3.17414  2.63381  3.42813  3.46772
3.18816  2.83950  2.60579  2.64040  3.43640  3.07998  3.74924
2.76417  2.99234  2.38882  2.69007  2.70540  4.83878  3.74650      17
- -
          2.68618  4.42225  2.77519  2.73123  3.46354  2.40513
3.72494  3.29354  2.67741  2.69355  4.24690  2.90347  2.73739
3.18146  2.89801  2.37887  2.77519  2.98518  4.58477  3.61503
          0.02747  4.00411  4.72645  0.61958  0.77255  0.50504
0.92504
      7   2.44673  4.91962  2.87597  2.30434  4.13380  3.38186
3.13884  3.55878  2.19267  3.19453  3.97963  2.71257  3.77515
2.53407  2.56593  2.44505  2.76587  3.23718  5.38654  4.02586      18
- -
          2.68618  4.42225  2.77519  2.73123  3.46354  2.40513
3.72494  3.29354  2.67741  2.69355  4.24690  2.90347  2.73739
3.18146  2.89801  2.37887  2.77519  2.98518  4.58477  3.61503
          0.02747  4.00411  4.72645  0.61958  0.77255  0.50504
0.92504
      8   2.25322  4.74160  2.86530  2.39814  3.93260  3.39314
3.54156  3.39737  2.42709  2.85527  3.82867  2.86928  3.28566
2.71114  2.84136  2.37832  2.62073  3.07975  5.26136  3.93292      19
- -
          2.68618  4.42225  2.77519  2.73123  3.46354  2.40513
3.72494  3.29354  2.67741  2.69355  4.24690  2.90347  2.73739
3.18146  2.89801  2.37887  2.77519  2.98518  4.58477  3.61503
          0.03176  4.00411  4.34130  0.61958  0.77255  0.50504
0.92504
      9   2.36165  4.83661  2.78685  1.95202  4.12369  3.11695
3.62540  3.45949  2.41250  3.10725  3.94736  2.89579  3.78880
2.73712  2.89579  2.53868  2.50256  3.20344  5.36623  4.01804      20
- -
          2.68618  4.42226  2.77520  2.73119  3.46355  2.40514
3.72495  3.29355  2.67742  2.69354  4.24691  2.90348  2.73740
3.18147  2.89802  2.37888  2.77514  2.98518  4.58478  3.61504
          0.10236  2.42585  4.72228  0.41148  1.08670  0.50226
0.92928
```

```
     10   1.88718  4.41031  3.21223  2.66423  3.88209  2.87190
3.80455  3.26591  2.69004  2.93689  3.65273  3.08328  3.78434
3.02436  3.05284  2.14306  2.58403  2.80524  5.23214  3.96199     22
- -
          2.68618  4.42225  2.77519  2.73123  3.46354  2.40513
3.72494  3.29354  2.67741  2.69355  4.24690  2.90347  2.73739
3.18146  2.89801  2.37887  2.77519  2.98518  4.58477  3.61503
          0.02747  4.00411  4.72645  0.61958  0.77255  0.50504
0.92504
     11   2.40756  4.35300  3.24942  2.73345  3.22081  3.50100
3.64750  2.77926  2.71146  2.26271  3.38910  3.16139  3.88305
2.93328  2.85091  2.56864  2.69301  2.52673  4.92396  3.67483     23
- -
          2.68618  4.42225  2.77519  2.73123  3.46354  2.40513
3.72494  3.29354  2.67741  2.69355  4.24690  2.90347  2.73739
3.18146  2.89801  2.37887  2.77519  2.98518  4.58477  3.61503
          0.02747  4.00411  4.72645  0.61958  0.77255  0.50504
0.92504
     12   2.35706  4.93789  2.52488  2.16923  4.22232  3.25683
3.44531  3.66777  2.29438  3.15628  3.90190  2.84102  3.76780
2.64718  2.81136  2.41318  2.71708  3.23333  5.40996  4.03728     24
- -
          2.68618  4.42225  2.77519  2.73123  3.46354  2.40513
3.72494  3.29354  2.67741  2.69355  4.24690  2.90347  2.73739
3.18146  2.89801  2.37887  2.77519  2.98518  4.58477  3.61503
          0.02747  4.00411  4.72645  0.61958  0.77255  0.50504
0.92504
     13   3.15844  4.98527  3.66296  3.14861  4.37339  3.61912
3.92654  3.92556  2.30000  3.41765  4.41579  3.49757  4.11656
3.14154  0.79488  3.23711  3.41624  3.64417  5.40386  4.27364     25
- -
          2.68618  4.42225  2.77519  2.73123  3.46354  2.40513
3.72494  3.29354  2.67741  2.69355  4.24690  2.90347  2.73739
3.18146  2.89801  2.37887  2.77519  2.98518  4.58477  3.61503
          0.02747  4.00411  4.72645  0.61958  0.77255  0.50504
0.92504
     14   2.56485  4.65109  3.08286  2.59119  4.10487  2.37502
3.71552  3.51856  2.42265  2.97122  3.96039  3.04319  3.81605
2.88828  2.11334  2.27977  2.85581  3.16589  5.34712  4.05531     26
- -
          2.68618  4.42225  2.77519  2.73123  3.46354  2.40513
3.72494  3.29354  2.67741  2.69355  4.24690  2.90347  2.73739
3.18146  2.89801  2.37887  2.77519  2.98518  4.58477  3.61503
          0.02747  4.00411  4.72645  0.61958  0.77255  0.50504
0.92504
     15   2.16437  4.32617  3.37252  2.78963  3.48211  3.51259
3.67700  2.41367  2.67194  2.54552  3.35341  3.22736  3.89887
3.06851  2.88465  2.72758  2.62137  2.46361  4.91086  3.66966     27
- -
          2.68618  4.42225  2.77519  2.73123  3.46354  2.40513
3.72494  3.29354  2.67741  2.69355  4.24690  2.90347  2.73739
3.18146  2.89801  2.37887  2.77519  2.98518  4.58477  3.61503
          0.02747  4.00411  4.72645  0.61958  0.77255  0.50504
0.92504
```

16 2.46174 4.83684 3.03429 2.49483 4.17438 3.41947
3.62497 3.57920 2.27160 3.12015 3.97341 2.98614 3.83541
2.35935 2.04668 2.47219 2.71786 3.24133 5.35285 4.05062 28
- -
2.68618 4.42225 2.77519 2.73123 3.46354 2.40513
3.72494 3.29354 2.67741 2.69355 4.24690 2.90347 2.73739
3.18146 2.89801 2.37887 2.77519 2.98518 4.58477 3.61503
0.02747 4.00411 4.72645 0.61958 0.77255 0.47390
0.97437
17 3.09302 4.61941 4.21342 3.82053 3.28161 3.94754
4.52101 2.47256 3.57968 1.85572 1.11700 4.07878 4.41101
3.95287 3.76690 3.45673 3.40235 2.49560 5.14109 3.91495 29
- -
2.68618 4.42225 2.77519 2.73123 3.46354 2.40513
3.72494 3.29354 2.67741 2.69355 4.24690 2.90347 2.73739
3.18146 2.89801 2.37887 2.77519 2.98518 4.58477 3.61503
0.02696 4.02262 4.74496 0.61958 0.77255 0.48576
0.95510
18 3.32709 4.74617 4.42998 4.07280 3.19879 4.17148
4.68160 2.41344 3.83907 0.71022 3.17896 4.33366 4.58250
4.17133 3.98965 3.76330 3.61534 2.48932 5.13017 3.88739 30
- -
2.68618 4.42225 2.77519 2.73123 3.46354 2.40513
3.72494 3.29354 2.67741 2.69355 4.24690 2.90347 2.73739
3.18146 2.89801 2.37887 2.77519 2.98518 4.58477 3.61503
0.02696 4.02262 4.74496 0.61958 0.77255 0.48576
0.95510
19 2.92428 4.96539 3.37342 2.79780 4.28977 3.58146
3.41584 3.76052 2.04394 3.27447 4.16283 3.19922 4.00306
2.83644 1.24115 2.96467 2.94684 3.45209 5.35706 4.11620 31
- -
2.68618 4.42225 2.77519 2.73123 3.46354 2.40513
3.72494 3.29354 2.67741 2.69355 4.24690 2.90347 2.73739
3.18146 2.89801 2.37887 2.77519 2.98518 4.58477 3.61503
0.02696 4.02262 4.74496 0.61958 0.77255 0.48576
0.95510
20 2.32754 4.23242 3.43794 3.17163 4.14106 3.04903
4.21350 3.38657 3.14650 3.21583 4.13437 3.35330 3.77615
3.50790 3.42624 2.33850 1.14665 2.97902 5.54496 4.30169 32
- -
2.68618 4.42225 2.77519 2.73123 3.46354 2.40513
3.72494 3.29354 2.67741 2.69355 4.24690 2.90347 2.73739
3.18146 2.89801 2.37887 2.77519 2.98518 4.58477 3.61503
0.02696 4.02262 4.74496 0.61958 0.77255 0.48576
0.95510
21 0.81728 4.27297 3.62041 3.42136 4.20706 3.09296
4.42492 3.35544 3.44162 3.22989 4.24718 3.53130 3.84663
3.76944 3.68019 2.62296 2.91086 2.98372 5.61916 4.43687 33
- -
2.68618 4.42225 2.77519 2.73123 3.46354 2.40513
3.72494 3.29354 2.67741 2.69355 4.24690 2.90347 2.73739
3.18146 2.89801 2.37887 2.77519 2.98518 4.58477 3.61503
0.02696 4.02262 4.74496 0.61958 0.77255 0.48576
0.95510

22 3.19985 4.55997 4.72221 4.14813 2.72494 4.41127
4.67527 2.19630 4.02916 1.14521 1.91779 4.40215 4.62096
4.13764 4.16163 3.73683 3.41869 2.23708 5.04138 3.93630 34
- -
2.68618 4.42225 2.77519 2.73123 3.46354 2.40513
3.72494 3.29354 2.67741 2.69355 4.24690 2.90347 2.73739
3.18146 2.89801 2.37887 2.77519 2.98518 4.58477 3.61503
0.02696 4.02262 4.74496 0.61958 0.77255 0.48576
0.95510
23 2.35923 3.41443 3.79079 3.52577 4.30782 0.89376
4.44179 3.70397 3.48788 3.46570 4.36057 3.56110 3.79794
3.79790 3.71475 2.56325 2.86499 3.20598 5.61468 4.48856 35
- -
2.68618 4.42225 2.77519 2.73123 3.46354 2.40513
3.72494 3.29354 2.67741 2.69355 4.24690 2.90347 2.73739
3.18146 2.89801 2.37887 2.77519 2.98518 4.58477 3.61503
0.02696 4.02262 4.74496 0.61958 0.77255 0.48576
0.95510
24 2.33471 4.46007 3.11827 2.76019 4.12964 3.12134
3.90036 3.52683 2.69882 3.13503 4.03984 3.13690 1.62368
3.11649 2.83449 2.51915 2.84142 3.13616 5.43408 4.16391 36
- -
2.68618 4.42225 2.77519 2.73123 3.46354 2.40513
3.72494 3.29354 2.67741 2.69355 4.24690 2.90347 2.73739
3.18146 2.89801 2.37887 2.77519 2.98518 4.58477 3.61503
0.02696 4.02262 4.74496 0.61958 0.77255 0.48576
0.95510
25 2.01566 4.40145 3.24369 2.62665 3.59127 3.47320
3.74965 2.93075 2.68222 2.32371 3.44358 3.16629 3.61127
3.01427 3.03741 2.66735 2.72847 2.55104 4.99820 3.74187 37
- -
2.68618 4.42225 2.77519 2.73123 3.46354 2.40513
3.72494 3.29354 2.67741 2.69355 4.24690 2.90347 2.73739
3.18146 2.89801 2.37887 2.77519 2.98518 4.58477 3.61503
0.02696 4.02262 4.74496 0.61958 0.77255 0.48576
0.95510
26 3.15498 4.47237 4.79650 4.26349 3.39751 4.48843
4.93221 1.10234 4.12894 1.79933 2.99301 4.51140 4.74729
4.35947 4.31280 3.85143 3.40216 1.66821 5.39618 4.23653 38
- -
2.68618 4.42225 2.77519 2.73123 3.46354 2.40513
3.72494 3.29354 2.67741 2.69355 4.24690 2.90347 2.73739
3.18146 2.89801 2.37887 2.77519 2.98518 4.58477 3.61503
0.02696 4.02262 4.74496 0.61958 0.77255 0.48576
0.95510
27 1.80235 4.45621 3.21952 2.67973 3.73620 3.13232
3.74490 3.03379 2.61984 2.70844 3.54302 3.12190 3.84049
2.90355 2.89174 2.57071 2.68569 2.82660 5.10582 3.83882 39
- -
2.68618 4.42225 2.77519 2.73123 3.46354 2.40513
3.72494 3.29354 2.67741 2.69355 4.24690 2.90347 2.73739
3.18146 2.89801 2.37887 2.77519 2.98518 4.58477 3.61503
0.02696 4.02262 4.74496 0.61958 0.77255 0.48576
0.95510

```
     28   2.05546  4.79072  2.92552  2.22467  4.07264  2.95250
3.51561  3.48640  2.40688  3.03480  3.90489  2.94326  3.80864
2.78921  2.58098  2.63841  2.73859  3.15746  5.32172  3.99439      40
- -
          2.68618  4.42225  2.77519  2.73123  3.46354  2.40513
3.72494  3.29354  2.67741  2.69355  4.24690  2.90347  2.73739
3.18146  2.89801  2.37887  2.77519  2.98518  4.58477  3.61503
          0.02696  4.02262  4.74496  0.61958  0.77255  0.48576
0.95510
     29   2.07985  4.20818  3.64798  3.07989  2.68925  3.60882
3.29566  2.68586  2.99999  2.26924  3.31518  3.41687  3.98277
3.28183  3.17035  2.87096  2.82577  2.47796  3.61001  3.20469      41
- -
          2.68618  4.42225  2.77519  2.73123  3.46354  2.40513
3.72494  3.29354  2.67741  2.69355  4.24690  2.90347  2.73739
3.18146  2.89801  2.37887  2.77519  2.98518  4.58477  3.61503
          0.02696  4.02262  4.74496  0.61958  0.77255  0.48576
0.95510
     30   3.27698  4.70300  4.53222  4.08186  3.09512  4.31512
4.67655  2.26529  3.81083  0.80343  2.95659  4.34634  4.63983
4.12118  3.97388  3.74950  3.54301  2.35550  5.10743  3.86784      42
- -
          2.68618  4.42225  2.77519  2.73123  3.46354  2.40513
3.72494  3.29354  2.67741  2.69355  4.24690  2.90347  2.73739
3.18146  2.89801  2.37887  2.77519  2.98518  4.58477  3.61503
          0.02696  4.02262  4.74496  0.61958  0.77255  0.48576
0.95510
     31   2.38721  5.16075  2.13940  1.71115  4.46703  3.31062
3.64473  3.93518  2.47673  3.45645  4.24789  2.78890  3.80050
2.75472  2.79872  2.64068  2.93235  3.53558  5.62829  4.21241      43
- -
          2.68618  4.42225  2.77519  2.73123  3.46354  2.40513
3.72494  3.29354  2.67741  2.69355  4.24690  2.90347  2.73739
3.18146  2.89801  2.37887  2.77519  2.98518  4.58477  3.61503
          0.02696  4.02262  4.74496  0.61958  0.77255  0.48576
0.95510
     32   2.73724  5.37359  1.10776  2.09189  4.72965  3.22528
3.82330  4.18233  2.86040  3.76815  4.65102  2.73543  3.86519
3.00387  3.44246  2.84595  3.24372  3.78165  5.95718  4.47416      44
- -
          2.68618  4.42225  2.77519  2.73123  3.46354  2.40513
3.72494  3.29354  2.67741  2.69355  4.24690  2.90347  2.73739
3.18146  2.89801  2.37887  2.77519  2.98518  4.58477  3.61503
          0.02696  4.02262  4.74496  0.61958  0.77255  0.48576
0.95510
     33   2.41449  4.68380  2.63936  2.48599  4.20897  3.23594
3.79740  3.62939  2.66231  3.25209  4.08832  2.89258  1.76610
2.94652  3.04132  2.57616  2.86285  3.25044  5.49048  4.16569      45
- -
          2.68618  4.42225  2.77519  2.73123  3.46354  2.40513
3.72494  3.29354  2.67741  2.69355  4.24690  2.90347  2.73739
3.18146  2.89801  2.37887  2.77519  2.98518  4.58477  3.61503
          0.02696  4.02262  4.74496  0.61958  0.77255  0.48576
0.95510
```

34  2.31098  4.92840  2.36516  2.27789  4.21761  3.10071
3.59749  3.65928  2.38193  3.12305  3.94624  2.86722  3.77865
2.65020  2.77081  2.34647  2.79901  3.29302  5.41849  4.04914  46
- -
2.68618  4.42225  2.77519  2.73123  3.46354  2.40513
3.72494  3.29354  2.67741  2.69355  4.24690  2.90347  2.73739
3.18146  2.89801  2.37887  2.77519  2.98518  4.58477  3.61503
0.02696  4.02262  4.74496  0.61958  0.77255  0.48576
0.95510
35  2.96947  4.35886  4.55241  4.05094  3.60581  4.20806
4.80235  1.61031  3.93576  2.19753  3.43296  4.28088  4.58679
4.22750  4.16177  3.58108  2.98882  1.06698  5.46010  4.24928  47
- -
2.68618  4.42225  2.77519  2.73123  3.46354  2.40513
3.72494  3.29354  2.67741  2.69355  4.24690  2.90347  2.73739
3.18146  2.89801  2.37887  2.77519  2.98518  4.58477  3.61503
0.02696  4.02262  4.74496  0.61958  0.77255  0.48576
0.95510
36  2.63582  4.27885  4.27330  3.76898  3.57396  3.71743
4.52054  1.75265  3.66133  2.31919  3.45423  4.00192  4.36702
3.95538  3.90072  3.29287  3.09497  1.17027  5.28335  4.06646  48
- -
2.68618  4.42225  2.77519  2.73123  3.46354  2.40513
3.72494  3.29354  2.67741  2.69355  4.24690  2.90347  2.73739
3.18146  2.89801  2.37887  2.77519  2.98518  4.58477  3.61503
0.02696  4.02262  4.74496  0.61958  0.77255  0.48576
0.95510
37  3.08846  5.22327  2.56059  0.82931  4.58940  3.37593
3.97883  4.07348  2.87880  3.66752  4.63740  3.02771  3.98839
3.19292  3.30140  3.05703  3.39464  3.74275  5.73268  4.45792  49
- -
2.68618  4.42225  2.77519  2.73123  3.46354  2.40513
3.72494  3.29354  2.67741  2.69355  4.24690  2.90347  2.73739
3.18146  2.89801  2.37887  2.77519  2.98518  4.58477  3.61503
0.02696  4.02262  4.74496  0.61958  0.77255  0.48576
0.95510
38  3.07605  4.41612  4.73312  4.24631  3.58108  4.37861
4.97828  1.52935  4.12658  2.10237  3.39984  4.46738  4.72077
4.40835  4.33479  3.77100  3.35083  1.01637  5.53084  4.31194  50
- -
2.68618  4.42225  2.77519  2.73123  3.46354  2.40513
3.72494  3.29354  2.67741  2.69355  4.24690  2.90347  2.73739
3.18146  2.89801  2.37887  2.77519  2.98518  4.58477  3.61503
0.02696  4.02262  4.74496  0.61958  0.77255  0.48576
0.95510
39  3.09302  4.61941  4.21342  3.82053  3.28161  3.94754
4.52101  2.47256  3.57968  1.85572  1.11700  4.07878  4.41101
3.95287  3.76690  3.45673  3.40235  2.49560  5.14109  3.91495  51
- -
2.68618  4.42225  2.77519  2.73123  3.46354  2.40513
3.72494  3.29354  2.67741  2.69355  4.24690  2.90347  2.73739
3.18146  2.89801  2.37887  2.77519  2.98518  4.58477  3.61503
0.02696  4.02262  4.74496  0.61958  0.77255  0.48576
0.95510

```
     40   2.82966  4.49111  4.37636  3.88327  3.27203  4.10209
4.59636  2.14998  3.71128  1.05610  3.12456  4.14348  4.48059
3.99594  3.91771  3.47546  3.29518  1.94826  5.19921  4.02699     52
- -
          2.68618  4.42225  2.77519  2.73123  3.46354  2.40513
3.72494  3.29354  2.67741  2.69355  4.24690  2.90347  2.73739
3.18146  2.89801  2.37887  2.77519  2.98518  4.58477  3.61503
          0.02696  4.02262  4.74496  0.61958  0.77255  0.48576
0.95510
     41   2.85426  4.85176  2.80169  2.70353  4.23951  3.28976
4.05172  3.98473  2.96940  3.61516  4.57129  0.91734  3.95002
3.32667  3.32750  2.91694  3.24227  3.59163  5.52835  4.17856     53
- -
          2.68618  4.42225  2.77519  2.73123  3.46354  2.40513
3.72494  3.29354  2.67741  2.69355  4.24690  2.90347  2.73739
3.18146  2.89801  2.37887  2.77519  2.98518  4.58477  3.61503
          0.02696  4.02262  4.74496  0.61958  0.77255  0.48576
0.95510
     42   2.48890  4.47103  3.32141  3.14997  4.26228  3.18238
4.28322  3.68779  3.21714  3.37082  4.36974  3.42592  0.94042
3.59333  3.49540  2.75487  3.05099  3.29393  5.57869  4.38375     54
- -
          2.68618  4.42225  2.77519  2.73123  3.46354  2.40513
3.72494  3.29354  2.67741  2.69355  4.24690  2.90347  2.73739
3.18146  2.89801  2.37887  2.77519  2.98518  4.58477  3.61503
          0.02696  4.02262  4.74496  0.61958  0.77255  0.48576
0.95510
     43   3.15706  5.30096  0.72443  2.44038  4.69960  3.32995
4.07187  4.32326  3.18574  3.90577  4.88941  3.00343  3.99358
3.31287  3.72060  3.10404  3.49962  3.94721  5.83844  4.55063     55
- -
          2.68618  4.42225  2.77519  2.73123  3.46354  2.40513
3.72494  3.29354  2.67741  2.69355  4.24690  2.90347  2.73739
3.18146  2.89801  2.37887  2.77519  2.98518  4.58477  3.61503
          0.02696  4.02262  4.74496  0.61958  0.77255  0.48576
0.95510
     44   2.65527  4.63846  3.16157  2.82806  4.30888  1.34001
3.94341  3.78300  2.63266  3.38504  4.25591  3.21742  3.88439
3.15663  2.17360  2.75924  3.01835  3.38059  5.49881  4.27630     56
- -
          2.68618  4.42225  2.77519  2.73123  3.46354  2.40513
3.72494  3.29354  2.67741  2.69355  4.24690  2.90347  2.73739
3.18146  2.89801  2.37887  2.77519  2.98518  4.58477  3.61503
          0.02696  4.02262  4.74496  0.61958  0.77255  0.48576
0.95510
     45   2.47470  4.53148  3.11870  2.55423  4.00522  3.41774
3.65840  3.39503  2.29522  3.01420  3.84899  3.03152  3.83526
2.79912  2.01273  2.53393  2.47474  3.00119  5.26075  3.97074     57
- -
          2.68618  4.42225  2.77519  2.73123  3.46354  2.40513
3.72494  3.29354  2.67741  2.69355  4.24690  2.90347  2.73739
3.18146  2.89801  2.37887  2.77519  2.98518  4.58477  3.61503
          0.02696  4.02262  4.74496  0.61958  0.77255  0.48576
0.95510
```

```
     46   3.26136  4.60353  4.78892  4.24967  3.17004  4.51192
4.86370  1.81572  4.08903  0.96343  2.97918  4.51855  4.73540
4.27160  4.24576  3.87321  3.49843  1.99622  5.23714  4.11677      58
- -
          2.68618  4.42225  2.77519  2.73123  3.46354  2.40513
3.72494  3.29354  2.67741  2.69355  4.24690  2.90347  2.73739
3.18146  2.89801  2.37887  2.77519  2.98518  4.58477  3.61503
          0.02696  4.02262  4.74496  0.61958  0.77255  0.48576
0.95510
     47   3.03248  4.65789  3.84572  3.34493  2.93094  3.75268
3.80923  3.24819  2.83659  2.77431  3.83359  3.61761  4.19286
3.41374  2.53452  3.21214  3.28451  3.06564  1.29506  2.94089      59
- -
          2.68618  4.42225  2.77519  2.73123  3.46354  2.40513
3.72494  3.29354  2.67741  2.69355  4.24690  2.90347  2.73739
3.18146  2.89801  2.37887  2.77519  2.98518  4.58477  3.61503
          0.02696  4.02262  4.74496  0.61958  0.77255  0.48576
0.95510
     48   3.09388  4.40673  4.77894  4.25035  3.49907  4.44873
4.93004  1.45136  4.14092  1.81832  3.30518  4.48336  4.72835
4.38009  4.33353  3.80974  3.28063  1.21222  5.44029  4.26120      60
- -
          2.68618  4.42225  2.77519  2.73123  3.46354  2.40513
3.72494  3.29354  2.67741  2.69355  4.24690  2.90347  2.73739
3.18146  2.89801  2.37887  2.77519  2.98518  4.58477  3.61503
          0.02696  4.02262  4.74496  0.61958  0.77255  0.48576
0.95510
     49   3.02893  5.48400  0.98636  2.05910  4.79876  3.24293
3.86424  4.32108  2.93478  3.87410  4.77862  2.75157  3.89584
3.05466  3.52904  2.92156  3.34072  3.91678  5.99549  4.52634      61
- -
          2.68618  4.42225  2.77519  2.73123  3.46354  2.40513
3.72494  3.29354  2.67741  2.69355  4.24690  2.90347  2.73739
3.18146  2.89801  2.37887  2.77519  2.98518  4.58477  3.61503
          0.02696  4.02262  4.74496  0.61958  0.77255  0.48576
0.95510
     50   2.91649  4.71513  3.49459  2.83325  4.26425  3.61800
3.66509  3.62926  1.96064  3.02631  4.08375  3.22724  4.01096
2.82722  1.27299  2.96958  3.12216  3.34530  5.33743  4.12317      62
- -
          2.68618  4.42225  2.77519  2.73123  3.46354  2.40513
3.72494  3.29354  2.67741  2.69355  4.24690  2.90347  2.73739
3.18146  2.89801  2.37887  2.77519  2.98518  4.58477  3.61503
          0.02696  4.02262  4.74496  0.61958  0.77255  0.48576
0.95510
     51   2.89892  4.56515  4.39049  3.90279  2.92728  4.18185
4.52506  2.25266  3.75023  0.94788  3.04327  4.17187  4.51816
3.99526  3.94638  3.54976  3.36777  2.28750  5.02583  3.76945      63
- -
          2.68618  4.42225  2.77519  2.73123  3.46354  2.40513
3.72494  3.29354  2.67741  2.69355  4.24690  2.90347  2.73739
3.18146  2.89801  2.37887  2.77519  2.98518  4.58477  3.61503
          0.02696  4.02262  4.74496  0.61958  0.77255  0.48576
0.95510
```

```
     52   2.36558  4.69870  2.75321  2.47022  4.17061  2.66434
3.70982  3.59746  2.52445  3.19535  4.00597  2.95790  3.77813
2.86879  2.65422  1.90356  2.82855  3.18707  5.42152  4.09106     64
- -
          2.68618  4.42225  2.77519  2.73123  3.46354  2.40513
3.72494  3.29354  2.67741  2.69355  4.24690  2.90347  2.73739
3.18146  2.89801  2.37887  2.77519  2.98518  4.58477  3.61503
          0.02696  4.02262  4.74496  0.61958  0.77255  0.48576
0.95510
     53   2.56802  4.87373  2.75394  1.98315  4.14744  3.12210
3.61193  3.57642  2.40698  2.97671  3.95925  2.88914  3.78867
2.65771  2.87865  2.26832  2.79682  3.04124  5.37671  4.02130     65
- -
          2.68618  4.42225  2.77519  2.73123  3.46354  2.40513
3.72494  3.29354  2.67741  2.69355  4.24690  2.90347  2.73739
3.18146  2.89801  2.37887  2.77519  2.98518  4.58477  3.61503
          0.02696  4.02262  4.74496  0.61958  0.77255  0.48576
0.95510
     54   2.86181  4.62005  3.56615  3.48578  4.62096  0.53093
4.59479  4.25913  3.68734  3.89379  4.87234  3.73076  4.01589
3.99980  3.90487  3.03839  3.35475  3.75848  5.67431  4.72868     66
- -
          2.68618  4.42225  2.77519  2.73123  3.46354  2.40513
3.72494  3.29354  2.67741  2.69355  4.24690  2.90347  2.73739
3.18146  2.89801  2.37887  2.77519  2.98518  4.58477  3.61503
          0.02696  4.02262  4.74496  0.61958  0.77255  0.48576
0.95510
     55   2.76103  4.44224  3.67578  3.08390  3.39328  3.73718
3.94570  2.46349  2.81521  1.67935  3.10812  3.47406  4.09229
3.25292  2.26228  3.00913  2.98718  2.51542  4.98560  3.77088     67
- -
          2.68618  4.42225  2.77519  2.73123  3.46354  2.40513
3.72494  3.29354  2.67741  2.69355  4.24690  2.90347  2.73739
3.18146  2.89801  2.37887  2.77519  2.98518  4.58477  3.61503
          0.02696  4.02262  4.74496  0.61958  0.77255  0.48576
0.95510
     56   2.04298  4.49917  3.16374  2.39385  3.62329  3.33692
3.70350  2.90730  2.59106  2.75170  3.61437  3.08702  3.84800
2.90177  2.94361  2.58262  2.75558  2.57565  5.07371  3.79745     68
- -
          2.68618  4.42225  2.77519  2.73123  3.46354  2.40513
3.72494  3.29354  2.67741  2.69355  4.24690  2.90347  2.73739
3.18146  2.89801  2.37887  2.77519  2.98518  4.58477  3.61503
          0.02696  4.02262  4.74496  0.61958  0.77255  0.48576
0.95510
     57   2.54500  4.28455  2.05053  2.34578  4.14339  3.34519
3.63852  3.56036  2.44483  3.15722  3.96441  2.90752  2.91204
2.78414  2.88545  2.58670  2.83549  3.07106  5.38452  4.03429     69
- -
          2.68618  4.42225  2.77519  2.73123  3.46354  2.40513
3.72494  3.29354  2.67741  2.69355  4.24690  2.90347  2.73739
3.18146  2.89801  2.37887  2.77519  2.98518  4.58477  3.61503
          0.02696  4.02262  4.74496  0.61958  0.77255  0.48576
0.95510
```

```
     58   2.46977  4.30570  3.49208  2.95782  3.56469  3.45876
3.90135  2.75552  2.86923  2.39653  3.03886  3.32661  3.92376
3.20991  3.20727  2.60305  1.82647  2.52907  5.03029  3.79435     70
- -
          2.68618  4.42225  2.77519  2.73123  3.46354  2.40513
3.72494  3.29354  2.67741  2.69355  4.24690  2.90347  2.73739
3.18146  2.89801  2.37887  2.77519  2.98518  4.58477  3.61503
          0.02696  4.02262  4.74496  0.61958  0.77255  0.48576
0.95510
     59   2.55814  4.78661  2.20252  2.40955  4.48845  1.40568
3.90813  3.95418  2.89466  3.56970  4.41601  2.85537  3.80615
3.10747  3.38282  2.67149  3.01172  3.49965  5.75934  4.38810     71
- -
          2.68618  4.42225  2.77519  2.73123  3.46354  2.40513
3.72494  3.29354  2.67741  2.69355  4.24690  2.90347  2.73739
3.18146  2.89801  2.37887  2.77519  2.98518  4.58477  3.61503
          0.02696  4.02262  4.74496  0.61958  0.77255  0.48576
0.95510
     60   2.49597  4.42122  2.86903  2.17954  3.86172  3.42756
3.64103  3.15750  2.46078  2.74588  3.68718  2.99304  3.81670
2.70361  2.84914  2.64060  2.76813  2.50969  5.17864  3.87089     72
- -
          2.68618  4.42225  2.77519  2.73123  3.46354  2.40513
3.72494  3.29354  2.67741  2.69355  4.24690  2.90347  2.73739
3.18146  2.89801  2.37887  2.77519  2.98518  4.58477  3.61503
          0.02696  4.02262  4.74496  0.61958  0.77255  0.48576
0.95510
     61   2.58465  4.71451  3.02625  2.23041  3.87480  3.42387
3.53774  3.31548  2.40539  2.94119  3.72586  2.97462  3.81348
2.79103  2.43175  2.56168  2.42783  2.81916  5.22585  3.86549     73
- -
          2.68618  4.42225  2.77519  2.73123  3.46354  2.40513
3.72494  3.29354  2.67741  2.69355  4.24690  2.90347  2.73739
3.18146  2.89801  2.37887  2.77519  2.98518  4.58477  3.61503
          0.02696  4.02262  4.74496  0.61958  0.77255  0.48576
0.95510
     62   3.26318  4.59092  4.83811  4.27045  3.02364  4.53245
4.83761  1.84331  4.15294  0.98619  2.62230  4.53422  4.71827
4.26015  4.28454  3.87146  3.48436  2.10831  5.17154  4.10067     74
- -
          2.68618  4.42225  2.77519  2.73123  3.46354  2.40513
3.72494  3.29354  2.67741  2.69355  4.24690  2.90347  2.73739
3.18146  2.89801  2.37887  2.77519  2.98518  4.58477  3.61503
          0.02696  4.02262  4.74496  0.61958  0.77255  0.48576
0.95510
     63   2.41360  4.71558  2.90569  2.39179  4.10021  3.15658
3.56688  3.52150  2.50346  3.12605  3.94187  2.77482  3.06732
2.84311  2.95778  2.03321  2.53955  3.16815  5.36540  4.03563     75
- -
          2.68618  4.42225  2.77519  2.73123  3.46354  2.40513
3.72494  3.29354  2.67741  2.69355  4.24690  2.90347  2.73739
3.18146  2.89801  2.37887  2.77519  2.98518  4.58477  3.61503
          0.02696  4.02262  4.74496  0.61958  0.77255  0.48576
0.95510
```

64 2.13160 4.79902 2.81838 2.12229 4.16555 3.33317
3.68907 3.57452 2.48250 3.17886 4.00510 2.93642 2.44202
2.85047 2.79646 2.65287 2.88790 3.22745 5.41251 4.08062 76
- -
2.68618 4.42225 2.77519 2.73123 3.46354 2.40513
3.72494 3.29354 2.67741 2.69355 4.24690 2.90347 2.73739
3.18146 2.89801 2.37887 2.77519 2.98518 4.58477 3.61503
0.02696 4.02262 4.74496 0.61958 0.77255 0.48576
0.95510
65 1.62852 4.58982 2.97830 2.52065 3.97509 3.27586
3.48532 3.37593 2.58855 3.02051 3.86970 3.03960 3.82471
2.92387 2.92608 2.41843 2.82007 3.05394 5.29164 4.00107 77
- -
2.68618 4.42225 2.77519 2.73123 3.46354 2.40513
3.72494 3.29354 2.67741 2.69355 4.24690 2.90347 2.73739
3.18146 2.89801 2.37887 2.77519 2.98518 4.58477 3.61503
0.02696 4.02262 4.74496 0.61958 0.77255 0.48576
0.95510
66 2.98340 5.56072 1.14813 1.91447 4.83212 3.22706
3.79017 4.36120 2.82860 3.86249 4.72613 2.64543 3.86264
2.87247 3.43102 2.86008 3.27741 3.93585 6.02356 4.50327 78
- -
2.68618 4.42225 2.77519 2.73123 3.46354 2.40513
3.72494 3.29354 2.67741 2.69355 4.24690 2.90347 2.73739
3.18146 2.89801 2.37887 2.77519 2.98518 4.58477 3.61503
0.02696 4.02262 4.74496 0.61958 0.77255 0.48576
0.95510
67 1.89806 4.30778 3.32279 2.90556 4.03263 1.77211
3.97959 3.37967 2.87514 3.09033 3.94777 3.21786 3.76752
3.22262 3.00124 2.52488 2.76373 2.68900 5.38856 4.14179 79
- -
2.68618 4.42225 2.77519 2.73123 3.46354 2.40513
3.72494 3.29354 2.67741 2.69355 4.24690 2.90347 2.73739
3.18146 2.89801 2.37887 2.77519 2.98518 4.58477 3.61503
0.02696 4.02262 4.74496 0.61958 0.77255 0.48576
0.95510
68 3.01368 5.26401 2.39764 0.93623 4.61151 3.32603
3.89459 4.08734 2.81135 3.68249 4.60583 2.91001 3.93780
3.09101 3.27166 2.96207 3.31408 3.73183 5.77856 4.43062 80
- -
2.68618 4.42225 2.77519 2.73123 3.46354 2.40513
3.72494 3.29354 2.67741 2.69355 4.24690 2.90347 2.73739
3.18146 2.89801 2.37887 2.77519 2.98518 4.58477 3.61503
0.02696 4.02262 4.74496 0.61958 0.77255 0.48576
0.95510
69 3.18120 5.00542 3.68780 3.17038 4.39834 3.63746
3.94268 3.95237 2.31116 3.44111 4.43934 3.51762 4.13451
3.15727 0.77162 3.25933 3.43807 3.67032 5.42160 4.29519 81
- -
2.68618 4.42225 2.77519 2.73123 3.46354 2.40513
3.72494 3.29354 2.67741 2.69355 4.24690 2.90347 2.73739
3.18146 2.89801 2.37887 2.77519 2.98518 4.58477 3.61503
0.02696 4.02262 4.74496 0.61958 0.77255 0.48576
0.95510

70 3.11119 4.46777 4.70557 4.24500 3.48636 4.35843
4.95445 1.01305 4.09156 1.98149 3.32576 4.46366 4.71286
4.38255 4.28469 3.76975 3.39259 1.63714 5.46980 4.24688 82
- -
2.68618 4.42225 2.77519 2.73123 3.46354 2.40513
3.72494 3.29354 2.67741 2.69355 4.24690 2.90347 2.73739
3.18146 2.89801 2.37887 2.77519 2.98518 4.58477 3.61503
0.02696 4.02262 4.74496 0.61958 0.77255 0.48576
0.95510
71 3.13656 4.43568 4.84937 4.33147 3.50523 4.51234
5.01671 1.21193 4.21771 1.87999 3.30545 4.56155 4.78359
4.45774 4.40603 3.88627 3.39000 1.34692 5.49058 4.30712 83
- -
2.68618 4.42225 2.77519 2.73123 3.46354 2.40513
3.72494 3.29354 2.67741 2.69355 4.24690 2.90347 2.73739
3.18146 2.89801 2.37887 2.77519 2.98518 4.58477 3.61503
0.02696 4.02262 4.74496 0.61958 0.77255 0.48576
0.95510
72 3.07513 5.07244 3.51319 2.91331 4.49254 3.65339
3.70331 3.89347 1.91091 3.37854 4.29270 3.28796 4.07154
2.86318 1.06253 3.10641 3.27420 3.59448 5.42882 4.26099 84
- -
2.68618 4.42225 2.77519 2.73123 3.46354 2.40513
3.72494 3.29354 2.67741 2.69355 4.24690 2.90347 2.73739
3.18146 2.89801 2.37887 2.77519 2.98518 4.58477 3.61503
0.02696 4.02262 4.74496 0.61958 0.77255 0.48576
0.95510
73 3.32709 4.74617 4.42998 4.07280 3.19879 4.17148
4.68160 2.41344 3.83907 0.71022 3.17896 4.33366 4.58250
4.17133 3.98965 3.76330 3.61534 2.48932 5.13017 3.88739 85
- -
2.68618 4.42225 2.77519 2.73123 3.46354 2.40513
3.72494 3.29354 2.67741 2.69355 4.24690 2.90347 2.73739
3.18146 2.89801 2.37887 2.77519 2.98518 4.58477 3.61503
0.02696 4.02262 4.74496 0.61958 0.77255 0.48576
0.95510
74 2.92784 4.43147 4.33116 4.00117 3.66379 3.87690
4.77821 2.05427 3.87125 2.32619 3.60830 4.16627 4.43068
4.22125 4.06895 3.39165 3.28677 0.85478 5.47286 4.21283 86
- -
2.68618 4.42225 2.77519 2.73123 3.46354 2.40513
3.72494 3.29354 2.67741 2.69355 4.24690 2.90347 2.73739
3.18146 2.89801 2.37887 2.77519 2.98518 4.58477 3.61503
0.02696 4.02262 4.74496 0.61958 0.77255 0.48576
0.95510
75 0.81728 4.27297 3.62041 3.42136 4.20706 3.09296
4.42492 3.35544 3.44162 3.22989 4.24718 3.53130 3.84663
3.76944 3.68019 2.62296 2.91086 2.98372 5.61916 4.43687 87
- -
2.68618 4.42225 2.77519 2.73123 3.46354 2.40513
3.72494 3.29354 2.67741 2.69355 4.24690 2.90347 2.73739
3.18146 2.89801 2.37887 2.77519 2.98518 4.58477 3.61503
0.02696 4.02262 4.74496 0.61958 0.77255 0.48576
0.95510

```
     76   2.11891  4.58616  2.99596  2.61073  3.98428  3.28063
2.10710  3.47818  2.60966  3.11274  3.96273  3.05526  3.81792
2.99402  2.99745  2.23522  2.86485  3.12786  5.31137  3.98071      88
- -
          2.68618  4.42225  2.77519  2.73123  3.46354  2.40513
3.72494  3.29354  2.67741  2.69355  4.24690  2.90347  2.73739
3.18146  2.89801  2.37887  2.77519  2.98518  4.58477  3.61503
          0.02696  4.02262  4.74496  0.61958  0.77255  0.48576
0.95510
     77   2.55845  4.16799  3.16223  2.73689  3.77741  3.10106
1.67599  3.41067  2.61815  3.05227  3.92223  3.14266  3.84995
3.05163  2.96249  2.55104  2.88084  3.07711  5.15823  3.78866      89
- -
          2.68618  4.42225  2.77519  2.73123  3.46354  2.40513
3.72494  3.29354  2.67741  2.69355  4.24690  2.90347  2.73739
3.18146  2.89801  2.37887  2.77519  2.98518  4.58477  3.61503
          0.02696  4.02262  4.74496  0.61958  0.77255  0.48576
0.95510
     78   2.67212  4.31809  4.31590  3.84339  3.65941  3.92159
4.64286  1.86860  3.72500  2.32056  3.50773  4.05737  4.39852
4.04052  3.96752  3.30956  2.94481  1.05963  5.43160  4.21520      90
- -
          2.68618  4.42225  2.77519  2.73123  3.46354  2.40513
3.72494  3.29354  2.67741  2.69355  4.24690  2.90347  2.73739
3.18146  2.89801  2.37887  2.77519  2.98518  4.58477  3.61503
          0.02696  4.02262  4.74496  0.61958  0.77255  0.48576
0.95510
     79   2.63366  4.72581  2.86167  2.56163  4.25999  1.60066
3.79245  3.71032  2.51687  3.30394  4.14911  2.83983  3.84079
2.88352  2.54690  2.69673  2.95621  3.32806  5.48853  4.18439      91
- -
          2.68618  4.42225  2.77519  2.73123  3.46354  2.40513
3.72494  3.29354  2.67741  2.69355  4.24690  2.90347  2.73739
3.18146  2.89801  2.37887  2.77519  2.98518  4.58477  3.61503
          0.02696  4.02262  4.74496  0.61958  0.77255  0.48576
0.95510
     80   1.51326  4.31773  3.36109  2.77072  3.82680  3.22387
3.96609  2.96674  2.90247  2.84015  3.74888  3.26917  3.85052
3.23734  3.25495  2.64439  2.66183  2.31465  5.24719  4.00473      92
- -
          2.68618  4.42225  2.77519  2.73123  3.46354  2.40513
3.72494  3.29354  2.67741  2.69355  4.24690  2.90347  2.73739
3.18146  2.89801  2.37887  2.77519  2.98518  4.58477  3.61503
          0.02696  4.02262  4.74496  0.61958  0.77255  0.48576
0.95510
     81   2.94064  5.19180  2.44185  1.00964  4.62051  2.98738
3.87307  4.09629  2.77536  3.67472  4.56368  2.89869  3.91056
3.06210  3.23411  2.90217  3.24790  3.71382  5.77696  4.43117      93
- -
          2.68618  4.42225  2.77519  2.73123  3.46354  2.40513
3.72494  3.29354  2.67741  2.69355  4.24690  2.90347  2.73739
3.18146  2.89801  2.37887  2.77519  2.98518  4.58477  3.61503
          0.02696  4.02262  4.74496  0.61958  0.77255  0.48576
0.95510
```

82 2.80679 4.37419 4.40040 3.98188 3.63585 3.97781
4.76981 1.88066 3.84041 2.24623 3.50598 4.16802 4.47374
4.17563 4.06235 3.40046 3.22342 0.96348 5.49091 4.24082 94
- -
2.68618 4.42225 2.77519 2.73123 3.46354 2.40513
3.72494 3.29354 2.67741 2.69355 4.24690 2.90347 2.73739
3.18146 2.89801 2.37887 2.77519 2.98518 4.58477 3.61503
0.02696 4.02262 4.74496 0.61958 0.77255 0.48576
0.95510
83 2.82504 5.01149 2.47955 2.40249 3.92878 3.35898
1.56515 3.79138 2.58520 3.35324 4.23089 2.76247 3.89684
2.95790 2.99303 2.82050 3.09922 3.45309 5.29350 3.81801 95
- -
2.68618 4.42225 2.77519 2.73123 3.46354 2.40513
3.72494 3.29354 2.67741 2.69355 4.24690 2.90347 2.73739
3.18146 2.89801 2.37887 2.77519 2.98518 4.58477 3.61503
0.02696 4.02262 4.74496 0.61958 0.77255 0.48576
0.95510
84 2.16886 4.89197 2.85922 2.21935 4.17411 3.30634
3.50980 3.60427 2.35640 3.17243 3.97166 2.89945 3.47503
2.65442 2.42297 2.50310 2.81941 3.25282 5.37978 4.02971 96
- -
2.68618 4.42225 2.77519 2.73123 3.46354 2.40513
3.72494 3.29354 2.67741 2.69355 4.24690 2.90347 2.73739
3.18146 2.89801 2.37887 2.77519 2.98518 4.58477 3.61503
0.02696 4.02262 4.74496 0.61958 0.77255 0.48576
0.95510
85 2.27940 4.75254 2.81616 2.38407 4.09946 2.22472
3.61569 3.51680 2.48646 3.09121 3.94484 2.94562 3.80342
2.84191 2.69389 2.46219 2.85535 3.17536 5.36246 4.03476 97
- -
2.68618 4.42225 2.77519 2.73123 3.46354 2.40513
3.72494 3.29354 2.67741 2.69355 4.24690 2.90347 2.73739
3.18146 2.89801 2.37887 2.77519 2.98518 4.58477 3.61503
0.02696 4.02262 4.74496 0.61958 0.77255 0.48576
0.95510
86 2.21857 4.76905 2.95438 2.44684 4.10239 3.36501
3.64329 3.52007 2.40904 3.11426 3.92779 2.78458 3.79880
2.44737 2.65529 2.21637 2.69959 3.14188 5.34236 4.01536 98
- -
2.68618 4.42225 2.77519 2.73123 3.46354 2.40513
3.72494 3.29354 2.67741 2.69355 4.24690 2.90347 2.73739
3.18146 2.89801 2.37887 2.77519 2.98518 4.58477 3.61503
0.02696 4.02262 4.74496 0.61958 0.77255 0.48576
0.95510
87 2.52616 4.43809 3.23851 3.06762 4.22632 3.13794
4.21697 3.72990 3.14243 3.41577 4.36624 3.34888 1.02423
3.51864 3.43129 2.57508 2.98769 3.29832 5.55894 4.30616 99
- -
2.68618 4.42225 2.77519 2.73123 3.46354 2.40513
3.72494 3.29354 2.67741 2.69355 4.24690 2.90347 2.73739
3.18146 2.89801 2.37887 2.77519 2.98518 4.58477 3.61503
0.02696 4.02262 4.74496 0.61958 0.77255 0.48576
0.95510

```
     88   2.75470  4.49702  3.58748  2.99219  3.52024  3.68998
3.88402  2.34992  2.65776  2.03049  3.44264  3.39519  4.05898
3.15731  1.97673  2.96708  2.98267  2.51467  5.04511  3.80701    100
- -
          2.68618  4.42225  2.77519  2.73123  3.46354  2.40513
3.72494  3.29354  2.67741  2.69355  4.24690  2.90347  2.73739
3.18146  2.89801  2.37887  2.77519  2.98518  4.58477  3.61503
          0.02696  4.02262  4.74496  0.61958  0.77255  0.48576
0.95510
     89   3.10771  4.45874  4.68120  4.15161  3.36896  4.38313
4.82125  1.83312  4.01567  1.46510  3.19678  4.40195  4.67218
4.25614  4.20876  3.74180  3.35950  1.25833  5.32748  4.15040    101
- -
          2.68618  4.42225  2.77519  2.73123  3.46354  2.40513
3.72494  3.29354  2.67741  2.69355  4.24690  2.90347  2.73739
3.18146  2.89801  2.37887  2.77519  2.98518  4.58477  3.61503
          0.02696  4.02262  4.74496  0.61958  0.77255  0.48576
0.95510
     90   2.21700  4.20982  3.38425  3.11040  4.28641  2.97978
4.20097  3.63603  3.14609  3.37605  4.23061  3.29506  3.72025
3.46786  3.45501  1.14168  2.32869  3.13282  5.64011  4.40878    102
- -
          2.68618  4.42225  2.77519  2.73123  3.46354  2.40513
3.72494  3.29354  2.67741  2.69355  4.24690  2.90347  2.73739
3.18146  2.89801  2.37887  2.77519  2.98518  4.58477  3.61503
          0.02696  4.02262  4.74496  0.61958  0.77255  0.48576
0.95510
     91   0.88356  4.21198  3.52839  3.32006  4.28353  2.83291
4.35865  3.52879  3.36974  3.36755  4.28205  3.43317  3.77304
3.67878  3.62827  2.51503  2.81796  3.07330  5.65280  4.46827    103
- -
          2.68618  4.42225  2.77519  2.73123  3.46354  2.40513
3.72494  3.29354  2.67741  2.69355  4.24690  2.90347  2.73739
3.18146  2.89801  2.37887  2.77519  2.98518  4.58477  3.61503
          0.02696  4.02262  4.74496  0.61958  0.77255  0.48576
0.95510
     92   2.89870  5.28625  2.28738  1.15446  4.59342  3.29409
3.76406  4.06859  2.64285  3.61654  4.47537  2.79924  3.87247
2.93158  3.00362  2.83262  3.17576  3.68859  5.77155  4.35608    104
- -
          2.68618  4.42225  2.77519  2.73123  3.46354  2.40513
3.72494  3.29354  2.67741  2.69355  4.24690  2.90347  2.73739
3.18146  2.89801  2.37887  2.77519  2.98518  4.58477  3.61503
          0.02696  4.02262  4.74496  0.61958  0.77255  0.48576
0.95510
     93   3.32779  4.70936  4.63990  4.16729  2.76600  4.41443
4.61773  2.27565  3.99584  0.80986  2.97704  4.41297  4.68477
4.18479  4.14582  3.82145  3.57394  2.38831  4.94145  3.61905    105
- -
          2.68618  4.42225  2.77519  2.73123  3.46354  2.40513
3.72494  3.29354  2.67741  2.69355  4.24690  2.90347  2.73739
3.18146  2.89801  2.37887  2.77519  2.98518  4.58477  3.61503
          0.02696  4.02262  4.74496  0.61958  0.77255  0.48576
0.95510
```

94 2.96212 4.69205 3.58591 3.44827 4.42684 3.37117
4.51157 4.01315 3.51887 3.62279 4.68053 3.73880 0.59042
3.90306 3.74552 3.13480 3.41594 3.64102 5.57002 4.55027 106
- -
2.68618 4.42225 2.77519 2.73123 3.46354 2.40513
3.72494 3.29354 2.67741 2.69355 4.24690 2.90347 2.73739
3.18146 2.89801 2.37887 2.77519 2.98518 4.58477 3.61503
0.02696 4.02262 4.74496 0.61958 0.77255 0.48576
0.95510
95 2.65219 4.95592 2.62692 1.68444 4.30124 2.74063
3.66268 3.74091 2.48686 3.21150 4.11369 2.85430 3.43828
2.80748 2.96773 2.60141 2.70018 3.36711 5.51318 4.13874 107
- -
2.68613 4.42227 2.77522 2.73121 3.46356 2.40507
3.72497 3.29356 2.67743 2.69354 4.24692 2.90349 2.73742
3.18142 2.89794 2.37889 2.77522 2.98521 4.58479 3.61505
0.11455 3.02711 2.81713 0.95317 0.48697 0.48576
0.95510
96 2.28814 4.33090 3.17575 2.79628 4.02893 2.93342
3.68262 3.42996 2.81725 3.10060 3.94543 3.14353 3.74413
3.15540 3.18634 2.08022 1.68675 3.03391 5.37129 4.11214 113
- -
2.68618 4.42225 2.77519 2.73123 3.46354 2.40513
3.72494 3.29354 2.67741 2.69355 4.24690 2.90347 2.73739
3.18146 2.89801 2.37887 2.77519 2.98518 4.58477 3.61503
0.02843 3.97018 4.69253 0.61958 0.77255 0.45411
1.00788
97 2.74140 4.58369 3.29463 3.10313 4.53895 0.68244
4.40968 4.09340 3.42546 3.75580 4.69783 3.50567 3.94887
3.74626 3.70160 2.89623 3.21533 3.60931 5.67744 4.61396 114
- -
2.68618 4.42225 2.77519 2.73123 3.46354 2.40513
3.72494 3.29354 2.67741 2.69355 4.24690 2.90347 2.73739
3.18146 2.89801 2.37887 2.77519 2.98518 4.58477 3.61503
0.02696 4.02262 4.74496 0.61958 0.77255 0.48576
0.95510
98 2.87240 5.16455 2.39918 1.12397 4.53346 3.27752
3.81279 3.98750 2.69176 3.58874 4.47755 2.84449 3.88136
2.99498 3.14617 2.66603 3.17787 3.61504 5.75291 4.34606 115
- -
2.68618 4.42225 2.77519 2.73123 3.46354 2.40513
3.72494 3.29354 2.67741 2.69355 4.24690 2.90347 2.73739
3.18146 2.89801 2.37887 2.77519 2.98518 4.58477 3.61503
0.02696 4.02262 4.74496 0.61958 0.77255 0.48576
0.95510
99 3.18120 5.00542 3.68780 3.17038 4.39834 3.63746
3.94268 3.95237 2.31116 3.44111 4.43934 3.51762 4.13451
3.15727 0.77162 3.25933 3.43807 3.67032 5.42160 4.29519 116
- -
2.68618 4.42225 2.77519 2.73123 3.46354 2.40513
3.72494 3.29354 2.67741 2.69355 4.24690 2.90347 2.73739
3.18146 2.89801 2.37887 2.77519 2.98518 4.58477 3.61503
0.02696 4.02262 4.74496 0.61958 0.77255 0.48576
0.95510

100 3.45487 4.81335 4.44471 4.18978 0.79468 4.10525
3.95198 2.96241 4.09358 2.29884 3.67170 4.19112 4.56204
4.22068 4.17611 3.71567 3.75688 2.96378 4.16042 2.51021 117
- -
2.68618 4.42225 2.77519 2.73123 3.46354 2.40513
3.72494 3.29354 2.67741 2.69355 4.24690 2.90347 2.73739
3.18146 2.89801 2.37887 2.77519 2.98518 4.58477 3.61503
0.02696 4.02262 4.74496 0.61958 0.77255 0.48576
0.95510
101 2.93884 5.28356 2.38424 1.12246 4.56270 3.31397
3.78387 4.05273 2.62255 3.58831 4.48514 2.83271 3.89833
2.77076 3.06132 2.87680 3.21804 3.69365 5.75043 4.34836 118
- -
2.68618 4.42225 2.77519 2.73123 3.46354 2.40513
3.72494 3.29354 2.67741 2.69355 4.24690 2.90347 2.73739
3.18146 2.89801 2.37887 2.77519 2.98518 4.58477 3.61503
0.02696 4.02262 4.74496 0.61958 0.77255 0.48576
0.95510
102 2.28395 4.33971 3.28698 3.16844 4.44460 0.91026
4.34967 3.88261 3.36162 3.60445 4.50514 3.38042 3.80439
3.65957 3.64342 2.59640 2.92752 3.35318 5.72734 4.55379 119
- -
2.68618 4.42225 2.77519 2.73123 3.46354 2.40513
3.72494 3.29354 2.67741 2.69355 4.24690 2.90347 2.73739
3.18146 2.89801 2.37887 2.77519 2.98518 4.58477 3.61503
0.02696 4.02262 4.74496 0.61958 0.77255 0.48576
0.95510
103 3.23968 4.56415 4.83033 4.28832 3.21358 4.52719
4.89761 1.61296 4.14381 1.08454 3.01463 4.54523 4.74794
4.31649 4.29619 3.88701 3.47620 1.90106 5.27031 4.15483 120
- -
2.68618 4.42225 2.77519 2.73123 3.46354 2.40513
3.72494 3.29354 2.67741 2.69355 4.24690 2.90347 2.73739
3.18146 2.89801 2.37887 2.77519 2.98518 4.58477 3.61503
0.02696 4.02262 4.74496 0.61958 0.77255 0.48576
0.95510
104 3.24147 4.64085 4.64480 4.13885 3.13357 4.38828
4.75888 2.10411 3.92543 0.87652 2.99067 4.40558 4.67153
4.17255 4.09458 3.77006 3.49695 2.10383 5.17787 3.99817 121
- -
2.68618 4.42225 2.77519 2.73123 3.46354 2.40513
3.72494 3.29354 2.67741 2.69355 4.24690 2.90347 2.73739
3.18146 2.89801 2.37887 2.77519 2.98518 4.58477 3.61503
0.02696 4.02262 4.74496 0.61958 0.77255 0.48576
0.95510
105 2.96212 4.69205 3.58591 3.44827 4.42684 3.37117
4.51157 4.01315 3.51887 3.62279 4.68053 3.73880 0.59042
3.90306 3.74552 3.13480 3.41594 3.64102 5.57002 4.55027 122
- -
2.68618 4.42225 2.77519 2.73123 3.46354 2.40513
3.72494 3.29354 2.67741 2.69355 4.24690 2.90347 2.73739
3.18146 2.89801 2.37887 2.77519 2.98518 4.58477 3.61503
0.02696 4.02262 4.74496 0.61958 0.77255 0.48576
0.95510

```
    106   2.96212  4.69205  3.58591  3.44827  4.42684  3.37117
4.51157  4.01315  3.51887  3.62279  4.68053  3.73880  0.59042
3.90306  3.74552  3.13480  3.41594  3.64102  5.57002  4.55027    123
- -
          2.68618  4.42225  2.77519  2.73123  3.46354  2.40513
3.72494  3.29354  2.67741  2.69355  4.24690  2.90347  2.73739
3.18146  2.89801  2.37887  2.77519  2.98518  4.58477  3.61503
          0.02696  4.02262  4.74496  0.61958  0.77255  0.48576
0.95510
    107   2.31968  4.31660  4.27526  3.75994  3.57402  3.97700
4.54479  1.86385  3.65554  2.19576  3.43669  4.01686  4.39680
3.95243  3.91027  3.33030  3.12713  1.20795  5.32010  4.10529    124
- -
          2.68618  4.42225  2.77519  2.73123  3.46354  2.40513
3.72494  3.29354  2.67741  2.69355  4.24690  2.90347  2.73739
3.18146  2.89801  2.37887  2.77519  2.98518  4.58477  3.61503
          0.02696  4.02262  4.74496  0.61958  0.77255  0.48576
0.95510
    108   2.00555  3.64933  3.98160  3.49800  3.63901  3.34810
4.25408  2.50075  3.40042  2.60819  3.58499  3.62731  3.95486
3.67222  3.64316  2.55010  2.81787  1.43376  5.17237  3.97177    125
- -
          2.68618  4.42225  2.77519  2.73123  3.46354  2.40513
3.72494  3.29354  2.67741  2.69355  4.24690  2.90347  2.73739
3.18146  2.89801  2.37887  2.77519  2.98518  4.58477  3.61503
          0.02696  4.02262  4.74496  0.61958  0.77255  0.48576
0.95510
    109   2.13202  4.50055  3.16694  2.54177  3.71174  3.43871
3.70208  3.05372  2.58708  2.72177  3.46544  3.08381  3.04005
2.92481  3.00350  2.54183  2.54965  2.75430  5.08100  3.68158    126
- -
          2.68618  4.42225  2.77519  2.73123  3.46354  2.40513
3.72494  3.29354  2.67741  2.69355  4.24690  2.90347  2.73739
3.18146  2.89801  2.37887  2.77519  2.98518  4.58477  3.61503
          0.02696  4.02262  4.74496  0.61958  0.77255  0.48576
0.95510
    110   1.20300  4.16727  3.41926  3.17206  4.32646  1.99262
4.26035  3.70110  3.26544  3.42843  4.26362  3.31681  3.70147
3.54192  3.56976  2.34877  2.72401  3.15937  5.67735  4.46922    127
- -
          2.68618  4.42225  2.77519  2.73123  3.46354  2.40513
3.72494  3.29354  2.67741  2.69355  4.24690  2.90347  2.73739
3.18146  2.89801  2.37887  2.77519  2.98518  4.58477  3.61503
          0.02696  4.02262  4.74496  0.61958  0.77255  0.48576
0.95510
    111   2.96212  4.69205  3.58591  3.44827  4.42684  3.37117
4.51157  4.01315  3.51887  3.62279  4.68053  3.73880  0.59042
3.90306  3.74552  3.13480  3.41594  3.64102  5.57002  4.55027    128
- -
          2.68618  4.42225  2.77519  2.73123  3.46354  2.40513
3.72494  3.29354  2.67741  2.69355  4.24690  2.90347  2.73739
3.18146  2.89801  2.37887  2.77519  2.98518  4.58477  3.61503
          0.02696  4.02262  4.74496  0.61958  0.77255  0.48576
0.95510
```

```
     112   1.64158  2.87951  3.77264  3.28641  3.74519  3.14364
4.12495  3.01570  3.21488  2.81578  3.70406  3.43437  3.79244
3.48786  3.49682  2.20493  2.33512  2.44775  5.19129  3.99235     129
- -
           2.68618  4.42225  2.77519  2.73123  3.46354  2.40513
3.72494  3.29354  2.67741  2.69355  4.24690  2.90347  2.73739
3.18146  2.89801  2.37887  2.77519  2.98518  4.58477  3.61503
           0.02696  4.02262  4.74496  0.61958  0.77255  0.48576
0.95510
     113   3.45487  4.81335  4.44471  4.18978  0.79468  4.10525
3.95198  2.96241  4.09358  2.29884  3.67170  4.19112  4.56204
4.22068  4.17611  3.71567  3.75688  2.96378  4.16042  2.51021     130
- -
           2.68618  4.42225  2.77519  2.73123  3.46354  2.40513
3.72494  3.29354  2.67741  2.69355  4.24690  2.90347  2.73739
3.18146  2.89801  2.37887  2.77519  2.98518  4.58477  3.61503
           0.02696  4.02262  4.74496  0.61958  0.77255  0.48576
0.95510
     114   1.10765  4.14602  3.52909  3.23897  4.24736  2.95997
4.26460  3.54206  3.25943  3.32465  4.18178  3.35509  3.71199
3.55564  3.54801  2.05437  2.54140  3.05252  5.62293  4.41973     131
- -
           2.68618  4.42225  2.77519  2.73123  3.46354  2.40513
3.72494  3.29354  2.67741  2.69355  4.24690  2.90347  2.73739
3.18146  2.89801  2.37887  2.77519  2.98518  4.58477  3.61503
           0.02696  4.02262  4.74496  0.61958  0.77255  0.48576
0.95510
     115   3.19003  4.50278  4.83006  4.30409  3.34729  4.51744
4.94767  1.18640  4.16160  1.53232  3.15764  4.54821  4.76672
4.37976  4.33261  3.88731  3.43647  1.73357  5.37013  4.20960     132
- -
           2.68618  4.42225  2.77519  2.73123  3.46354  2.40513
3.72494  3.29354  2.67741  2.69355  4.24690  2.90347  2.73739
3.18146  2.89801  2.37887  2.77519  2.98518  4.58477  3.61503
           0.02696  4.02262  4.74496  0.61958  0.77255  0.48576
0.95510
     116   3.18120  5.00542  3.68780  3.17038  4.39834  3.63746
3.94268  3.95237  2.31116  3.44111  4.43934  3.51762  4.13451
3.15727  0.77162  3.25933  3.43807  3.67032  5.42160  4.29519     133
- -
           2.68618  4.42225  2.77519  2.73123  3.46354  2.40513
3.72494  3.29354  2.67741  2.69355  4.24690  2.90347  2.73739
3.18146  2.89801  2.37887  2.77519  2.98518  4.58477  3.61503
           0.02696  4.02262  4.74496  0.61958  0.77255  0.48576
0.95510
     117   3.04481  5.12346  3.32454  2.79158  4.54915  3.61845
3.68110  3.93741  1.04397  3.41560  4.30981  3.20479  4.03898
2.83125  2.11594  3.05269  3.24275  3.62293  5.46930  4.28059     134
- -
           2.68618  4.42225  2.77519  2.73123  3.46354  2.40513
3.72494  3.29354  2.67741  2.69355  4.24690  2.90347  2.73739
3.18146  2.89801  2.37887  2.77519  2.98518  4.58477  3.61503
           0.02696  4.02262  4.74496  0.61958  0.77255  0.48576
0.95510
```

```
    118   2.71376  4.85168  3.02402  2.54322  4.18588  3.43934
3.51627  3.60062  2.20246  3.17450  4.01212  3.02248  2.06530
2.81759  2.23169  2.66462  2.95395  3.27258  5.36260  4.07380    135
- -
          2.68618  4.42225  2.77519  2.73123  3.46354  2.40513
3.72494  3.29354  2.67741  2.69355  4.24690  2.90347  2.73739
3.18146  2.89801  2.37887  2.77519  2.98518  4.58477  3.61503
          0.02696  4.02262  4.74496  0.61958  0.77255  0.48576
0.95510
    119   0.81728  4.27297  3.62041  3.42136  4.20706  3.09296
4.42492  3.35544  3.44162  3.22989  4.24718  3.53130  3.84663
3.76944  3.68019  2.62296  2.91086  2.98372  5.61916  4.43687    136
- -
          2.68618  4.42225  2.77519  2.73123  3.46354  2.40513
3.72494  3.29354  2.67741  2.69355  4.24690  2.90347  2.73739
3.18146  2.89801  2.37887  2.77519  2.98518  4.58477  3.61503
          0.02696  4.02262  4.74496  0.61958  0.77255  0.48576
0.95510
    120   2.50248  4.18529  3.70776  3.14478  3.22355  3.56819
3.93459  2.33797  3.06228  2.29683  3.30514  3.47660  4.00783
3.18428  3.35220  2.63381  2.82923  1.73787  4.83230  3.61500    137
- -
          2.68618  4.42225  2.77519  2.73123  3.46354  2.40513
3.72494  3.29354  2.67741  2.69355  4.24690  2.90347  2.73739
3.18146  2.89801  2.37887  2.77519  2.98518  4.58477  3.61503
          0.02696  4.02262  4.74496  0.61958  0.77255  0.48576
0.95510
    121   1.96173  4.53601  3.14185  2.59872  3.80683  2.93498
3.70327  3.19221  2.51487  2.76719  3.49472  3.01024  3.82746
2.90970  2.77694  2.48230  2.80057  2.86249  5.14942  3.86638    138
- -
          2.68618  4.42225  2.77519  2.73123  3.46354  2.40513
3.72494  3.29354  2.67741  2.69355  4.24690  2.90347  2.73739
3.18146  2.89801  2.37887  2.77519  2.98518  4.58477  3.61503
          0.02696  4.02262  4.74496  0.61958  0.77255  0.48576
0.95510
    122   2.80053  4.28588  4.15671  3.62202  3.45066  3.92299
4.36999  1.81150  3.51225  2.17258  3.34900  3.89814  3.44705
3.80092  3.76245  3.25008  3.06496  1.31665  5.13956  3.92923    139
- -
          2.68618  4.42225  2.77519  2.73123  3.46354  2.40513
3.72494  3.29354  2.67741  2.69355  4.24690  2.90347  2.73739
3.18146  2.89801  2.37887  2.77519  2.98518  4.58477  3.61503
          0.02696  4.02262  4.74496  0.61958  0.77255  0.48576
0.95510
    123   2.90771  4.34739  4.19218  3.66934  1.56893  3.92724
3.41280  2.23297  3.53737  2.25267  3.37167  3.79240  4.27181
3.71450  3.70451  3.22844  3.13584  2.58391  4.11860  2.35511    140
- -
          2.68618  4.42225  2.77519  2.73123  3.46354  2.40513
3.72494  3.29354  2.67741  2.69355  4.24690  2.90347  2.73739
3.18146  2.89801  2.37887  2.77519  2.98518  4.58477  3.61503
          0.02696  4.02262  4.74496  0.61958  0.77255  0.48576
0.95510
```

124 2.22352 4.35819 3.26663 2.79107 3.96812 2.89434
3.88316 3.35665 2.77947 3.02825 3.69045 3.10867 3.03412
3.10897 3.16965 2.39585 1.76523 2.98763 5.31468 4.05389 141
- -
2.68618 4.42225 2.77519 2.73123 3.46354 2.40513
3.72494 3.29354 2.67741 2.69355 4.24690 2.90347 2.73739
3.18146 2.89801 2.37887 2.77519 2.98518 4.58477 3.61503
0.02696 4.02262 4.74496 0.61958 0.77255 0.48576
0.95510
125 3.28700 4.68723 4.60986 4.13576 3.09929 4.38586
4.72763 2.12651 3.89188 0.82702 2.99679 4.40310 4.67769
4.16657 4.05383 3.79789 3.54357 2.27778 5.13226 3.91227 142
- -
2.68618 4.42225 2.77519 2.73123 3.46354 2.40513
3.72494 3.29354 2.67741 2.69355 4.24690 2.90347 2.73739
3.18146 2.89801 2.37887 2.77519 2.98518 4.58477 3.61503
0.02696 4.02262 4.74496 0.61958 0.77255 0.48576
0.95510
126 2.37076 5.19624 1.94110 2.00711 4.49750 3.22083
3.59647 3.97530 2.49969 3.48861 4.27793 2.58374 3.79886
2.63938 3.02216 2.49147 2.94990 3.56886 5.65764 4.22997 143
- -
2.68618 4.42225 2.77519 2.73123 3.46354 2.40513
3.72494 3.29354 2.67741 2.69355 4.24690 2.90347 2.73739
3.18146 2.89801 2.37887 2.77519 2.98518 4.58477 3.61503
0.02696 4.02262 4.74496 0.61958 0.77255 0.48576
0.95510
127 2.72767 5.17971 1.52695 2.24587 4.49888 3.30256
3.69017 3.97896 2.51918 3.50774 4.32425 2.73770 3.82984
2.80546 2.45935 2.73079 3.04205 3.58530 5.66960 4.25696 144
- -
2.68618 4.42225 2.77519 2.73123 3.46354 2.40513
3.72494 3.29354 2.67741 2.69355 4.24690 2.90347 2.73739
3.18146 2.89801 2.37887 2.77519 2.98518 4.58477 3.61503
0.02696 4.02262 4.74496 0.61958 0.77255 0.48576
0.95510
128 3.47032 4.89825 4.06283 3.82242 2.31343 4.00733
3.69832 3.45928 3.67573 2.87390 4.11913 3.91273 4.47865
3.94536 3.82775 3.59122 3.75827 3.33721 3.96223 0.74848 145
- -
2.68618 4.42225 2.77519 2.73123 3.46354 2.40513
3.72494 3.29354 2.67741 2.69355 4.24690 2.90347 2.73739
3.18146 2.89801 2.37887 2.77519 2.98518 4.58477 3.61503
0.02696 4.02262 4.74496 0.61958 0.77255 0.48576
0.95510
129 2.84338 4.37881 4.51463 4.04635 3.61901 4.13984
4.81976 1.74356 3.91852 2.19866 3.45714 4.26155 4.56425
4.22974 4.14497 3.53584 3.25747 0.98864 5.49463 4.26683 146
- -
2.68618 4.42225 2.77519 2.73123 3.46354 2.40513
3.72494 3.29354 2.67741 2.69355 4.24690 2.90347 2.73739
3.18146 2.89801 2.37887 2.77519 2.98518 4.58477 3.61503
0.02696 4.02262 4.74496 0.61958 0.77255 0.48576
0.95510

```
    130   1.99628  4.78509  2.76794  2.29789  4.04057  3.38823
3.56230  3.39105  2.42328  3.05996  3.87564  2.83216  3.79770
2.77524  2.73992  2.54714  2.77462  2.99407  5.30250  3.96740    147
- -
          2.68618  4.42225  2.77519  2.73123  3.46354  2.40513
3.72494  3.29354  2.67741  2.69355  4.24690  2.90347  2.73739
3.18146  2.89801  2.37887  2.77519  2.98518  4.58477  3.61503
          0.02696  4.02262  4.74496  0.61958  0.77255  0.48576
0.95510
    131   1.95523  5.02157  2.06482  2.24528  4.34819  3.30602
3.67414  3.79200  2.52572  3.35635  4.16645  2.82708  3.80557
2.53938  3.02271  2.59952  2.88465  3.41743  5.56149  4.17263    148
- -
          2.68618  4.42225  2.77519  2.73123  3.46354  2.40513
3.72494  3.29354  2.67741  2.69355  4.24690  2.90347  2.73739
3.18146  2.89801  2.37887  2.77519  2.98518  4.58477  3.61503
          0.02696  4.02262  4.74496  0.61958  0.77255  0.48576
0.95510
    132   2.51310  4.76602  2.67222  2.40602  4.24302  1.69815
3.76543  3.66299  2.58014  3.27025  4.10691  2.91371  3.82079
2.94411  2.70076  2.67406  2.93474  3.29353  5.49558  4.17032    149
- -
          2.68618  4.42225  2.77519  2.73123  3.46354  2.40513
3.72494  3.29354  2.67741  2.69355  4.24690  2.90347  2.73739
3.18146  2.89801  2.37887  2.77519  2.98518  4.58477  3.61503
          0.02696  4.02262  4.74496  0.61958  0.77255  0.48576
0.95510
    133   2.81289  4.25577  4.26395  3.70905  3.37540  3.97629
4.37541  1.38820  3.58650  2.12835  2.80316  3.95915  4.33574
3.85056  3.80495  3.09585  2.95165  1.80418  5.06904  3.87674    150
- -
          2.68618  4.42225  2.77519  2.73123  3.46354  2.40513
3.72494  3.29354  2.67741  2.69355  4.24690  2.90347  2.73739
3.18146  2.89801  2.37887  2.77519  2.98518  4.58477  3.61503
          0.02696  4.02262  4.74496  0.61958  0.77255  0.48576
0.95510
    134   3.20687  4.59535  4.62664  4.06447  3.10281  4.39404
4.68800  2.12086  3.89636  1.29935  1.60876  4.35357  4.62613
4.09280  4.06808  3.72870  3.43479  2.24233  5.12496  4.01622    151
- -
          2.68618  4.42225  2.77519  2.73123  3.46354  2.40513
3.72494  3.29354  2.67741  2.69355  4.24690  2.90347  2.73739
3.18146  2.89801  2.37887  2.77519  2.98518  4.58477  3.61503
          0.02696  4.02262  4.74496  0.61958  0.77255  0.48576
0.95510
    135   2.44054  4.56869  3.03964  2.54197  3.74133  3.35910
3.68746  3.17116  2.54158  2.79578  3.71039  3.03919  3.76853
2.79014  2.92623  2.33935  2.10346  2.89832  5.15802  3.86654    152
- -
          2.68618  4.42225  2.77519  2.73123  3.46354  2.40513
3.72494  3.29354  2.67741  2.69355  4.24690  2.90347  2.73739
3.18146  2.89801  2.37887  2.77519  2.98518  4.58477  3.61503
          0.02696  4.02262  4.74496  0.61958  0.77255  0.48576
0.95510
```

```
    136   1.91982  4.73368  2.81825  2.34527  4.03372  3.36179
3.65584  3.44597  2.47390  3.05932  3.88132  2.94793  3.38686
2.79315  2.83078  2.45717  2.75284  3.04307  5.30989  3.98275    153
- -
          2.68618  4.42225  2.77519  2.73123  3.46354  2.40513
3.72494  3.29354  2.67741  2.69355  4.24690  2.90347  2.73739
3.18146  2.89801  2.37887  2.77519  2.98518  4.58477  3.61503
          0.02696  4.02262  4.74496  0.61958  0.77255  0.48576
0.95510
    137   2.24661  4.54238  2.85072  2.28302  4.02119  2.94611
3.60638  3.43839  2.36413  2.99212  3.70721  2.93094  3.69679
2.73173  2.71012  2.59210  2.67775  3.04616  5.28182  3.94627    154
- -
          2.68618  4.42225  2.77519  2.73123  3.46354  2.40513
3.72494  3.29354  2.67741  2.69355  4.24690  2.90347  2.73739
3.18146  2.89801  2.37887  2.77519  2.98518  4.58477  3.61503
          0.02696  4.02262  4.74496  0.61958  0.77255  0.48576
0.95510
    138   2.54168  4.95791  2.77163  2.32179  4.27340  3.38731
3.51906  3.69538  2.36443  3.26234  4.09312  2.92557  3.85186
1.69990  2.75052  2.70673  2.83386  3.35279  5.45743  4.12706    155
- -
          2.68618  4.42225  2.77519  2.73123  3.46354  2.40513
3.72494  3.29354  2.67741  2.69355  4.24690  2.90347  2.73739
3.18146  2.89801  2.37887  2.77519  2.98518  4.58477  3.61503
          0.02696  4.02262  4.74496  0.61958  0.77255  0.48576
0.95510
    139   1.01938  4.25231  3.30658  3.08348  4.22286  2.97446
4.19946  3.54735  3.15840  3.33727  4.21571  3.00083  3.75767
3.48451  3.46025  2.50119  2.79807  3.09128  5.59350  4.34834    156
- -
          2.68618  4.42225  2.77519  2.73123  3.46354  2.40513
3.72494  3.29354  2.67741  2.69355  4.24690  2.90347  2.73739
3.18146  2.89801  2.37887  2.77519  2.98518  4.58477  3.61503
          0.02696  4.02262  4.74496  0.61958  0.77255  0.48576
0.95510
    140   2.35005  5.00569  2.33006  2.06359  4.30733  2.96858
3.60787  3.76058  2.39371  3.30284  4.08770  2.83611  3.78036
2.69744  2.85836  2.42731  2.83475  3.22593  5.49040  4.10339    157
- -
          2.68618  4.42225  2.77519  2.73123  3.46354  2.40513
3.72494  3.29354  2.67741  2.69355  4.24690  2.90347  2.73739
3.18146  2.89801  2.37887  2.77519  2.98518  4.58477  3.61503
          0.02696  4.02262  4.74496  0.61958  0.77255  0.48576
0.95510
    141   2.21356  4.11875  3.91952  3.34391  2.50129  3.68932
3.99137  2.25213  3.24637  2.08080  3.19618  3.61267  4.05898
3.49297  3.47853  2.91790  2.71708  2.14215  4.74229  3.43428    158
- -
          2.68618  4.42225  2.77519  2.73123  3.46354  2.40513
3.72494  3.29354  2.67741  2.69355  4.24690  2.90347  2.73739
3.18146  2.89801  2.37887  2.77519  2.98518  4.58477  3.61503
          0.02696  4.02262  4.74496  0.61958  0.77255  0.48576
0.95510
```

```
    142   3.27669  4.70308  4.53056  4.08115  3.09546  4.31347
4.67589  2.26615  3.81016  0.80315  2.95886  4.34544  4.63914
4.12099  3.97320  3.74867  3.54303  2.35614  5.10714  3.86687    159
- -
          2.68618  4.42225  2.77519  2.73123  3.46354  2.40513
3.72494  3.29354  2.67741  2.69355  4.24690  2.90347  2.73739
3.18146  2.89801  2.37887  2.77519  2.98518  4.58477  3.61503
          0.02696  4.02262  4.74496  0.61958  0.77255  0.48576
0.95510
    143   2.49352  4.11169  3.24134  2.66319  3.79189  3.47228
3.64977  3.08723  2.45385  2.81778  3.68846  3.11714  3.87423
2.80121  1.90656  2.72390  2.84836  2.64182  5.12388  3.85821    160
- -
          2.68618  4.42225  2.77519  2.73123  3.46354  2.40513
3.72494  3.29354  2.67741  2.69355  4.24690  2.90347  2.73739
3.18146  2.89801  2.37887  2.77519  2.98518  4.58477  3.61503
          0.02696  4.02262  4.74496  0.61958  0.77255  0.48576
0.95510
    144   2.44458  4.46018  2.79365  2.30427  4.11989  3.25868
3.51466  3.50701  2.32878  2.98160  3.86476  2.84923  3.78310
2.57255  2.30267  2.59478  2.81173  3.16802  5.33947  3.98980    161
- -
          2.68618  4.42225  2.77519  2.73123  3.46354  2.40513
3.72494  3.29354  2.67741  2.69355  4.24690  2.90347  2.73739
3.18146  2.89801  2.37887  2.77519  2.98518  4.58477  3.61503
          0.02696  4.02262  4.74496  0.61958  0.77255  0.48576
0.95510
    145   1.00844  4.18463  3.46597  3.23808  4.31702  2.36946
4.30669  3.64577  3.31963  3.41393  4.27803  3.36765  3.73054
3.60562  3.60371  2.45380  2.76392  3.13602  5.66954  4.47893    162
- -
          2.68618  4.42225  2.77519  2.73123  3.46354  2.40513
3.72494  3.29354  2.67741  2.69355  4.24690  2.90347  2.73739
3.18146  2.89801  2.37887  2.77519  2.98518  4.58477  3.61503
          0.02696  4.02262  4.74496  0.61958  0.77255  0.48576
0.95510
    146   3.03454  4.41781  4.61544  4.12674  3.55141  4.26975
4.86759  1.73697  3.98028  2.09822  3.32328  4.35454  4.64406
4.28626  4.19828  3.65866  3.31802  0.97242  5.47279  4.24020    163
- -
          2.68618  4.42225  2.77519  2.73123  3.46354  2.40513
3.72494  3.29354  2.67741  2.69355  4.24690  2.90347  2.73739
3.18146  2.89801  2.37887  2.77519  2.98518  4.58477  3.61503
          0.02696  4.02262  4.74496  0.61958  0.77255  0.48576
0.95510
    147   2.18453  4.64369  3.04915  2.49845  3.85323  3.39677
3.27566  3.16404  2.43237  2.88787  3.57110  3.00933  3.82576
2.80855  2.44472  2.58611  2.81053  2.80490  5.16823  3.81737    164
- -
          2.68618  4.42225  2.77519  2.73123  3.46354  2.40513
3.72494  3.29354  2.67741  2.69355  4.24690  2.90347  2.73739
3.18146  2.89801  2.37887  2.77519  2.98518  4.58477  3.61503
          0.02696  4.02262  4.74496  0.61958  0.77255  0.48576
0.95510
```

148 2.42477 5.12404 2.29203 1.76670 4.43970 3.08702
3.64099 3.90871 2.47737 3.43305 4.22123 2.63709 3.79271
2.77263 2.95876 2.50973 2.80557 3.50821 5.60855 4.19525 165
- -
2.68618 4.42225 2.77519 2.73123 3.46354 2.40513
3.72494 3.29354 2.67741 2.69355 4.24690 2.90347 2.73739
3.18146 2.89801 2.37887 2.77519 2.98518 4.58477 3.61503
0.02696 4.02262 4.74496 0.61958 0.77255 0.48576
0.95510
149 3.04214 5.10914 3.43301 2.81133 4.44199 3.66346
3.31765 3.88614 1.87400 3.35537 4.23513 3.21148 4.04454
2.78605 1.19603 3.05314 3.21768 3.58093 5.39235 4.18518 166
- -
2.68618 4.42225 2.77519 2.73123 3.46354 2.40513
3.72494 3.29354 2.67741 2.69355 4.24690 2.90347 2.73739
3.18146 2.89801 2.37887 2.77519 2.98518 4.58477 3.61503
0.02696 4.02262 4.74496 0.61958 0.77255 0.48576
0.95510
150 2.31083 4.93962 2.93806 2.40822 4.23487 3.42972
3.46622 3.65721 2.09342 3.06186 4.00571 2.91569 3.82102
2.29352 2.34990 2.63913 2.87883 3.30603 5.38625 4.05703 167
- -
2.68618 4.42225 2.77519 2.73123 3.46354 2.40513
3.72494 3.29354 2.67741 2.69355 4.24690 2.90347 2.73739
3.18146 2.89801 2.37887 2.77519 2.98518 4.58477 3.61503
0.02696 4.02262 4.74496 0.61958 0.77255 0.48576
0.95510
151 2.85426 4.85176 2.80169 2.70353 4.23951 3.28976
4.05172 3.98473 2.96940 3.61516 4.57129 0.91734 3.95002
3.32667 3.32750 2.91694 3.24227 3.59163 5.52835 4.17856 168
- -
2.68618 4.42225 2.77519 2.73123 3.46354 2.40513
3.72494 3.29354 2.67741 2.69355 4.24690 2.90347 2.73739
3.18146 2.89801 2.37887 2.77519 2.98518 4.58477 3.61503
0.02696 4.02262 4.74496 0.61958 0.77255 0.48576
0.95510
152 3.12892 4.44458 4.81217 4.31197 3.51780 4.47015
5.01554 1.09984 4.18591 1.97019 3.32600 4.53903 4.76984
4.44670 4.37987 3.85642 3.39226 1.44382 5.50729 4.30911 169
- -
2.68618 4.42225 2.77519 2.73123 3.46354 2.40513
3.72494 3.29354 2.67741 2.69355 4.24690 2.90347 2.73739
3.18146 2.89801 2.37887 2.77519 2.98518 4.58477 3.61503
0.02696 4.02262 4.74496 0.61958 0.77255 0.48576
0.95510
153 3.19779 4.62134 4.56816 4.09107 3.16038 4.29710
4.72639 2.16792 3.88196 0.89667 3.03277 4.34722 4.62517
4.14718 4.05476 3.69942 3.46738 2.03918 5.17879 3.98466 170
- -
2.68618 4.42225 2.77519 2.73123 3.46354 2.40513
3.72494 3.29354 2.67741 2.69355 4.24690 2.90347 2.73739
3.18146 2.89801 2.37887 2.77519 2.98518 4.58477 3.61503
0.02696 4.02262 4.74496 0.61958 0.77255 0.48576
0.95510

```
    154   3.09348  4.41609  4.78567  4.29381  3.58255  4.42722
5.01586  1.40640  4.17989  2.09231  3.39199  4.51264  4.75084
4.45115  4.38221  3.81760  3.36313  1.08058  5.54385  4.33301    171
- -
          2.68618  4.42225  2.77519  2.73123  3.46354  2.40513
3.72494  3.29354  2.67741  2.69355  4.24690  2.90347  2.73739
3.18146  2.89801  2.37887  2.77519  2.98518  4.58477  3.61503
          0.02696  4.02262  4.74496  0.61958  0.77255  0.48576
0.95510
    155   1.14448  4.16385  3.68284  3.32025  4.03703  3.08296
4.27056  3.04497  3.26685  3.01302  3.96465  3.44470  3.79637
3.58119  3.54652  2.48522  2.59350  2.44682  5.49798  4.28909    172
- -
          2.68618  4.42225  2.77519  2.73123  3.46354  2.40513
3.72494  3.29354  2.67741  2.69355  4.24690  2.90347  2.73739
3.18146  2.89801  2.37887  2.77519  2.98518  4.58477  3.61503
          0.02696  4.02262  4.74496  0.61958  0.77255  0.48576
0.95510
    156   2.86181  4.62005  3.56615  3.48578  4.62096  0.53093
4.59479  4.25913  3.68734  3.89379  4.87234  3.73076  4.01589
3.99980  3.90487  3.03839  3.35475  3.75848  5.67431  4.72868    173
- -
          2.68618  4.42225  2.77519  2.73123  3.46354  2.40513
3.72494  3.29354  2.67741  2.69355  4.24690  2.90347  2.73739
3.18146  2.89801  2.37887  2.77519  2.98518  4.58477  3.61503
          0.02696  4.02262  4.74496  0.61958  0.77255  0.48576
0.95510
    157   1.96574  4.23718  3.26384  3.06259  4.39116  1.22970
4.23059  3.79372  3.21660  3.49103  4.32962  3.27144  3.33965
3.49868  3.53762  2.45067  2.77267  3.24214  5.71482  4.50007    174
- -
          2.68618  4.42225  2.77519  2.73123  3.46354  2.40513
3.72494  3.29354  2.67741  2.69355  4.24690  2.90347  2.73739
3.18146  2.89801  2.37887  2.77519  2.98518  4.58477  3.61503
          0.02696  4.02262  4.74496  0.61958  0.77255  0.48576
0.95510
    158   2.49976  4.34431  3.56805  3.33000  4.10800  3.17788
4.33500  3.25370  3.27593  3.11180  4.15459  3.51326  3.89436
3.66375  3.52580  2.70296  0.94063  2.93233  5.53249  4.32538    175
- -
          2.68618  4.42225  2.77519  2.73123  3.46354  2.40513
3.72494  3.29354  2.67741  2.69355  4.24690  2.90347  2.73739
3.18146  2.89801  2.37887  2.77519  2.98518  4.58477  3.61503
          0.02696  4.02262  4.74496  0.61958  0.77255  0.48576
0.95510
    159   2.29660  4.25470  3.26092  3.06305  4.37522  2.53496
4.22530  3.85825  3.20034  3.53542  4.37056  3.27425  3.72648
3.49775  3.51644  1.03762  2.78809  3.28834  5.69711  4.45772    176
- -
          2.68618  4.42225  2.77519  2.73123  3.46354  2.40513
3.72494  3.29354  2.67741  2.69355  4.24690  2.90347  2.73739
3.18146  2.89801  2.37887  2.77519  2.98518  4.58477  3.61503
          0.02696  4.02262  4.74496  0.61958  0.77255  0.48576
0.95510
```

160 2.20241 4.20753 3.42640 3.09539 4.19452 3.01138
4.15578 3.52333 3.10202 3.26886 4.12309 3.29550 3.72813
3.42231 3.42068 1.72686 1.51419 3.06026 5.55604 4.33069 177
- -
2.68618 4.42225 2.77519 2.73123 3.46354 2.40513
3.72494 3.29354 2.67741 2.69355 4.24690 2.90347 2.73739
3.18146 2.89801 2.37887 2.77519 2.98518 4.58477 3.61503
0.02696 4.02262 4.74496 0.61958 0.77255 0.48576
0.95510
161 2.86181 4.62005 3.56615 3.48578 4.62096 0.53093
4.59479 4.25913 3.68734 3.89379 4.87234 3.73076 4.01589
3.99980 3.90487 3.03839 3.35475 3.75848 5.67431 4.72868 178
- -
2.68618 4.42225 2.77519 2.73123 3.46354 2.40513
3.72494 3.29354 2.67741 2.69355 4.24690 2.90347 2.73739
3.18146 2.89801 2.37887 2.77519 2.98518 4.58477 3.61503
0.02696 4.02262 4.74496 0.61958 0.77255 0.48576
0.95510
162 3.11448 5.07907 3.30011 2.89647 4.49993 3.58511
3.85036 3.93922 0.84983 3.47273 4.42835 3.31347 4.07671
3.03496 2.47086 3.14796 3.35771 3.63935 5.49460 4.33003 179
- -
2.68618 4.42225 2.77519 2.73123 3.46354 2.40513
3.72494 3.29354 2.67741 2.69355 4.24690 2.90347 2.73739
3.18146 2.89801 2.37887 2.77519 2.98518 4.58477 3.61503
0.02696 4.02262 4.74496 0.61958 0.77255 0.48576
0.95510
163 2.49976 4.34431 3.56805 3.33000 4.10800 3.17788
4.33500 3.25370 3.27593 3.11180 4.15459 3.51326 3.89436
3.66375 3.52580 2.70296 0.94063 2.93233 5.53249 4.32538 180
- -
2.68618 4.42225 2.77519 2.73123 3.46354 2.40513
3.72494 3.29354 2.67741 2.69355 4.24690 2.90347 2.73739
3.18146 2.89801 2.37887 2.77519 2.98518 4.58477 3.61503
0.02696 4.02262 4.74496 0.61958 0.77255 0.48576
0.95510
164 2.49976 4.34431 3.56805 3.33000 4.10800 3.17788
4.33500 3.25370 3.27593 3.11180 4.15459 3.51326 3.89436
3.66375 3.52580 2.70296 0.94063 2.93233 5.53249 4.32538 181
- -
2.68618 4.42225 2.77519 2.73123 3.46354 2.40513
3.72494 3.29354 2.67741 2.69355 4.24690 2.90347 2.73739
3.18146 2.89801 2.37887 2.77519 2.98518 4.58477 3.61503
0.02696 4.02262 4.74496 0.61958 0.77255 0.48576
0.95510
165 3.33472 4.70480 4.67585 4.19023 2.64262 4.44145
4.59352 2.27986 4.03098 0.82562 2.96078 4.42630 4.69334
4.19265 4.17392 3.83352 3.57422 2.40018 4.89527 3.55683 182
- -
2.68618 4.42225 2.77519 2.73123 3.46354 2.40513
3.72494 3.29354 2.67741 2.69355 4.24690 2.90347 2.73739
3.18146 2.89801 2.37887 2.77519 2.98518 4.58477 3.61503
0.02696 4.02262 4.74496 0.61958 0.77255 0.48576
0.95510

166 1.80169 4.20262 3.70877 3.25052 3.80703 3.26274
4.17245 2.72083 3.18800 2.75607 3.72601 3.46750 3.88718
3.49520 3.49256 2.65474 1.93794 1.98352 5.30145 4.08652 183
- -
2.68618 4.42225 2.77519 2.73123 3.46354 2.40513
3.72494 3.29354 2.67741 2.69355 4.24690 2.90347 2.73739
3.18146 2.89801 2.37887 2.77519 2.98518 4.58477 3.61503
0.02696 4.02262 4.74496 0.61958 0.77255 0.48576
0.95510
167 2.85426 4.85176 2.80169 2.70353 4.23951 3.28976
4.05172 3.98473 2.96940 3.61516 4.57129 0.91734 3.95002
3.32667 3.32750 2.91694 3.24227 3.59163 5.52835 4.17856 184
- -
2.68618 4.42225 2.77519 2.73123 3.46354 2.40513
3.72494 3.29354 2.67741 2.69355 4.24690 2.90347 2.73739
3.18146 2.89801 2.37887 2.77519 2.98518 4.58477 3.61503
0.02696 4.02262 4.74496 0.61958 0.77255 0.48576
0.95510
168 0.81728 4.27297 3.62041 3.42136 4.20706 3.09296
4.42492 3.35544 3.44162 3.22989 4.24718 3.53130 3.84663
3.76944 3.68019 2.62296 2.91086 2.98372 5.61916 4.43687 185
- -
2.68618 4.42225 2.77519 2.73123 3.46354 2.40513
3.72494 3.29354 2.67741 2.69355 4.24690 2.90347 2.73739
3.18146 2.89801 2.37887 2.77519 2.98518 4.58477 3.61503
0.02696 4.02262 4.74496 0.61958 0.77255 0.48576
0.95510
169 3.27212 4.70198 4.49805 4.07494 3.10839 4.27953
4.67075 2.25113 3.81850 0.79860 3.05236 4.33404 4.62752
4.13153 3.98072 3.73765 3.54596 2.35525 5.10451 3.84781 186
- -
2.68618 4.42225 2.77519 2.73123 3.46354 2.40513
3.72494 3.29354 2.67741 2.69355 4.24690 2.90347 2.73739
3.18146 2.89801 2.37887 2.77519 2.98518 4.58477 3.61503
0.02696 4.02262 4.74496 0.61958 0.77255 0.48576
0.95510
170 3.32709 4.74617 4.42998 4.07280 3.19879 4.17148
4.68160 2.41344 3.83907 0.71022 3.17896 4.33366 4.58250
4.17133 3.98965 3.76330 3.61534 2.48932 5.13017 3.88739 187
- -
2.68618 4.42225 2.77519 2.73123 3.46354 2.40513
3.72494 3.29354 2.67741 2.69355 4.24690 2.90347 2.73739
3.18146 2.89801 2.37887 2.77519 2.98518 4.58477 3.61503
0.02696 4.02262 4.74496 0.61958 0.77255 0.48576
0.95510
171 1.33026 4.49741 2.85604 2.66471 4.05013 3.03476
3.51121 3.47734 2.79721 3.14707 4.00602 3.08320 3.80580
3.12186 3.18982 2.61945 2.86436 3.11056 5.39524 4.09629 188
- -
2.68618 4.42225 2.77519 2.73123 3.46354 2.40513
3.72494 3.29354 2.67741 2.69355 4.24690 2.90347 2.73739
3.18146 2.89801 2.37887 2.77519 2.98518 4.58477 3.61503
0.02696 4.02262 4.74496 0.61958 0.77255 0.48576
0.95510

172 2.88423 5.09939 2.51921 1.10347 4.39405 3.32987
3.83693 3.76707 2.68488 3.41692 4.36703 2.92059 3.91307
3.02552 3.11199 2.87041 3.18214 3.30243 5.65927 4.28131 189
- -
2.68618 4.42225 2.77519 2.73123 3.46354 2.40513
3.72494 3.29354 2.67741 2.69355 4.24690 2.90347 2.73739
3.18146 2.89801 2.37887 2.77519 2.98518 4.58477 3.61503
0.02696 4.02262 4.74496 0.61958 0.77255 0.48576
0.95510
173 2.76777 4.26960 3.77066 3.64091 3.44551 4.00785
4.37559 1.59308 3.55353 2.14559 3.21016 3.93106 4.35262
3.82276 3.80566 3.31286 3.06842 1.44126 5.12282 3.91540 190
- -
2.68618 4.42225 2.77519 2.73123 3.46354 2.40513
3.72494 3.29354 2.67741 2.69355 4.24690 2.90347 2.73739
3.18146 2.89801 2.37887 2.77519 2.98518 4.58477 3.61503
0.02696 4.02262 4.74496 0.61958 0.77255 0.48576
0.95510
174 1.07037 4.14639 3.47667 3.21114 4.29290 2.57273
4.26565 3.64974 3.27179 3.38784 4.22455 3.33439 3.69956
3.55372 3.56621 2.25157 2.63205 3.12114 5.65249 4.45172 191
- -
2.68618 4.42225 2.77519 2.73123 3.46354 2.40513
3.72494 3.29354 2.67741 2.69355 4.24690 2.90347 2.73739
3.18146 2.89801 2.37887 2.77519 2.98518 4.58477 3.61503
0.02696 4.02262 4.74496 0.61958 0.77255 0.48576
0.95510
175 2.24353 4.86570 2.82161 2.36030 3.84453 3.31375
3.59929 3.55950 1.92914 3.13617 3.93924 2.89354 3.79591
2.68336 2.69220 2.61866 2.80143 3.21668 5.35097 4.00705 192
- -
2.68618 4.42225 2.77519 2.73123 3.46354 2.40513
3.72494 3.29354 2.67741 2.69355 4.24690 2.90347 2.73739
3.18146 2.89801 2.37887 2.77519 2.98518 4.58477 3.61503
0.02696 4.02262 4.74496 0.61958 0.77255 0.48576
0.95510
176 2.27603 4.04762 3.35987 2.86377 3.84577 3.02125
3.66681 3.22438 2.81795 2.85584 3.77203 3.20968 3.78904
3.15181 3.18535 2.30453 1.68449 2.88731 5.22278 3.97528 193
- -
2.68618 4.42225 2.77519 2.73123 3.46354 2.40513
3.72494 3.29354 2.67741 2.69355 4.24690 2.90347 2.73739
3.18146 2.89801 2.37887 2.77519 2.98518 4.58477 3.61503
0.02696 4.02262 4.74496 0.61958 0.77255 0.48576
0.95510
177 2.48194 4.88785 2.50491 2.26853 4.08812 2.42580
3.64314 3.64629 2.46443 3.22244 4.02596 2.87467 3.78822
2.72275 2.94688 2.29301 2.71048 3.28411 5.43902 4.07534 194
- -
2.68618 4.42225 2.77519 2.73123 3.46354 2.40513
3.72494 3.29354 2.67741 2.69355 4.24690 2.90347 2.73739
3.18146 2.89801 2.37887 2.77519 2.98518 4.58477 3.61503
0.02696 4.02262 4.74496 0.61958 0.77255 0.48576
0.95510

```
    178   3.01138  5.55566  1.10566  1.85535  4.84546  3.23282
3.82099  4.36701  2.87843  3.88665  4.76707  2.70829  3.87685
2.99945  3.48561  2.88929  3.31173  3.94792  6.03190  4.52873    195
- -
          2.68618  4.42225  2.77519  2.73123  3.46354  2.40513
3.72494  3.29354  2.67741  2.69355  4.24690  2.90347  2.73739
3.18146  2.89801  2.37887  2.77519  2.98518  4.58477  3.61503
          0.02696  4.02262  4.74496  0.61958  0.77255  0.48576
0.95510
    179   3.18120  5.00542  3.68780  3.17038  4.39834  3.63746
3.94268  3.95237  2.31116  3.44111  4.43934  3.51762  4.13451
3.15727  0.77162  3.25933  3.43807  3.67032  5.42160  4.29519    196
- -
          2.68618  4.42225  2.77519  2.73123  3.46354  2.40513
3.72494  3.29354  2.67741  2.69355  4.24690  2.90347  2.73739
3.18146  2.89801  2.37887  2.77519  2.98518  4.58477  3.61503
          0.02696  4.02262  4.74496  0.61958  0.77255  0.48576
0.95510
    180   3.09019  4.42266  4.73426  4.22835  3.54627  4.41171
4.95066  1.58985  4.10462  2.00703  3.36604  4.46434  4.72741
4.38098  4.31438  3.78980  3.35501  1.02151  5.49591  4.28201    197
- -
          2.68618  4.42225  2.77519  2.73123  3.46354  2.40513
3.72494  3.29354  2.67741  2.69355  4.24690  2.90347  2.73739
3.18146  2.89801  2.37887  2.77519  2.98518  4.58477  3.61503
          0.02696  4.02262  4.74496  0.61958  0.77255  0.48576
0.95510
    181   3.11348  4.43724  4.75366  4.22299  3.43368  4.44249
4.89745  1.66694  4.10486  1.64011  3.24680  4.46643  4.71781
4.33858  4.29670  3.80195  3.36289  1.17939  5.39404  4.21883    198
- -
          2.68618  4.42225  2.77519  2.73123  3.46354  2.40513
3.72494  3.29354  2.67741  2.69355  4.24690  2.90347  2.73739
3.18146  2.89801  2.37887  2.77519  2.98518  4.58477  3.61503
          0.02696  4.02262  4.74496  0.61958  0.77255  0.48576
0.95510
    182   3.17148  4.48712  4.82190  4.27904  3.30774  4.49359
4.90156  1.57991  4.15852  1.28434  3.10664  4.52092  4.73273
4.34068  4.31994  3.84906  3.41120  1.57930  5.31817  4.19031    199
- -
          2.68618  4.42225  2.77519  2.73123  3.46354  2.40513
3.72494  3.29354  2.67741  2.69355  4.24690  2.90347  2.73739
3.18146  2.89801  2.37887  2.77519  2.98518  4.58477  3.61503
          0.02696  4.02262  4.74496  0.61958  0.77255  0.48576
0.95510
    183   3.22147  4.54141  4.80627  4.29083  3.28022  4.51184
4.90766  1.22924  4.12279  1.39926  3.11120  4.53824  4.75969
4.34732  4.28501  3.89111  3.46660  1.87730  5.30940  4.13657    200
- -
          2.68618  4.42225  2.77519  2.73123  3.46354  2.40513
3.72494  3.29354  2.67741  2.69355  4.24690  2.90347  2.73739
3.18146  2.89801  2.37887  2.77519  2.98518  4.58477  3.61503
          0.02696  4.02262  4.74496  0.61958  0.77255  0.48576
0.95510
```

184 2.94977 5.27365 2.38943 1.08504 4.55504 3.31409
3.80363 4.04597 2.65401 3.59069 4.49990 2.84570 3.90580
2.85087 3.09329 2.89103 3.23463 3.69156 5.75148 4.35362 201
- -
2.68618 4.42225 2.77519 2.73123 3.46354 2.40513
3.72494 3.29354 2.67741 2.69355 4.24690 2.90347 2.73739
3.18146 2.89801 2.37887 2.77519 2.98518 4.58477 3.61503
0.02696 4.02262 4.74496 0.61958 0.77255 0.48576
0.95510
185 3.15706 5.30096 0.72443 2.44038 4.69960 3.32995
4.07187 4.32326 3.18574 3.90577 4.88941 3.00343 3.99358
3.31287 3.72060 3.10404 3.49962 3.94721 5.83844 4.55063 202
- -
2.68618 4.42225 2.77519 2.73123 3.46354 2.40513
3.72494 3.29354 2.67741 2.69355 4.24690 2.90347 2.73739
3.18146 2.89801 2.37887 2.77519 2.98518 4.58477 3.61503
0.02696 4.02262 4.74496 0.61958 0.77255 0.48576
0.95510
186 2.46810 4.28774 3.64662 3.25869 3.83532 3.27031
4.20850 2.79949 3.13315 2.65992 3.79883 3.47895 3.91836
3.51550 3.41506 2.69502 1.29071 2.41936 5.37651 4.12085 203
- -
2.68618 4.42225 2.77519 2.73123 3.46354 2.40513
3.72494 3.29354 2.67741 2.69355 4.24690 2.90347 2.73739
3.18146 2.89801 2.37887 2.77519 2.98518 4.58477 3.61503
0.02696 4.02262 4.74496 0.61958 0.77255 0.48576
0.95510
187 2.65080 4.53562 3.33486 2.75393 3.71186 3.53422
3.75099 2.72485 2.50951 2.59268 3.61879 3.19454 3.63194
2.97918 1.90383 2.79350 2.88601 2.50113 5.10074 3.84391 204
- -
2.68618 4.42225 2.77519 2.73123 3.46354 2.40513
3.72494 3.29354 2.67741 2.69355 4.24690 2.90347 2.73739
3.18146 2.89801 2.37887 2.77519 2.98518 4.58477 3.61503
0.02696 4.02262 4.74496 0.61958 0.77255 0.48576
0.95510
188 3.08846 5.22327 2.56059 0.82931 4.58940 3.37593
3.97883 4.07348 2.87880 3.66752 4.63740 3.02771 3.98839
3.19292 3.30140 3.05703 3.39464 3.74275 5.73268 4.45792 205
- -
2.68618 4.42225 2.77519 2.73123 3.46354 2.40513
3.72494 3.29354 2.67741 2.69355 4.24690 2.90347 2.73739
3.18146 2.89801 2.37887 2.77519 2.98518 4.58477 3.61503
0.02696 4.02262 4.74496 0.61958 0.77255 0.48576
0.95510
189 3.32709 4.74617 4.42998 4.07280 3.19879 4.17148
4.68160 2.41344 3.83907 0.71022 3.17896 4.33366 4.58250
4.17133 3.98965 3.76330 3.61534 2.48932 5.13017 3.88739 206
- -
2.68618 4.42225 2.77519 2.73123 3.46354 2.40513
3.72494 3.29354 2.67741 2.69355 4.24690 2.90347 2.73739
3.18146 2.89801 2.37887 2.77519 2.98518 4.58477 3.61503
0.02696 4.02262 4.74496 0.61958 0.77255 0.48576
0.95510

```
    190   2.90640  5.07149  2.92382  2.56368  4.38380  3.48385
3.68733  3.84236  2.10996  3.35218  4.23505  3.04127  3.95203
1.40617  2.42175  2.90579  3.13750  3.51812  5.47576  4.18966    207
- -
          2.68618  4.42225  2.77519  2.73123  3.46354  2.40513
3.72494  3.29354  2.67741  2.69355  4.24690  2.90347  2.73739
3.18146  2.89801  2.37887  2.77519  2.98518  4.58477  3.61503
          0.02696  4.02262  4.74496  0.61958  0.77255  0.48576
0.95510
    191   2.34185  1.11971  4.07533  3.78040  3.89092  3.09438
4.45797  3.08418  3.59041  3.02023  4.01893  3.67404  3.84704
3.91796  3.74675  2.44498  2.83296  2.74810  5.38541  4.14433    208
- -
          2.68618  4.42225  2.77519  2.73123  3.46354  2.40513
3.72494  3.29354  2.67741  2.69355  4.24690  2.90347  2.73739
3.18146  2.89801  2.37887  2.77519  2.98518  4.58477  3.61503
          0.02696  4.02262  4.74496  0.61958  0.77255  0.48576
0.95510
    192   1.77936  4.55796  3.02522  2.58512  3.89197  3.35655
3.72107  3.27927  2.53765  2.88911  3.77832  3.05000  3.52390
2.87115  2.82956  2.59977  2.56306  2.93282  5.21875  3.93027    209
- -
          2.68618  4.42225  2.77519  2.73123  3.46354  2.40513
3.72494  3.29354  2.67741  2.69355  4.24690  2.90347  2.73739
3.18146  2.89801  2.37887  2.77519  2.98518  4.58477  3.61503
          0.02696  4.02262  4.74496  0.61958  0.77255  0.48576
0.95510
    193   1.01885  4.20538  3.37500  3.14355  4.22660  2.99160
4.22856  3.54914  3.20134  3.34196  4.21278  3.14217  3.73771
3.52004  3.49296  2.35186  2.76414  3.07653  5.60671  4.36644    210
- -
          2.68618  4.42225  2.77519  2.73123  3.46354  2.40513
3.72494  3.29354  2.67741  2.69355  4.24690  2.90347  2.73739
3.18146  2.89801  2.37887  2.77519  2.98518  4.58477  3.61503
          0.02696  4.02262  4.74496  0.61958  0.77255  0.48576
0.95510
    194   2.54307  4.76413  3.00064  2.44990  4.08069  3.39741
3.64954  3.48589  2.34277  3.08650  3.91218  2.98397  2.39784
2.78848  2.37473  2.47113  2.86184  3.05078  5.31783  4.00935    211
- -
          2.68618  4.42225  2.77519  2.73123  3.46354  2.40513
3.72494  3.29354  2.67741  2.69355  4.24690  2.90347  2.73739
3.18146  2.89801  2.37887  2.77519  2.98518  4.58477  3.61503
          0.02696  4.02262  4.74496  0.61958  0.77255  0.48576
0.95510
    195   2.90488  5.37085  1.62114  2.16168  4.69446  3.23262
3.79205  4.25025  2.79472  3.77423  4.62671  1.58783  3.85355
2.96521  3.36478  2.81858  3.20711  3.82337  5.90248  4.41770    212
- -
          2.68618  4.42225  2.77519  2.73123  3.46354  2.40513
3.72494  3.29354  2.67741  2.69355  4.24690  2.90347  2.73739
3.18146  2.89801  2.37887  2.77519  2.98518  4.58477  3.61503
          0.02696  4.02262  4.74496  0.61958  0.77255  0.48576
0.95510
```

```
    196   2.56503  3.75574  3.58294  3.01114  3.25818  3.57262
3.05553  2.65392  2.92400  1.89247  3.31204  3.37329  3.95292
3.22784  3.10340  2.83311  2.64726  2.36783  4.79717  3.56942    213
- -
          2.68618  4.42225  2.77519  2.73123  3.46354  2.40513
3.72494  3.29354  2.67741  2.69355  4.24690  2.90347  2.73739
3.18146  2.89801  2.37887  2.77519  2.98518  4.58477  3.61503
          0.02696  4.02262  4.74496  0.61958  0.77255  0.48576
0.95510
    197   3.08660  4.42397  4.71842  4.21259  3.54101  4.39957
4.93629  1.62073  4.08602  1.99767  3.36422  4.45033  4.71870
4.36528  4.29732  3.77727  3.35213  1.01484  5.48752  4.27138    214
- -
          2.68618  4.42225  2.77519  2.73123  3.46354  2.40513
3.72494  3.29354  2.67741  2.69355  4.24690  2.90347  2.73739
3.18146  2.89801  2.37887  2.77519  2.98518  4.58477  3.61503
          0.02696  4.02262  4.74496  0.61958  0.77255  0.48576
0.95510
    198   1.58225  4.37678  3.24358  2.78700  3.51872  3.26248
3.86533  3.20410  2.78129  2.89874  3.77336  3.17174  2.83628
3.02005  3.16247  2.54997  2.78838  2.88747  5.21603  3.94565    215
- -
          2.68618  4.42225  2.77519  2.73123  3.46354  2.40513
3.72494  3.29354  2.67741  2.69355  4.24690  2.90347  2.73739
3.18146  2.89801  2.37887  2.77519  2.98518  4.58477  3.61503
          0.02696  4.02262  4.74496  0.61958  0.77255  0.48576
0.95510
    199   3.30228  4.67642  4.71043  4.16693  3.06443  4.47559
4.76357  2.21771  3.96557  0.89985  2.24821  4.45424  4.69143
4.16177  4.11777  3.83535  3.53106  2.34864  5.13601  4.02609    216
- -
          2.68618  4.42225  2.77519  2.73123  3.46354  2.40513
3.72494  3.29354  2.67741  2.69355  4.24690  2.90347  2.73739
3.18146  2.89801  2.37887  2.77519  2.98518  4.58477  3.61503
          0.02696  4.02262  4.74496  0.61958  0.77255  0.48576
0.95510
    200   2.81525  4.00430  3.60822  3.01444  4.14371  3.49715
3.84816  3.54077  2.31751  3.15680  4.10292  3.35846  3.99635
3.05427  1.09884  2.92901  3.10726  3.24776  5.33523  4.12283    217
- -
          2.68618  4.42225  2.77519  2.73123  3.46354  2.40513
3.72494  3.29354  2.67741  2.69355  4.24690  2.90347  2.73739
3.18146  2.89801  2.37887  2.77519  2.98518  4.58477  3.61503
          0.02696  4.02262  4.74496  0.61958  0.77255  0.48576
0.95510
    201   2.23321  4.24097  3.51628  3.24665  4.11727  3.09040
4.27092  3.19028  3.20683  3.09095  4.07774  3.41430  3.81487
3.56962  3.47753  2.56210  1.12788  2.84215  5.56418  4.34043    218
- -
          2.68618  4.42225  2.77519  2.73123  3.46354  2.40513
3.72494  3.29354  2.67741  2.69355  4.24690  2.90347  2.73739
3.18146  2.89801  2.37887  2.77519  2.98518  4.58477  3.61503
          0.02696  4.02262  4.74496  0.61958  0.77255  0.48576
0.95510
```

```
    202   2.77447  5.08145  3.29265  2.65033  4.45951  3.59253
3.33919  3.83439  1.58982  3.20960  4.15635  3.10065  3.95752
2.69095  1.70229  2.89078  3.03552  3.50365  5.41981  4.19058    219
- -
          2.68618  4.42225  2.77519  2.73123  3.46354  2.40513
3.72494  3.29354  2.67741  2.69355  4.24690  2.90347  2.73739
3.18146  2.89801  2.37887  2.77519  2.98518  4.58477  3.61503
          0.02696  4.02262  4.74496  0.61958  0.77255  0.48576
0.95510
    203   2.31987  5.13210  1.91899  2.10237  4.43112  3.31865
3.63279  3.89784  2.37912  3.42130  4.20959  2.79646  3.32170
2.68959  2.90087  2.65593  2.94495  3.50209  5.59579  4.18550    220
- -
          2.68618  4.42225  2.77519  2.73123  3.46354  2.40513
3.72494  3.29354  2.67741  2.69355  4.24690  2.90347  2.73739
3.18146  2.89801  2.37887  2.77519  2.98518  4.58477  3.61503
          0.02696  4.02262  4.74496  0.61958  0.77255  0.48576
0.95510
    204   2.70093  4.66990  2.58860  2.77616  4.51323  0.92064
4.17171  4.04322  3.19730  3.68704  4.57914  3.19982  3.88755
3.43640  3.59320  2.80170  3.13307  3.57047  5.71158  4.50887    221
- -
          2.68618  4.42225  2.77519  2.73123  3.46354  2.40513
3.72494  3.29354  2.67741  2.69355  4.24690  2.90347  2.73739
3.18146  2.89801  2.37887  2.77519  2.98518  4.58477  3.61503
          0.02696  4.02262  4.74496  0.61958  0.77255  0.48576
0.95510
    205   2.54167  4.26839  3.86348  3.33926  3.51445  3.72197
4.16223  2.16236  3.02275  2.37125  3.44167  3.65099  4.15040
3.54609  3.47754  2.96951  2.96091  1.34513  5.08802  3.86493    222
- -
          2.68618  4.42225  2.77519  2.73123  3.46354  2.40513
3.72494  3.29354  2.67741  2.69355  4.24690  2.90347  2.73739
3.18146  2.89801  2.37887  2.77519  2.98518  4.58477  3.61503
          0.03705  4.02262  3.99187  0.61958  0.77255  0.48576
0.95510
    206   1.78024  4.26459  3.99795  3.52134  3.65093  3.60830
4.37039  2.13719  3.43151  2.43172  3.54334  3.75049  4.14835
3.73909  3.70675  2.98869  2.97971  1.48753  5.30126  4.08114    223
- -
          2.68618  4.42225  2.77519  2.73123  3.46354  2.40513
3.72494  3.29354  2.67741  2.69355  4.24690  2.90347  2.73739
3.18146  2.89801  2.37887  2.77519  2.98518  4.58477  3.61503
          0.02723  4.01280  4.73514  0.61958  0.77255  0.47937
0.96541
    207   2.31365  4.28024  3.26925  2.95180  4.22136  3.03653
4.07691  3.62077  2.99097  3.31416  4.15142  3.05500  3.73332
3.32091  3.33518  1.39212  1.99173  3.14458  5.55564  4.30420    224
- -
          2.68618  4.42225  2.77519  2.73123  3.46354  2.40513
3.72494  3.29354  2.67741  2.69355  4.24690  2.90347  2.73739
3.18146  2.89801  2.37887  2.77519  2.98518  4.58477  3.61503
          0.02696  4.02262  4.74496  0.61958  0.77255  0.48576
0.95510
```

208 3.28217 4.65965 4.69372 4.14678 3.06488 4.45454
4.74266 2.22272 3.95124 0.98810 1.99286 4.43215 4.67406
4.14506 4.10443 3.80962 3.51033 2.35036 5.12527 4.01757 225
- -
2.68618 4.42225 2.77519 2.73123 3.46354 2.40513
3.72494 3.29354 2.67741 2.69355 4.24690 2.90347 2.73739
3.18146 2.89801 2.37887 2.77519 2.98518 4.58477 3.61503
0.02696 4.02262 4.74496 0.61958 0.77255 0.48576
0.95510
209 2.24312 4.56654 3.01168 2.58140 4.08806 3.25177
3.77405 3.49676 2.60603 3.12693 3.95507 2.88556 3.78365
2.65136 3.02471 1.89373 2.30799 3.12768 5.37828 4.07815 226
- -
2.68618 4.42225 2.77519 2.73123 3.46354 2.40513
3.72494 3.29354 2.67741 2.69355 4.24690 2.90347 2.73739
3.18146 2.89801 2.37887 2.77519 2.98518 4.58477 3.61503
0.02696 4.02262 4.74496 0.61958 0.77255 0.48576
0.95510
210 2.94549 5.54281 1.39623 1.64036 4.81802 3.23740
3.75870 4.34229 2.75829 3.82705 4.66776 2.65606 3.84973
2.74303 3.34939 2.82647 3.22791 3.90970 5.98426 4.47283 227
- -
2.68618 4.42225 2.77519 2.73123 3.46354 2.40513
3.72494 3.29354 2.67741 2.69355 4.24690 2.90347 2.73739
3.18146 2.89801 2.37887 2.77519 2.98518 4.58477 3.61503
0.02696 4.02262 4.74496 0.61958 0.77255 0.48576
0.95510
211 3.32709 4.74617 4.42998 4.07280 3.19879 4.17148
4.68160 2.41344 3.83907 0.71022 3.17896 4.33366 4.58250
4.17133 3.98965 3.76330 3.61534 2.48932 5.13017 3.88739 228
- -
2.68618 4.42225 2.77519 2.73123 3.46354 2.40513
3.72494 3.29354 2.67741 2.69355 4.24690 2.90347 2.73739
3.18146 2.89801 2.37887 2.77519 2.98518 4.58477 3.61503
0.02696 4.02262 4.74496 0.61958 0.77255 0.48576
0.95510
212 2.92784 4.43147 4.33116 4.00117 3.66379 3.87690
4.77821 2.05427 3.87125 2.32619 3.60830 4.16627 4.43068
4.22125 4.06895 3.39165 3.28677 0.85478 5.47286 4.21283 229
- -
2.68618 4.42225 2.77519 2.73123 3.46354 2.40513
3.72494 3.29354 2.67741 2.69355 4.24690 2.90347 2.73739
3.18146 2.89801 2.37887 2.77519 2.98518 4.58477 3.61503
0.02696 4.02262 4.74496 0.61958 0.77255 0.48576
0.95510
213 3.07753 5.08484 3.51154 2.90282 4.50949 3.65942
3.69172 3.90063 1.88010 3.38139 4.29045 3.27943 4.07056
2.84846 1.07980 3.10473 3.27109 3.60070 5.43168 4.26586 230
- -
2.68618 4.42225 2.77519 2.73123 3.46354 2.40513
3.72494 3.29354 2.67741 2.69355 4.24690 2.90347 2.73739
3.18146 2.89801 2.37887 2.77519 2.98518 4.58477 3.61503
0.02696 4.02262 4.74496 0.61958 0.77255 0.48576
0.95510

214 1.86457 4.19229 3.40251 3.15231 4.30801 2.95953
4.24251 3.67584 3.21684 3.41670 4.27050 3.31239 3.71389
3.51961 3.51977 1.14114 2.74148 3.15250 5.66669 4.43630 231
- -
2.68618 4.42225 2.77519 2.73123 3.46354 2.40513
3.72494 3.29354 2.67741 2.69355 4.24690 2.90347 2.73739
3.18146 2.89801 2.37887 2.77519 2.98518 4.58477 3.61503
0.02696 4.02262 4.74496 0.61958 0.77255 0.48576
0.95510
215 1.98750 4.19746 3.43721 3.07663 4.22190 2.88339
4.14075 3.60022 3.09572 3.30264 4.13178 3.27885 3.70837
3.39711 3.43124 1.50608 1.92179 3.10542 5.56597 4.34478 232
- -
2.68618 4.42225 2.77519 2.73123 3.46354 2.40513
3.72494 3.29354 2.67741 2.69355 4.24690 2.90347 2.73739
3.18146 2.89801 2.37887 2.77519 2.98518 4.58477 3.61503
0.02696 4.02262 4.74496 0.61958 0.77255 0.48576
0.95510
216 3.29838 4.67256 4.70927 4.16444 3.06387 4.47257
4.76025 2.21884 3.96534 0.91790 2.18554 4.45120 4.68822
4.15930 4.11722 3.83089 3.52672 2.34982 5.13361 4.02563 233
- -
2.68618 4.42225 2.77519 2.73123 3.46354 2.40513
3.72494 3.29354 2.67741 2.69355 4.24690 2.90347 2.73739
3.18146 2.89801 2.37887 2.77519 2.98518 4.58477 3.61503
0.02696 4.02262 4.74496 0.61958 0.77255 0.48576
0.95510
217 3.18120 5.00542 3.68780 3.17038 4.39834 3.63746
3.94268 3.95237 2.31116 3.44111 4.43934 3.51762 4.13451
3.15727 0.77162 3.25933 3.43807 3.67032 5.42160 4.29519 234
- -
2.68618 4.42225 2.77519 2.73123 3.46354 2.40513
3.72494 3.29354 2.67741 2.69355 4.24690 2.90347 2.73739
3.18146 2.89801 2.37887 2.77519 2.98518 4.58477 3.61503
0.02696 4.02262 4.74496 0.61958 0.77255 0.48576
0.95510
218 3.13684 4.63695 4.19709 3.75073 3.01359 4.12287
4.40098 2.40230 3.51639 0.90526 3.07119 4.05056 4.47478
3.62184 3.71824 3.51510 3.39411 2.47671 4.96341 3.66446 235
- -
2.68618 4.42225 2.77519 2.73123 3.46354 2.40513
3.72494 3.29354 2.67741 2.69355 4.24690 2.90347 2.73739
3.18146 2.89801 2.37887 2.77519 2.98518 4.58477 3.61503
0.02696 4.02262 4.74496 0.61958 0.77255 0.48576
0.95510
219 3.18120 5.00542 3.68780 3.17038 4.39834 3.63746
3.94268 3.95237 2.31116 3.44111 4.43934 3.51762 4.13451
3.15727 0.77162 3.25933 3.43807 3.67032 5.42160 4.29519 236
- -
2.68618 4.42225 2.77519 2.73123 3.46354 2.40513
3.72494 3.29354 2.67741 2.69355 4.24690 2.90347 2.73739
3.18146 2.89801 2.37887 2.77519 2.98518 4.58477 3.61503
0.02696 4.02262 4.74496 0.61958 0.77255 0.48576
0.95510

220 2.96212 4.69205 3.58591 3.44827 4.42684 3.37117
4.51157 4.01315 3.51887 3.62279 4.68053 3.73880 0.59042
3.90306 3.74552 3.13480 3.41594 3.64102 5.57002 4.55027 237
- -
2.68618 4.42225 2.77519 2.73123 3.46354 2.40513
3.72494 3.29354 2.67741 2.69355 4.24690 2.90347 2.73739
3.18146 2.89801 2.37887 2.77519 2.98518 4.58477 3.61503
0.02696 4.02262 4.74496 0.61958 0.77255 0.48576
0.95510
221 2.90927 5.17012 1.01308 2.30580 4.48242 3.25612
3.90184 3.96044 2.93996 3.61936 4.57050 2.85980 3.89913
3.10959 3.47715 2.87475 3.24416 3.40056 5.78335 4.34427 238
- -
2.68618 4.42225 2.77519 2.73123 3.46354 2.40513
3.72494 3.29354 2.67741 2.69355 4.24690 2.90347 2.73739
3.18146 2.89801 2.37887 2.77519 2.98518 4.58477 3.61503
0.02696 4.02262 4.74496 0.61958 0.77255 0.48576
0.95510
222 3.18120 5.00542 3.68780 3.17038 4.39834 3.63746
3.94268 3.95237 2.31116 3.44111 4.43934 3.51762 4.13451
3.15727 0.77162 3.25933 3.43807 3.67032 5.42160 4.29519 239
- -
2.68618 4.42225 2.77519 2.73123 3.46354 2.40513
3.72494 3.29354 2.67741 2.69355 4.24690 2.90347 2.73739
3.18146 2.89801 2.37887 2.77519 2.98518 4.58477 3.61503
0.02696 4.02262 4.74496 0.61958 0.77255 0.48576
0.95510
223 3.11435 4.44022 4.79051 4.30435 3.53699 4.43291
5.01716 1.08905 4.17939 2.02370 3.34807 4.52365 4.75500
4.44729 4.37454 3.82868 3.38503 1.43335 5.52166 4.31775 240
- -
2.68618 4.42225 2.77519 2.73123 3.46354 2.40513
3.72494 3.29354 2.67741 2.69355 4.24690 2.90347 2.73739
3.18146 2.89801 2.37887 2.77519 2.98518 4.58477 3.61503
0.02696 4.02262 4.74496 0.61958 0.77255 0.48576
0.95510
224 2.67322 4.31233 3.77727 3.25786 3.49483 3.66191
4.13428 1.97259 3.17004 2.32524 3.41612 3.58927 2.25431
3.48813 3.47532 2.98427 2.95695 1.90455 5.09791 3.87111 241
- -
2.68618 4.42225 2.77519 2.73123 3.46354 2.40513
3.72494 3.29354 2.67741 2.69355 4.24690 2.90347 2.73739
3.18146 2.89801 2.37887 2.77519 2.98518 4.58477 3.61503
0.02696 4.02262 4.74496 0.61958 0.77255 0.48576
0.95510
225 3.10538 4.41955 4.81540 4.32210 3.57684 4.45314
5.03708 1.27313 4.20958 2.07696 3.38171 4.53899 4.76720
4.47453 4.40721 3.84367 3.37280 1.18355 5.54803 4.34251 242
- -
2.68618 4.42225 2.77519 2.73123 3.46354 2.40513
3.72494 3.29354 2.67741 2.69355 4.24690 2.90347 2.73739
3.18146 2.89801 2.37887 2.77519 2.98518 4.58477 3.61503
0.02696 4.02262 4.74496 0.61958 0.77255 0.48576
0.95510

226 2.86181 4.62005 3.56615 3.48578 4.62096 0.53093
4.59479 4.25913 3.68734 3.89379 4.87234 3.73076 4.01589
3.99980 3.90487 3.03839 3.35475 3.75848 5.67431 4.72868 243
- -
2.68618 4.42225 2.77519 2.73123 3.46354 2.40513
3.72494 3.29354 2.67741 2.69355 4.24690 2.90347 2.73739
3.18146 2.89801 2.37887 2.77519 2.98518 4.58477 3.61503
0.02696 4.02262 4.74496 0.61958 0.77255 0.48576
0.95510
227 3.08846 5.22327 2.56059 0.82931 4.58940 3.37593
3.97883 4.07348 2.87880 3.66752 4.63740 3.02771 3.98839
3.19292 3.30140 3.05703 3.39464 3.74275 5.73268 4.45792 244
- -
2.68618 4.42225 2.77519 2.73123 3.46354 2.40513
3.72494 3.29354 2.67741 2.69355 4.24690 2.90347 2.73739
3.18146 2.89801 2.37887 2.77519 2.98518 4.58477 3.61503
0.02696 4.02262 4.74496 0.61958 0.77255 0.48576
0.95510
228 2.92784 4.43147 4.33116 4.00117 3.66379 3.87690
4.77821 2.05427 3.87125 2.32619 3.60830 4.16627 4.43068
4.22125 4.06895 3.39165 3.28677 0.85478 5.47286 4.21283 245
- -
2.68618 4.42225 2.77519 2.73123 3.46354 2.40513
3.72494 3.29354 2.67741 2.69355 4.24690 2.90347 2.73739
3.18146 2.89801 2.37887 2.77519 2.98518 4.58477 3.61503
0.02696 4.02262 4.74496 0.61958 0.77255 0.48576
0.95510
229 3.18120 5.00542 3.68780 3.17038 4.39834 3.63746
3.94268 3.95237 2.31116 3.44111 4.43934 3.51762 4.13451
3.15727 0.77162 3.25933 3.43807 3.67032 5.42160 4.29519 246
- -
2.68618 4.42225 2.77519 2.73123 3.46354 2.40513
3.72494 3.29354 2.67741 2.69355 4.24690 2.90347 2.73739
3.18146 2.89801 2.37887 2.77519 2.98518 4.58477 3.61503
0.02696 4.02262 4.74496 0.61958 0.77255 0.48576
0.95510
230 2.86181 4.62005 3.56615 3.48578 4.62096 0.53093
4.59479 4.25913 3.68734 3.89379 4.87234 3.73076 4.01589
3.99980 3.90487 3.03839 3.35475 3.75848 5.67431 4.72868 247
- -
2.68618 4.42225 2.77519 2.73123 3.46354 2.40513
3.72494 3.29354 2.67741 2.69355 4.24690 2.90347 2.73739
3.18146 2.89801 2.37887 2.77519 2.98518 4.58477 3.61503
0.02696 4.02262 4.74496 0.61958 0.77255 0.48576
0.95510
231 1.61188 4.38469 3.01311 2.74962 4.19754 2.40134
3.94054 3.61821 2.84977 3.26060 4.08058 3.11450 2.87438
3.14874 3.25966 2.26223 2.76552 3.16767 5.50712 4.23141 248
- -
2.68618 4.42225 2.77519 2.73123 3.46354 2.40513
3.72494 3.29354 2.67741 2.69355 4.24690 2.90347 2.73739
3.18146 2.89801 2.37887 2.77519 2.98518 4.58477 3.61503
0.02696 4.02262 4.74496 0.61958 0.77255 0.48576
0.95510

```
    232   3.08846  5.22327  2.56059  0.82931  4.58940  3.37593
3.97883  4.07348  2.87880  3.66752  4.63740  3.02771  3.98839
3.19292  3.30140  3.05703  3.39464  3.74275  5.73268  4.45792    249
- -
          2.68618  4.42225  2.77519  2.73123  3.46354  2.40513
3.72494  3.29354  2.67741  2.69355  4.24690  2.90347  2.73739
3.18146  2.89801  2.37887  2.77519  2.98518  4.58477  3.61503
          0.02696  4.02262  4.74496  0.61958  0.77255  0.48576
0.95510
    233   0.81728  4.27297  3.62041  3.42136  4.20706  3.09296
4.42492  3.35544  3.44162  3.22989  4.24718  3.53130  3.84663
3.76944  3.68019  2.62296  2.91086  2.98372  5.61916  4.43687    250
- -
          2.68618  4.42225  2.77519  2.73123  3.46354  2.40513
3.72494  3.29354  2.67741  2.69355  4.24690  2.90347  2.73739
3.18146  2.89801  2.37887  2.77519  2.98518  4.58477  3.61503
          0.02696  4.02262  4.74496  0.61958  0.77255  0.48576
0.95510
    234   3.32709  4.74617  4.42998  4.07280  3.19879  4.17148
4.68160  2.41344  3.83907  0.71022  3.17896  4.33366  4.58250
4.17133  3.98965  3.76330  3.61534  2.48932  5.13017  3.88739    251
- -
          2.68618  4.42225  2.77519  2.73123  3.46354  2.40513
3.72494  3.29354  2.67741  2.69355  4.24690  2.90347  2.73739
3.18146  2.89801  2.37887  2.77519  2.98518  4.58477  3.61503
          0.02696  4.02262  4.74496  0.61958  0.77255  0.48576
0.95510
    235   2.93419  5.38162  1.08643  2.10703  4.73536  3.21579
3.82738  4.24599  2.87591  3.80745  4.68602  2.72969  3.86347
3.01111  3.46240  2.67125  3.25179  3.82869  5.96703  4.47181    252
- -
          2.68618  4.42225  2.77519  2.73123  3.46354  2.40513
3.72494  3.29354  2.67741  2.69355  4.24690  2.90347  2.73739
3.18146  2.89801  2.37887  2.77519  2.98518  4.58477  3.61503
          0.02696  4.02262  4.74496  0.61958  0.77255  0.48576
0.95510
    236   3.30910  4.69575  4.67257  4.14994  3.07556  4.45295
4.75210  2.21647  3.91691  0.82780  2.63893  4.43612  4.69281
4.15499  4.07615  3.83076  3.54706  2.33403  5.14069  3.99014    253
- -
          2.68618  4.42225  2.77519  2.73123  3.46354  2.40513
3.72494  3.29354  2.67741  2.69355  4.24690  2.90347  2.73739
3.18146  2.89801  2.37887  2.77519  2.98518  4.58477  3.61503
          0.02696  4.02262  4.74496  0.61958  0.77255  0.48576
0.95510
    237   3.23861  4.57004  4.80978  4.26621  3.20344  4.51507
4.88144  1.72390  4.12245  1.03749  3.00432  4.52800  4.73763
4.29560  4.27813  3.87277  3.47498  1.89195  5.26020  4.14736    254
- -
          2.68618  4.42225  2.77519  2.73123  3.46354  2.40513
3.72494  3.29354  2.67741  2.69355  4.24690  2.90347  2.73739
3.18146  2.89801  2.37887  2.77519  2.98518  4.58477  3.61503
          0.02696  4.02262  4.74496  0.61958  0.77255  0.48576
0.95510
```

```
    238   3.11448  5.07907  3.30011  2.89647  4.49993  3.58511
3.85036  3.93922  0.84983  3.47273  4.42835  3.31347  4.07671
3.03496  2.47086  3.14796  3.35771  3.63935  5.49460  4.33003    255
- -
          2.68618  4.42225  2.77519  2.73123  3.46354  2.40513
3.72494  3.29354  2.67741  2.69355  4.24690  2.90347  2.73739
3.18146  2.89801  2.37887  2.77519  2.98518  4.58477  3.61503
          0.02696  4.02262  4.74496  0.61958  0.77255  0.48576
0.95510
    239   1.27292  4.21216  3.80005  3.41997  3.87285  3.24446
4.33277  2.67009  3.35319  2.74215  3.79996  3.56516  3.91955
3.66792  3.62422  2.67781  2.85438  1.96379  5.43373  4.21265    256
- -
          2.68618  4.42225  2.77519  2.73123  3.46354  2.40513
3.72494  3.29354  2.67741  2.69355  4.24690  2.90347  2.73739
3.18146  2.89801  2.37887  2.77519  2.98518  4.58477  3.61503
          0.02696  4.02262  4.74496  0.61958  0.77255  0.48576
0.95510
    240   3.59642  4.91567  4.32809  4.07288  2.83521  3.90453
4.09650  3.62874  3.79302  2.99671  4.25522  4.19217  4.43527
4.17286  3.89421  3.79859  3.90492  3.52317  0.63014  2.82983    257
- -
          2.68618  4.42225  2.77519  2.73123  3.46354  2.40513
3.72494  3.29354  2.67741  2.69355  4.24690  2.90347  2.73739
3.18146  2.89801  2.37887  2.77519  2.98518  4.58477  3.61503
          0.02696  4.02262  4.74496  0.61958  0.77255  0.48576
0.95510
    241   2.53851  4.57434  2.80341  2.67185  4.42164  1.29886
4.03707  3.95495  2.97545  3.58070  4.43329  2.26873  3.80554
3.26931  3.36661  2.63731  2.96755  3.45641  5.69216  4.38806    258
- -
          2.68618  4.42225  2.77519  2.73123  3.46354  2.40513
3.72494  3.29354  2.67741  2.69355  4.24690  2.90347  2.73739
3.18146  2.89801  2.37887  2.77519  2.98518  4.58477  3.61503
          0.02696  4.02262  4.74496  0.61958  0.77255  0.48576
0.95510
    242   2.49976  4.34431  3.56805  3.33000  4.10800  3.17788
4.33500  3.25370  3.27593  3.11180  4.15459  3.51326  3.89436
3.66375  3.52580  2.70296  0.94063  2.93233  5.53249  4.32538    259
- -
          2.68618  4.42225  2.77519  2.73123  3.46354  2.40513
3.72494  3.29354  2.67741  2.69355  4.24690  2.90347  2.73739
3.18146  2.89801  2.37887  2.77519  2.98518  4.58477  3.61503
          0.02696  4.02262  4.74496  0.61958  0.77255  0.48576
0.95510
    243   2.86181  4.62005  3.56615  3.48578  4.62096  0.53093
4.59479  4.25913  3.68734  3.89379  4.87234  3.73076  4.01589
3.99980  3.90487  3.03839  3.35475  3.75848  5.67431  4.72868    260
- -
          2.68618  4.42225  2.77519  2.73123  3.46354  2.40513
3.72494  3.29354  2.67741  2.69355  4.24690  2.90347  2.73739
3.18146  2.89801  2.37887  2.77519  2.98518  4.58477  3.61503
          0.02696  4.02262  4.74496  0.61958  0.77255  0.48576
0.95510
```

244 3.17169 4.95855 3.28396 3.06125 3.39905 3.58234
0.86156 3.89784 2.90062 3.37193 4.42840 3.46359 4.14428
3.43008 3.15617 3.24168 3.48110 3.63022 4.86163 3.34615 261
- -
2.68618 4.42225 2.77519 2.73123 3.46354 2.40513
3.72494 3.29354 2.67741 2.69355 4.24690 2.90347 2.73739
3.18146 2.89801 2.37887 2.77519 2.98518 4.58477 3.61503
0.02696 4.02262 4.74496 0.61958 0.77255 0.48576
0.95510
245 2.96212 4.69205 3.58591 3.44827 4.42684 3.37117
4.51157 4.01315 3.51887 3.62279 4.68053 3.73880 0.59042
3.90306 3.74552 3.13480 3.41594 3.64102 5.57002 4.55027 262
- -
2.68618 4.42225 2.77519 2.73123 3.46354 2.40513
3.72494 3.29354 2.67741 2.69355 4.24690 2.90347 2.73739
3.18146 2.89801 2.37887 2.77519 2.98518 4.58477 3.61503
0.02696 4.02262 4.74496 0.61958 0.77255 0.48576
0.95510
246 2.86181 4.62005 3.56615 3.48578 4.62096 0.53093
4.59479 4.25913 3.68734 3.89379 4.87234 3.73076 4.01589
3.99980 3.90487 3.03839 3.35475 3.75848 5.67431 4.72868 263
- -
2.68618 4.42225 2.77519 2.73123 3.46354 2.40513
3.72494 3.29354 2.67741 2.69355 4.24690 2.90347 2.73739
3.18146 2.89801 2.37887 2.77519 2.98518 4.58477 3.61503
0.02696 4.02262 4.74496 0.61958 0.77255 0.48576
0.95510
247 2.55426 4.46611 3.38098 3.28393 4.49890 0.71724
4.44424 4.03451 3.48427 3.71982 4.65753 3.51545 3.89781
3.79214 3.74191 2.77855 3.10585 3.52013 5.69336 4.60055 264
- -
2.68618 4.42225 2.77519 2.73123 3.46354 2.40513
3.72494 3.29354 2.67741 2.69355 4.24690 2.90347 2.73739
3.18146 2.89801 2.37887 2.77519 2.98518 4.58477 3.61503
0.02696 4.02262 4.74496 0.61958 0.77255 0.48576
0.95510
248 2.94991 3.81794 4.70253 4.16730 3.52028 4.30317
4.80608 1.21509 4.05805 2.04759 3.35614 4.36838 4.62520
4.29813 4.24443 3.65802 3.24824 1.44072 5.36357 4.16959 265
- -
2.68618 4.42225 2.77519 2.73123 3.46354 2.40513
3.72494 3.29354 2.67741 2.69355 4.24690 2.90347 2.73739
3.18146 2.89801 2.37887 2.77519 2.98518 4.58477 3.61503
0.02696 4.02262 4.74496 0.61958 0.77255 0.48576
0.95510
249 1.55533 4.16934 3.42406 3.13381 4.32015 1.65557
4.21955 3.71431 3.20949 3.41272 4.23385 3.29556 3.69241
3.48651 3.53116 2.23236 2.52963 3.16646 5.66041 4.45089 266
- -
2.68618 4.42225 2.77519 2.73123 3.46354 2.40513
3.72494 3.29354 2.67741 2.69355 4.24690 2.90347 2.73739
3.18146 2.89801 2.37887 2.77519 2.98518 4.58477 3.61503
0.02696 4.02262 4.74496 0.61958 0.77255 0.48576
0.95510

```
    250   2.49976  4.34431  3.56805  3.33000  4.10800  3.17788
4.33500  3.25370  3.27593  3.11180  4.15459  3.51326  3.89436
3.66375  3.52580  2.70296  0.94063  2.93233  5.53249  4.32538    267
- -
          2.68618  4.42225  2.77519  2.73123  3.46354  2.40513
3.72494  3.29354  2.67741  2.69355  4.24690  2.90347  2.73739
3.18146  2.89801  2.37887  2.77519  2.98518  4.58477  3.61503
          0.02696  4.02262  4.74496  0.61958  0.77255  0.48576
0.95510
    251   3.17827  4.48512  4.84624  4.31432  3.36300  4.52119
4.95686  1.24299  4.18757  1.51951  3.16393  4.55468  4.76618
4.39242  4.35667  3.88648  3.42283  1.65026  5.37964  4.23085    268
- -
          2.68618  4.42225  2.77519  2.73123  3.46354  2.40513
3.72494  3.29354  2.67741  2.69355  4.24690  2.90347  2.73739
3.18146  2.89801  2.37887  2.77519  2.98518  4.58477  3.61503
          0.02696  4.02262  4.74496  0.61958  0.77255  0.48576
0.95510
    252   3.17169  4.95855  3.28396  3.06125  3.39905  3.58234
0.86156  3.89784  2.90062  3.37193  4.42840  3.46359  4.14428
3.43008  3.15617  3.24168  3.48110  3.63022  4.86163  3.34615    269
- -
          2.68618  4.42225  2.77519  2.73123  3.46354  2.40513
3.72494  3.29354  2.67741  2.69355  4.24690  2.90347  2.73739
3.18146  2.89801  2.37887  2.77519  2.98518  4.58477  3.61503
          0.02696  4.02262  4.74496  0.61958  0.77255  0.48576
0.95510
    253   0.90846  4.20649  3.54679  3.33772  4.19249  3.02519
4.35858  3.36193  3.36057  3.24581  4.21363  3.44274  3.78628
3.68417  3.61296  2.48133  2.81643  2.95740  5.62499  4.39847    270
- -
          2.68618  4.42225  2.77519  2.73123  3.46354  2.40513
3.72494  3.29354  2.67741  2.69355  4.24690  2.90347  2.73739
3.18146  2.89801  2.37887  2.77519  2.98518  4.58477  3.61503
          0.02696  4.02262  4.74496  0.61958  0.77255  0.48576
0.95510
    254   2.49276  4.55741  2.76055  2.61176  4.41589  1.46173
3.98507  3.91788  2.91355  3.53873  4.37562  2.22433  3.77668
3.20166  3.32100  2.51281  2.91525  3.41693  5.70041  4.37227    271
- -
          2.68618  4.42225  2.77519  2.73123  3.46354  2.40513
3.72494  3.29354  2.67741  2.69355  4.24690  2.90347  2.73739
3.18146  2.89801  2.37887  2.77519  2.98518  4.58477  3.61503
          0.02696  4.02262  4.74496  0.61958  0.77255  0.48576
0.95510
    255   2.43131  4.43648  3.09706  2.72576  4.09315  3.00886
3.51998  3.52400  2.75707  3.17171  4.01011  3.11235  3.77355
3.10695  3.14098  1.52685  2.26598  3.12398  5.41396  4.13016    272
- -
          2.68618  4.42225  2.77519  2.73123  3.46354  2.40513
3.72494  3.29354  2.67741  2.69355  4.24690  2.90347  2.73739
3.18146  2.89801  2.37887  2.77519  2.98518  4.58477  3.61503
          0.02696  4.02262  4.74496  0.61958  0.77255  0.48576
0.95510
```

```
    256   1.27835  4.25954  3.34267  2.90833  4.05297  2.72788
4.04720  3.38004  2.99184  3.11561  3.97508  3.24842  3.45571
3.30801  3.33847  2.50483  2.75369  2.77344  5.42362  4.18799    273
- -
          2.68618  4.42225  2.77519  2.73123  3.46354  2.40513
3.72494  3.29354  2.67741  2.69355  4.24690  2.90347  2.73739
3.18146  2.89801  2.37887  2.77519  2.98518  4.58477  3.61503
          0.02696  4.02262  4.74496  0.61958  0.77255  0.48576
0.95510
    257   2.54050  4.21279  3.67115  3.09273  3.30998  3.62297
3.88063  2.23728  2.83261  1.81203  3.24626  3.43788  3.99431
3.23542  3.02030  2.89101  2.78704  2.41734  4.80605  3.48379    274
- -
          2.68618  4.42225  2.77519  2.73123  3.46354  2.40513
3.72494  3.29354  2.67741  2.69355  4.24690  2.90347  2.73739
3.18146  2.89801  2.37887  2.77519  2.98518  4.58477  3.61503
          0.02696  4.02262  4.74496  0.61958  0.77255  0.48576
0.95510
    258   2.05323  4.20525  3.27371  3.08142  4.41507  1.26462
4.24790  3.84044  3.24678  3.53078  4.35425  3.26749  3.69967
3.51761  3.56462  2.08698  2.74521  3.25478  5.74525  4.52295    275
- -
          2.68618  4.42225  2.77519  2.73123  3.46354  2.40513
3.72494  3.29354  2.67741  2.69355  4.24690  2.90347  2.73739
3.18146  2.89801  2.37887  2.77519  2.98518  4.58477  3.61503
          0.02696  4.02262  4.74496  0.61958  0.77255  0.48576
0.95510
    259   1.01480  4.15533  3.49931  3.24868  4.27617  2.65025
4.29284  3.54694  3.29492  3.34907  4.21698  3.36313  3.72021
3.58953  3.57558  2.43085  2.59878  3.05914  5.65267  4.45392    276
- -
          2.68618  4.42225  2.77519  2.73123  3.46354  2.40513
3.72494  3.29354  2.67741  2.69355  4.24690  2.90347  2.73739
3.18146  2.89801  2.37887  2.77519  2.98518  4.58477  3.61503
          0.02696  4.02262  4.74496  0.61958  0.77255  0.48576
0.95510
    260   3.27233  4.70095  4.50429  4.07784  3.10671  4.28578
4.67301  2.24410  3.82122  0.80077  3.04820  4.33725  4.63041
4.13251  3.98357  3.74018  3.54518  2.35032  5.10524  3.84990    277
- -
          2.68618  4.42225  2.77519  2.73123  3.46354  2.40513
3.72494  3.29354  2.67741  2.69355  4.24690  2.90347  2.73739
3.18146  2.89801  2.37887  2.77519  2.98518  4.58477  3.61503
          0.02696  4.02262  4.74496  0.61958  0.77255  0.48576
0.95510
    261   2.61931  4.54633  3.23926  2.65779  3.70503  3.50243
3.53697  2.99031  2.40398  2.36873  3.54747  3.02285  3.88169
2.90810  2.16001  2.73918  2.84371  2.84909  5.05691  3.37794    278
- -
          2.68618  4.42225  2.77519  2.73123  3.46354  2.40513
3.72494  3.29354  2.67741  2.69355  4.24690  2.90347  2.73739
3.18146  2.89801  2.37887  2.77519  2.98518  4.58477  3.61503
          0.02696  4.02262  4.74496  0.61958  0.77255  0.48576
0.95510
```

```
    262   3.18120  5.00542  3.68780  3.17038  4.39834  3.63746
3.94268  3.95237  2.31116  3.44111  4.43934  3.51762  4.13451
3.15727  0.77162  3.25933  3.43807  3.67032  5.42160  4.29519    279
- -
          2.68618  4.42225  2.77519  2.73123  3.46354  2.40513
3.72494  3.29354  2.67741  2.69355  4.24690  2.90347  2.73739
3.18146  2.89801  2.37887  2.77519  2.98518  4.58477  3.61503
          0.02696  4.02262  4.74496  0.61958  0.77255  0.48576
0.95510
    263   3.29187  4.63821  4.79138  4.22568  3.09141  4.52985
4.82525  2.04432  4.07634  0.91740  2.43919  4.51084  4.71744
4.21836  4.22245  3.87152  3.51388  2.20090  5.17483  4.10610    280
- -
          2.68618  4.42225  2.77519  2.73123  3.46354  2.40513
3.72494  3.29354  2.67741  2.69355  4.24690  2.90347  2.73739
3.18146  2.89801  2.37887  2.77519  2.98518  4.58477  3.61503
          0.02696  4.02262  4.74496  0.61958  0.77255  0.48576
0.95510
    264   3.08846  5.22327  2.56059  0.82931  4.58940  3.37593
3.97883  4.07348  2.87880  3.66752  4.63740  3.02771  3.98839
3.19292  3.30140  3.05703  3.39464  3.74275  5.73268  4.45792    281
- -
          2.68618  4.42225  2.77519  2.73123  3.46354  2.40513
3.72494  3.29354  2.67741  2.69355  4.24690  2.90347  2.73739
3.18146  2.89801  2.37887  2.77519  2.98518  4.58477  3.61503
          0.02696  4.02262  4.74496  0.61958  0.77255  0.48576
0.95510
    265   2.80481  4.90814  2.84095  2.58393  4.20435  3.36663
3.79318  3.78194  2.44355  3.33406  4.25921  3.05151  3.92100
1.30307  2.75219  2.70568  3.10767  3.44453  5.45529  4.10084    282
- -
          2.68618  4.42225  2.77519  2.73123  3.46354  2.40513
3.72494  3.29354  2.67741  2.69355  4.24690  2.90347  2.73739
3.18146  2.89801  2.37887  2.77519  2.98518  4.58477  3.61503
          0.02696  4.02262  4.74496  0.61958  0.77255  0.48576
0.95510
    266   3.32709  4.74617  4.42998  4.07280  3.19879  4.17148
4.68160  2.41344  3.83907  0.71022  3.17896  4.33366  4.58250
4.17133  3.98965  3.76330  3.61534  2.48932  5.13017  3.88739    283
- -
          2.68618  4.42225  2.77519  2.73123  3.46354  2.40513
3.72494  3.29354  2.67741  2.69355  4.24690  2.90347  2.73739
3.18146  2.89801  2.37887  2.77519  2.98518  4.58477  3.61503
          0.02696  4.02262  4.74496  0.61958  0.77255  0.48576
0.95510
    267   2.86005  4.34649  4.49739  3.99380  3.57624  4.15245
4.73709  1.16949  3.87585  2.17950  3.40747  4.22367  4.53929
4.16455  4.09964  3.52119  2.91290  1.56264  5.40821  4.20825    284
- -
          2.68618  4.42225  2.77519  2.73123  3.46354  2.40513
3.72494  3.29354  2.67741  2.69355  4.24690  2.90347  2.73739
3.18146  2.89801  2.37887  2.77519  2.98518  4.58477  3.61503
          0.02696  4.02262  4.74496  0.61958  0.77255  0.48576
0.95510
```

```
    268   2.83146  4.78038  3.21550  2.76886  3.70463  3.57504
3.80010  3.16401  2.48315  2.12033  3.74571  3.21923  4.01048
1.66607  2.78586  2.92778  3.07637  3.00244  5.16790  3.83071     285
- -
          2.68618  4.42225  2.77519  2.73123  3.46354  2.40513
3.72494  3.29354  2.67741  2.69355  4.24690  2.90347  2.73739
3.18146  2.89801  2.37887  2.77519  2.98518  4.58477  3.61503
          0.02696  4.02262  4.74496  0.61958  0.77255  0.48576
0.95510
    269   3.08846  5.22327  2.56059  0.82931  4.58940  3.37593
3.97883  4.07348  2.87880  3.66752  4.63740  3.02771  3.98839
3.19292  3.30140  3.05703  3.39464  3.74275  5.73268  4.45792     286
- -
          2.68618  4.42225  2.77519  2.73123  3.46354  2.40513
3.72494  3.29354  2.67741  2.69355  4.24690  2.90347  2.73739
3.18146  2.89801  2.37887  2.77519  2.98518  4.58477  3.61503
          0.02696  4.02262  4.74496  0.61958  0.77255  0.48576
0.95510
    270   1.55085  4.23153  3.73746  3.24773  3.67114  3.40471
4.14290  2.39577  3.17264  2.60408  3.60147  3.50892  3.96628
3.48422  3.47738  2.65550  2.85275  1.97193  5.19741  3.97243     287
- -
          2.68618  4.42225  2.77519  2.73123  3.46354  2.40513
3.72494  3.29354  2.67741  2.69355  4.24690  2.90347  2.73739
3.18146  2.89801  2.37887  2.77519  2.98518  4.58477  3.61503
          0.02696  4.02262  4.74496  0.61958  0.77255  0.48576
0.95510
    271   2.24469  4.23731  4.03625  3.52952  3.56784  3.70153
4.32220  2.00873  3.43578  2.41298  3.47829  3.77463  4.18410
3.73137  3.69946  2.89939  2.97675  1.33818  5.18732  3.97038     288
- -
          2.68618  4.42225  2.77519  2.73123  3.46354  2.40513
3.72494  3.29354  2.67741  2.69355  4.24690  2.90347  2.73739
3.18146  2.89801  2.37887  2.77519  2.98518  4.58477  3.61503
          0.02696  4.02262  4.74496  0.61958  0.77255  0.48576
0.95510
    272   2.48231  4.22334  3.91582  3.34580  3.51006  3.68006
4.19385  2.08932  3.29983  2.41710  3.43663  3.67517  4.00527
3.60112  3.57257  2.92913  2.79284  1.37164  5.08215  3.86815     289
- -
          2.68618  4.42225  2.77519  2.73123  3.46354  2.40513
3.72494  3.29354  2.67741  2.69355  4.24690  2.90347  2.73739
3.18146  2.89801  2.37887  2.77519  2.98518  4.58477  3.61503
          0.02696  4.02262  4.74496  0.61958  0.77255  0.48576
0.95510
    273   2.44282  4.63754  3.04415  2.52124  3.86840  3.41812
3.57171  3.26770  2.48504  2.90882  3.75181  2.61644  3.82813
2.80264  2.83043  2.64401  2.10288  2.90488  5.18654  3.64108     290
- -
          2.68618  4.42225  2.77519  2.73123  3.46354  2.40513
3.72494  3.29354  2.67741  2.69355  4.24690  2.90347  2.73739
3.18146  2.89801  2.37887  2.77519  2.98518  4.58477  3.61503
          0.02696  4.02262  4.74496  0.61958  0.77255  0.48576
0.95510
```

```
    274   2.47768  4.26661  3.68368  3.26884  3.70147  3.39575
4.19008  2.57430  3.19918  2.60064  3.64279  3.52412  2.54415
3.53191  3.48838  2.79428  2.89587  1.42435  5.25106  4.01948    291
- -
          2.68618  4.42225  2.77519  2.73123  3.46354  2.40513
3.72494  3.29354  2.67741  2.69355  4.24690  2.90347  2.73739
3.18146  2.89801  2.37887  2.77519  2.98518  4.58477  3.61503
          0.02696  4.02262  4.74496  0.61958  0.77255  0.48576
0.95510
    275   2.45764  4.39290  3.19605  3.01336  4.23553  3.09515
4.17653  3.72471  3.09626  3.41366  4.33007  3.29303  1.13183
3.46213  3.39743  2.43246  2.92003  3.26952  5.57580  4.30809    292
- -
          2.68618  4.42225  2.77519  2.73123  3.46354  2.40513
3.72494  3.29354  2.67741  2.69355  4.24690  2.90347  2.73739
3.18146  2.89801  2.37887  2.77519  2.98518  4.58477  3.61503
          0.02696  4.02262  4.74496  0.61958  0.77255  0.48576
0.95510
    276   2.85553  4.88047  3.34907  2.75275  3.65398  3.58218
3.66674  3.53521  2.05026  3.09774  4.00225  3.17792  3.97926
2.84628  1.37143  2.91125  3.06835  3.25956  5.25512  3.97585    293
- -
          2.68618  4.42225  2.77519  2.73123  3.46354  2.40513
3.72494  3.29354  2.67741  2.69355  4.24690  2.90347  2.73739
3.18146  2.89801  2.37887  2.77519  2.98518  4.58477  3.61503
          0.02696  4.02262  4.74496  0.61958  0.77255  0.48576
0.95510
    277   1.66647  4.66483  2.88952  2.35155  4.03639  3.25037
3.72273  3.43047  2.53981  3.06641  3.90576  2.99611  3.68259
2.90637  2.77073  2.63248  2.83076  3.07384  5.32864  4.02328    294
- -
          2.68618  4.42225  2.77519  2.73123  3.46354  2.40513
3.72494  3.29354  2.67741  2.69355  4.24690  2.90347  2.73739
3.18146  2.89801  2.37887  2.77519  2.98518  4.58477  3.61503
          0.02696  4.02262  4.74496  0.61958  0.77255  0.48576
0.95510
    278   3.30023  4.69967  4.63105  4.12815  3.08165  4.41382
4.72742  2.22560  3.87478  0.81607  2.77878  4.40788  4.67955
4.14218  4.03672  3.80712  3.54677  2.33254  5.13155  3.94796    295
- -
          2.68618  4.42225  2.77519  2.73123  3.46354  2.40513
3.72494  3.29354  2.67741  2.69355  4.24690  2.90347  2.73739
3.18146  2.89801  2.37887  2.77519  2.98518  4.58477  3.61503
          0.02696  4.02262  4.74496  0.61958  0.77255  0.48576
0.95510
    279   3.15037  4.51337  4.62848  4.19442  3.41414  4.33960
4.87364  0.98488  4.00374  1.90686  3.30757  4.42638  4.69774
4.32534  4.18959  3.77680  3.42957  1.80026  5.37820  4.11944    296
- -
          2.68618  4.42225  2.77519  2.73123  3.46354  2.40513
3.72494  3.29354  2.67741  2.69355  4.24690  2.90347  2.73739
3.18146  2.89801  2.37887  2.77519  2.98518  4.58477  3.61503
          0.02696  4.02262  4.74496  0.61958  0.77255  0.48576
0.95510
```

```
    280   0.81728  4.27297  3.62041  3.42136  4.20706  3.09296
4.42492  3.35544  3.44162  3.22989  4.24718  3.53130  3.84663
3.76944  3.68019  2.62296  2.91086  2.98372  5.61916  4.43687     297
- -
          2.68618  4.42225  2.77519  2.73123  3.46354  2.40513
3.72494  3.29354  2.67741  2.69355  4.24690  2.90347  2.73739
3.18146  2.89801  2.37887  2.77519  2.98518  4.58477  3.61503
          0.02696  4.02262  4.74496  0.61958  0.77255  0.48576
0.95510
    281   2.83784  5.14938  2.39402  1.21431  4.42273  3.31263
3.75755  3.63029  2.62337  3.45106  4.33168  2.83825  3.87010
2.92811  3.09199  2.80258  3.11399  3.48155  5.66276  4.25599     298
- -
          2.68618  4.42225  2.77519  2.73123  3.46354  2.40513
3.72494  3.29354  2.67741  2.69355  4.24690  2.90347  2.73739
3.18146  2.89801  2.37887  2.77519  2.98518  4.58477  3.61503
          0.02696  4.02262  4.74496  0.61958  0.77255  0.48576
0.95510
    282   1.62693  4.21717  3.57342  3.16405  3.97107  3.14071
4.15001  2.94255  3.10794  2.95559  3.89332  3.37761  3.80913
3.43769  3.41964  2.55373  1.67968  2.72263  5.41260  4.18991     299
- -
          2.68618  4.42225  2.77519  2.73123  3.46354  2.40513
3.72494  3.29354  2.67741  2.69355  4.24690  2.90347  2.73739
3.18146  2.89801  2.37887  2.77519  2.98518  4.58477  3.61503
          0.02696  4.02262  4.74496  0.61958  0.77255  0.48576
0.95510
    283   3.13093  4.43260  4.84360  4.33161  3.52670  4.50308
5.02657  1.19742  4.21665  1.94639  3.32808  4.55952  4.78391
4.46559  4.40848  3.88120  3.38775  1.32151  5.51011  4.31907     300
- -
          2.68618  4.42225  2.77519  2.73123  3.46354  2.40513
3.72494  3.29354  2.67741  2.69355  4.24690  2.90347  2.73739
3.18146  2.89801  2.37887  2.77519  2.98518  4.58477  3.61503
          0.02696  4.02262  4.74496  0.61958  0.77255  0.48576
0.95510
    284   2.88028  5.39059  1.41116  2.14734  4.68403  3.25209
3.74090  4.20651  2.69216  3.71082  4.54215  2.11239  3.84100
2.78889  3.13368  2.78970  3.15992  3.78715  5.86501  4.38792     301
- -
          2.68618  4.42225  2.77519  2.73123  3.46354  2.40513
3.72494  3.29354  2.67741  2.69355  4.24690  2.90347  2.73739
3.18146  2.89801  2.37887  2.77519  2.98518  4.58477  3.61503
          0.02696  4.02262  4.74496  0.61958  0.77255  0.48576
0.95510
    285   3.13256  4.47219  4.72508  4.18719  3.33129  4.42193
4.83840  1.80419  4.05871  1.34170  3.14647  4.43779  4.68899
4.27371  4.24055  3.77613  3.37796  1.37843  5.31020  4.15668     302
- -
          2.68618  4.42225  2.77519  2.73123  3.46354  2.40513
3.72494  3.29354  2.67741  2.69355  4.24690  2.90347  2.73739
3.18146  2.89801  2.37887  2.77519  2.98518  4.58477  3.61503
          0.02696  4.02262  4.74496  0.61958  0.77255  0.48576
0.95510
```

```
    286   2.81816  4.27820  4.28451  3.73875  3.35802  4.01329
4.11630  1.61129  3.59908  2.07979  3.28247  3.98848  4.36700
3.87697  3.81340  3.33624  3.09383  1.39571  5.06634  3.84742    303
- -
          2.68618  4.42225  2.77519  2.73123  3.46354  2.40513
3.72494  3.29354  2.67741  2.69355  4.24690  2.90347  2.73739
3.18146  2.89801  2.37887  2.77519  2.98518  4.58477  3.61503
          0.02696  4.02262  4.74496  0.61958  0.77255  0.48576
0.95510
    287   1.82648  4.27526  3.82842  3.20687  3.52934  3.69299
4.16482  1.96497  3.22501  2.39611  3.44247  3.62613  4.13158
3.53317  3.52654  3.01154  2.94965  1.82726  5.11234  3.89159    304
- -
          2.68618  4.42225  2.77519  2.73123  3.46354  2.40513
3.72494  3.29354  2.67741  2.69355  4.24690  2.90347  2.73739
3.18146  2.89801  2.37887  2.77519  2.98518  4.58477  3.61503
          0.02696  4.02262  4.74496  0.61958  0.77255  0.48576
0.95510
    288   2.65544  4.24907  3.71462  3.16401  2.56599  3.66044
2.91692  2.68732  3.07371  2.39578  3.34894  3.47686  4.03715
3.35278  3.35440  2.87491  2.88973  1.82789  4.53190  3.00296    305
- -
          2.68618  4.42225  2.77519  2.73123  3.46354  2.40513
3.72494  3.29354  2.67741  2.69355  4.24690  2.90347  2.73739
3.18146  2.89801  2.37887  2.77519  2.98518  4.58477  3.61503
          0.02696  4.02262  4.74496  0.61958  0.77255  0.48576
0.95510
    289   3.25456  4.57328  4.85608  4.29749  3.17290  4.54777
4.88716  1.60029  4.16106  1.08495  2.78370  4.55735  4.74477
4.30397  4.30145  3.89691  3.48224  2.00841  5.23783  4.15329    306
- -
          2.68618  4.42225  2.77519  2.73123  3.46354  2.40513
3.72494  3.29354  2.67741  2.69355  4.24690  2.90347  2.73739
3.18146  2.89801  2.37887  2.77519  2.98518  4.58477  3.61503
          0.02696  4.02262  4.74496  0.61958  0.77255  0.48576
0.95510
    290   2.14769  4.67000  2.91946  2.39197  3.97858  3.36235
3.67275  3.38221  2.49730  2.95987  3.83944  2.94878  3.68296
2.84569  2.84676  2.05316  2.79571  2.94149  5.27308  3.95879    307
- -
          2.68618  4.42225  2.77519  2.73123  3.46354  2.40513
3.72494  3.29354  2.67741  2.69355  4.24690  2.90347  2.73739
3.18146  2.89801  2.37887  2.77519  2.98518  4.58477  3.61503
          0.03776  4.02262  3.95540  0.61958  0.77255  0.48576
0.95510
    291   2.63203  4.59443  3.15542  2.86286  4.30710  1.21941
3.99595  3.79451  2.73096  3.40745  4.28526  3.23770  3.87450
3.22485  2.38775  2.74700  3.01623  3.37971  5.51251  4.29738    308
- -
          2.68618  4.42225  2.77519  2.73123  3.46354  2.40513
3.72494  3.29354  2.67741  2.69355  4.24690  2.90347  2.73739
3.18146  2.89801  2.37887  2.77519  2.98518  4.58477  3.61503
          0.02725  4.01210  4.73445  0.61958  0.77255  0.49680
0.93771
```

292 2.69943 4.18694 2.98841 2.58974 4.13495 3.46252
3.65123 3.52175 2.17646 3.11765 3.96114 3.05684 3.87869
2.81131 1.58709 2.74798 2.93479 3.20573 5.31867 4.04200 309
- -
2.68618 4.42225 2.77519 2.73123 3.46354 2.40513
3.72494 3.29354 2.67741 2.69355 4.24690 2.90347 2.73739
3.18146 2.89801 2.37887 2.77519 2.98518 4.58477 3.61503
0.02725 4.01210 4.73445 0.61958 0.77255 0.49680
0.93771
293 2.69186 4.66193 2.57981 2.76446 4.50056 0.93766
4.15944 4.02880 3.18335 3.67327 4.56516 3.18890 3.87860
3.42338 3.57978 2.79223 3.12291 3.55790 5.70063 4.49596 310
- -
2.68618 4.42225 2.77519 2.73123 3.46354 2.40513
3.72494 3.29354 2.67741 2.69355 4.24690 2.90347 2.73739
3.18146 2.89801 2.37887 2.77519 2.98518 4.58477 3.61503
0.02725 4.01210 4.73445 0.61958 0.77255 0.49680
0.93771
294 2.29314 4.76574 2.96375 2.44463 4.07066 3.16059
3.53132 3.48597 2.36647 3.08332 3.89937 2.87111 3.80034
2.74664 2.46049 2.22740 2.65330 3.15158 5.31642 3.99193 311
- -
2.68618 4.42225 2.77519 2.73123 3.46354 2.40513
3.72494 3.29354 2.67741 2.69355 4.24690 2.90347 2.73739
3.18146 2.89801 2.37887 2.77519 2.98518 4.58477 3.61503
0.02725 4.01210 4.73445 0.61958 0.77255 0.49680
0.93771
295 2.22740 4.87928 2.77385 2.26322 4.15195 3.03473
3.59853 3.40558 2.15002 3.15690 3.95556 2.86781 3.78404
2.65557 2.84587 2.47834 2.78540 3.23323 5.37104 4.01572 312
- -
2.68618 4.42225 2.77519 2.73123 3.46354 2.40513
3.72494 3.29354 2.67741 2.69355 4.24690 2.90347 2.73739
3.18146 2.89801 2.37887 2.77519 2.98518 4.58477 3.61503
0.02725 4.01210 4.73445 0.61958 0.77255 0.49680
0.93771
296 3.16824 4.99396 3.67365 3.15796 4.38416 3.62702
3.93346 3.93711 2.30475 3.42776 4.42593 3.50618 4.12428
3.14828 0.78476 3.24667 3.42563 3.65544 5.41150 4.28292 313
- -
2.68618 4.42225 2.77519 2.73123 3.46354 2.40513
3.72494 3.29354 2.67741 2.69355 4.24690 2.90347 2.73739
3.18146 2.89801 2.37887 2.77519 2.98518 4.58477 3.61503
0.02725 4.01210 4.73445 0.61958 0.77255 0.49680
0.93771
297 3.00805 5.09672 3.39132 2.77144 4.41115 3.64619
3.09454 3.86162 1.88483 3.33677 4.20847 3.18333 4.02352
2.77191 1.27164 3.01684 3.18490 3.55346 5.38025 4.16041 314
- -
2.68618 4.42225 2.77519 2.73123 3.46354 2.40513
3.72494 3.29354 2.67741 2.69355 4.24690 2.90347 2.73739
3.18146 2.89801 2.37887 2.77519 2.98518 4.58477 3.61503
0.02725 4.01210 4.73445 0.61958 0.77255 0.49680
0.93771

```
    298   3.15832  4.48794  4.76676  4.22344  3.28951  4.45781
4.85770  1.72097  4.09929  1.24176  3.09374  4.47424  4.70464
4.29050  4.26912  3.81040  3.39875  1.54891  5.29320  4.16131    315
- -
          2.68618  4.42225  2.77519  2.73123  3.46354  2.40513
3.72494  3.29354  2.67741  2.69355  4.24690  2.90347  2.73739
3.18146  2.89801  2.37887  2.77519  2.98518  4.58477  3.61503
          0.02725  4.01210  4.73445  0.61958  0.77255  0.49680
0.93771
    299   2.33062  4.82706  2.83201  2.08214  4.10647  3.30986
3.61530  3.52969  2.33881  3.11891  3.92599  2.90642  3.78815
2.76010  2.82132  2.34881  2.59223  3.03449  5.34670  4.00106    316
- -
          2.68618  4.42225  2.77519  2.73123  3.46354  2.40513
3.72494  3.29354  2.67741  2.69355  4.24690  2.90347  2.73739
3.18146  2.89801  2.37887  2.77519  2.98518  4.58477  3.61503
          0.02725  4.01210  4.73445  0.61958  0.77255  0.49680
0.93771
    300   2.56744  4.56526  2.72811  1.69333  4.22056  3.32138
3.66108  3.64268  2.46360  3.23086  4.04086  2.89452  3.79311
2.80925  2.88533  2.23894  2.82153  3.27884  5.44690  4.09285    317
- -
          2.68618  4.42225  2.77519  2.73123  3.46354  2.40513
3.72494  3.29354  2.67741  2.69355  4.24690  2.90347  2.73739
3.18146  2.89801  2.37887  2.77519  2.98518  4.58477  3.61503
          0.02725  4.01210  4.73445  0.61958  0.77255  0.49680
0.93771
    301   2.89597  4.45747  4.61216  4.06817  3.28025  4.32611
4.72271  1.63110  3.93444  1.22908  3.09722  4.32728  4.60676
4.15307  4.12047  3.66964  3.33046  1.86463  5.23068  4.09350    318
- -
          2.68618  4.42225  2.77519  2.73123  3.46354  2.40513
3.72494  3.29354  2.67741  2.69355  4.24690  2.90347  2.73739
3.18146  2.89801  2.37887  2.77519  2.98518  4.58477  3.61503
          0.02725  4.01210  4.73445  0.61958  0.77255  0.49680
0.93771
    302   1.60692  4.08977  3.70231  3.17295  3.58586  3.44647
4.05784  2.55608  3.08915  2.43982  3.37355  3.46903  3.96252
3.40100  3.40274  2.69933  2.83874  2.07662  5.09917  3.87438    319
- -
          2.68618  4.42225  2.77519  2.73123  3.46354  2.40513
3.72494  3.29354  2.67741  2.69355  4.24690  2.90347  2.73739
3.18146  2.89801  2.37887  2.77519  2.98518  4.58477  3.61503
          0.02725  4.01210  4.73445  0.61958  0.77255  0.49680
0.93771
    303   2.60930  4.56697  3.07003  2.36628  4.13053  3.45840
3.46971  3.53174  2.24819  3.07680  3.93445  2.99661  3.84940
2.68455  1.83925  2.67933  2.71349  3.21020  5.31568  4.01888    320
- -
          2.68618  4.42225  2.77519  2.73123  3.46354  2.40513
3.72494  3.29354  2.67741  2.69355  4.24690  2.90347  2.73739
3.18146  2.89801  2.37887  2.77519  2.98518  4.58477  3.61503
          0.02725  4.01210  4.73445  0.61958  0.77255  0.49680
0.93771
```

304 2.92117 4.36113 4.39868 3.90935 3.54452 3.94865
4.65673 1.83139 3.75891 2.13396 3.43251 4.14541 4.49280
4.08385 3.99003 3.45669 3.21006 1.03916 5.35728 4.10653 321
- -
2.68618 4.42225 2.77519 2.73123 3.46354 2.40513
3.72494 3.29354 2.67741 2.69355 4.24690 2.90347 2.73739
3.18146 2.89801 2.37887 2.77519 2.98518 4.58477 3.61503
0.02725 4.01210 4.73445 0.61958 0.77255 0.49680
0.93771
305 2.54831 4.82700 2.62904 2.07360 4.07934 3.38657
3.59920 3.46686 2.37738 3.09193 3.89761 2.88691 3.78590
2.63797 2.85623 2.60259 2.56162 2.75231 4.98250 3.97559 322
- -
2.68618 4.42225 2.77519 2.73123 3.46354 2.40513
3.72494 3.29354 2.67741 2.69355 4.24690 2.90347 2.73739
3.18146 2.89801 2.37887 2.77519 2.98518 4.58477 3.61503
0.02725 4.01210 4.73445 0.61958 0.77255 0.49680
0.93771
306 2.41045 4.63170 2.63838 2.40759 4.36308 1.37238
3.93873 3.76417 2.87192 3.42649 4.28533 2.99401 3.80327
3.15312 3.29290 2.64647 2.95609 3.33893 5.65329 4.33789 323
- -
2.68618 4.42225 2.77519 2.73123 3.46354 2.40513
3.72494 3.29354 2.67741 2.69355 4.24690 2.90347 2.73739
3.18146 2.89801 2.37887 2.77519 2.98518 4.58477 3.61503
0.02725 4.01210 4.73445 0.61958 0.77255 0.49680
0.93771
307 2.96299 4.17518 4.45175 3.88834 2.35301 4.11509
4.36229 2.19696 3.76295 1.21304 3.02174 4.10258 4.41764
3.93780 3.92514 3.37472 3.19740 2.14039 4.83985 3.58029 324
- -
2.68618 4.42225 2.77519 2.73123 3.46354 2.40513
3.72494 3.29354 2.67741 2.69355 4.24690 2.90347 2.73739
3.18146 2.89801 2.37887 2.77519 2.98518 4.58477 3.61503
0.02725 4.01210 4.73445 0.61958 0.77255 0.49680
0.93771
308 2.68693 5.09143 1.75460 2.20072 4.42333 2.50886
3.68435 3.88361 2.56211 3.22009 4.24305 2.76308 3.80512
2.79205 3.07710 2.55030 2.95399 3.49566 5.62813 4.21985 325
- -
2.68618 4.42225 2.77519 2.73123 3.46354 2.40513
3.72494 3.29354 2.67741 2.69355 4.24690 2.90347 2.73739
3.18146 2.89801 2.37887 2.77519 2.98518 4.58477 3.61503
0.02725 4.01210 4.73445 0.61958 0.77255 0.49680
0.93771
309 2.34099 4.66264 2.88918 2.40576 4.10811 2.50663
3.63715 3.52533 2.56424 3.14054 3.96564 2.97161 2.41066
2.90296 2.96147 2.50915 2.76803 3.16605 5.38635 4.06800 326
- -
2.68618 4.42225 2.77519 2.73123 3.46354 2.40513
3.72494 3.29354 2.67741 2.69355 4.24690 2.90347 2.73739
3.18146 2.89801 2.37887 2.77519 2.98518 4.58477 3.61503
0.04961 4.01210 3.49661 0.61958 0.77255 0.49680
0.93771

```
    310   2.36102  4.80351  2.42059  2.33492  4.07002  3.18131
3.61967  3.48994  2.43313  3.05376  3.90218  2.84846  3.78638
2.70429  2.87693  2.47325  2.78270  2.69914  5.32755  3.98421    327
- -
          2.68619  4.42226  2.77520  2.73124  3.46355  2.40514
3.72495  3.29353  2.67742  2.69356  4.24691  2.90346  2.73740
3.18147  2.89802  2.37883  2.77514  2.98519  4.58478  3.61504
          0.25865  2.49162  1.93011  0.42444  1.06171  0.48283
0.95980
    311   2.31146  4.31749  2.97015  2.73447  4.15138  1.43652
4.00422  3.50972  2.91124  3.14882  4.14732  3.10990  3.71169
3.26086  3.24337  2.50073  2.75351  3.08591  5.50170  4.22383    329
- -
          2.68618  4.42225  2.77519  2.73123  3.46354  2.40513
3.72494  3.29354  2.67741  2.69355  4.24690  2.90347  2.73739
3.18146  2.89801  2.37887  2.77519  2.98518  4.58477  3.61503
          0.03151  3.86913  4.59148  0.61958  0.77255  0.42151
1.06728
    312   2.57364  5.04672  1.82235  2.22788  4.38170  2.62524
3.65741  3.75755  2.49520  3.38350  4.18388  2.81563  3.79590
2.79775  2.79695  2.62500  2.90808  3.44870  5.57444  4.17848    330
- -
          2.68618  4.42225  2.77519  2.73123  3.46354  2.40513
3.72494  3.29354  2.67741  2.69355  4.24690  2.90347  2.73739
3.18146  2.89801  2.37887  2.77519  2.98518  4.58477  3.61503
          0.02725  4.01210  4.73445  0.61958  0.77255  0.49680
0.93771
    313   3.45411  4.88634  4.04680  3.80408  2.31065  3.99469
3.69188  3.44442  3.65767  2.86185  4.10530  3.89907  4.46609
3.92990  3.81200  3.57621  3.74192  3.32181  3.95916  0.76181    331
- -
          2.68618  4.42225  2.77519  2.73123  3.46354  2.40513
3.72494  3.29354  2.67741  2.69355  4.24690  2.90347  2.73739
3.18146  2.89801  2.37887  2.77519  2.98518  4.58477  3.61503
          0.02725  4.01210  4.73445  0.61958  0.77255  0.49680
0.93771
    314   2.54539  4.41780  3.14667  2.56794  3.86557  3.46891
3.41388  3.19452  2.38983  2.70087  3.74017  3.04890  3.85316
2.63289  2.04991  2.69947  2.82728  2.82621  5.16411  3.87982    332
- -
          2.68618  4.42225  2.77519  2.73123  3.46354  2.40513
3.72494  3.29354  2.67741  2.69355  4.24690  2.90347  2.73739
3.18146  2.89801  2.37887  2.77519  2.98518  4.58477  3.61503
          0.02725  4.01210  4.73445  0.61958  0.77255  0.49680
0.93771
    315   3.02679  4.40081  4.55957  4.00361  3.30585  4.27166
4.65450  1.70482  3.88521  1.32916  3.08723  4.26031  4.55788
4.10238  4.07758  3.60177  3.00859  1.71373  5.19973  4.05739    333
- -
          2.68583  4.42234  2.77496  2.73110  3.46362  2.40520
3.72503  3.29363  2.67748  2.69341  4.24698  2.90355  2.73748
3.18155  2.89809  2.37895  2.77526  2.98522  4.58457  3.61512
          0.15943  2.14495  3.49661  1.33167  0.30657  0.49680
0.93771
```

316 2.34970 4.55274 3.04497 2.55735 3.76448 3.42643
3.61977 2.95265 2.48601 2.79009 3.61785 3.01513 3.82602
2.85232 2.85729 2.38213 2.31373 2.85228 5.11189 3.82431 343
- -
2.68618 4.42225 2.77519 2.73123 3.46354 2.40513
3.72494 3.29354 2.67741 2.69355 4.24690 2.90347 2.73739
3.18146 2.89801 2.37887 2.77519 2.98518 4.58477 3.61503
0.02786 3.99036 4.71270 0.61958 0.77255 0.51889
0.90431
317 2.59744 4.88607 2.57081 2.25887 4.17774 3.32805
3.56625 3.60709 2.50743 3.12268 4.03001 2.79121 2.06211
2.80660 2.96702 2.60890 2.92273 3.26807 5.43205 4.07986 344
- -
2.68618 4.42225 2.77519 2.73123 3.46354 2.40513
3.72494 3.29354 2.67741 2.69355 4.24690 2.90347 2.73739
3.18146 2.89801 2.37887 2.77519 2.98518 4.58477 3.61503
0.02786 3.99036 4.71270 0.61958 0.77255 0.51889
0.90431
318 2.19635 4.14415 3.86122 3.25702 2.60098 3.68168
3.90697 2.33677 3.19440 1.95753 3.14728 3.57891 4.04962
3.45046 3.44378 2.96032 2.78593 2.18692 4.75675 3.54872 345
- -
2.68575 4.42247 2.77523 2.73127 3.46365 2.40537
3.72522 3.29359 2.67765 2.69319 4.24720 2.90374 2.73663
3.18162 2.89828 2.37863 2.77531 2.98531 4.58507 3.61534
0.54962 1.44386 1.67762 1.94058 0.15504 0.51889
0.90431
319 2.49891 4.56965 2.93284 2.37822 3.73849 3.37226
3.56998 3.17554 2.45826 2.69369 3.67018 2.89911 3.27082
2.78804 2.89062 2.53887 2.59081 2.79252 5.11408 3.75239 362
- -
2.68618 4.42225 2.77519 2.73123 3.46354 2.40513
3.72494 3.29354 2.67741 2.69355 4.24690 2.90347 2.73739
3.18146 2.89801 2.37887 2.77519 2.98518 4.58477 3.61503
0.03399 3.79454 4.51689 0.61958 0.77255 0.57247
0.83041
320 2.40819 4.61990 2.84649 2.45591 3.87961 3.35610
3.52647 3.15413 2.41679 2.86886 3.74937 2.95363 3.29667
2.80457 2.90010 2.27471 2.61650 2.94948 5.18841 3.87864 363
- -
2.68618 4.42225 2.77519 2.73123 3.46354 2.40513
3.72494 3.29354 2.67741 2.69355 4.24690 2.90347 2.73739
3.18146 2.89801 2.37887 2.77519 2.98518 4.58477 3.61503
0.03131 3.87522 4.59757 0.61958 0.77255 0.53928
0.87508
321 2.42806 4.39996 2.91668 2.49132 3.69182 3.36262
3.55313 3.16017 2.48815 2.72978 3.68891 2.99458 2.73278
2.76948 2.92551 2.54743 2.70226 2.91032 5.13572 3.83625 364
- -
2.68618 4.42225 2.77519 2.73123 3.46354 2.40513
3.72494 3.29354 2.67741 2.69355 4.24690 2.90347 2.73739
3.18146 2.89801 2.37887 2.77519 2.98518 4.58477 3.61503
0.02958 3.93123 4.65357 0.61958 0.77255 0.56525
0.83984

322 2.50204 4.83113 2.56246 2.27529 4.02429 3.29920
3.40230 3.52077 2.37028 2.95244 3.90340 2.82007 3.45261
2.70045 2.84241 2.40022 2.67851 3.05859 5.32542 3.97327 365
- -
2.68618 4.42225 2.77519 2.73123 3.46354 2.40513
3.72494 3.29354 2.67741 2.69355 4.24690 2.90347 2.73739
3.18146 2.89801 2.37887 2.77519 2.98518 4.58477 3.61503
0.02941 3.93704 4.65939 0.61958 0.77255 0.56218
0.84389
323 2.42200 4.84464 2.81784 2.31044 4.10941 3.36860
3.42892 3.48607 2.33381 3.04304 3.91262 2.80058 3.33540
2.54296 2.69050 2.38688 2.72518 3.16805 5.33077 3.97922 366
- -
2.68618 4.42225 2.77519 2.73123 3.46354 2.40513
3.72494 3.29354 2.67741 2.69355 4.24690 2.90347 2.73739
3.18146 2.89801 2.37887 2.77519 2.98518 4.58477 3.61503
0.02927 3.94149 4.66383 0.61958 0.77255 0.56521
0.83989
324 2.50573 4.04704 3.00849 2.34855 3.71393 3.40920
3.51212 3.22997 2.44776 2.70845 3.70753 2.90724 3.33775
2.60472 2.90409 2.53873 2.63017 2.90971 5.15230 3.84656 367
- -
2.68618 4.42225 2.77519 2.73123 3.46354 2.40513
3.72494 3.29354 2.67741 2.69355 4.24690 2.90347 2.73739
3.18146 2.89801 2.37887 2.77519 2.98518 4.58477 3.61503
0.02927 3.94149 4.66383 0.61958 0.77255 0.46967
0.98138
325 2.46268 4.57999 2.93946 2.54047 3.97693 1.76648
3.61895 3.17509 2.61743 3.03736 3.89491 3.03865 3.82355
2.91472 3.00783 2.61614 2.86895 3.06054 5.30317 4.00675 368
- -
2.68618 4.42225 2.77519 2.73123 3.46354 2.40513
3.72494 3.29354 2.67741 2.69355 4.24690 2.90347 2.73739
3.18146 2.89801 2.37887 2.77519 2.98518 4.58477 3.61503
0.02751 4.00262 4.72496 0.61958 0.77255 0.50656
0.92273
326 2.58692 5.07644 2.21762 1.83216 4.37458 3.32583
3.60725 3.83623 2.39971 3.23326 4.14673 2.70641 3.51166
2.68814 2.91184 2.62331 2.85239 3.29134 5.54030 4.13888 369
- -
2.68618 4.42225 2.77519 2.73123 3.46354 2.40513
3.72494 3.29354 2.67741 2.69355 4.24690 2.90347 2.73739
3.18146 2.89801 2.37887 2.77519 2.98518 4.58477 3.61503
0.05111 4.00262 3.45592 0.61958 0.77255 0.50656
0.92273
327 2.37933 3.93635 3.22921 2.65494 3.61623 3.44592
3.67431 2.99074 2.63068 2.33053 3.54546 3.06672 3.05226
2.89784 2.95098 2.51391 2.68766 2.73464 5.00789 3.74562 370
- -
2.68618 4.42225 2.77519 2.73123 3.46354 2.40513
3.72494 3.29354 2.67741 2.69355 4.24690 2.90347 2.73739
3.18146 2.89801 2.37887 2.77519 2.98518 4.58477 3.61503
0.04469 3.97967 3.68820 0.61958 0.77255 0.52940
0.88906

```
    328   2.46429  4.73309  2.97443  2.35760  3.97315  3.40215
3.42612  3.34299  2.31984  2.81497  3.81867  2.95028  2.76264
2.77513  2.73127  2.46323  2.80104  3.06899  5.24386  3.92645    371
- -
          2.68622  4.42229  2.77518  2.73122  3.46358  2.40513
3.72499  3.29358  2.67740  2.69348  4.24694  2.90344  2.73738
3.18151  2.89798  2.37879  2.77524  2.98523  4.58481  3.61507
          0.04644  3.09277        *  1.34909  0.30040  0.00000
*
//
```
